# *Tcf4* Deficiency Causes Recurrent Seizures in Mice

**DOI:** 10.1101/2024.11.01.621590

**Authors:** Laura Craciun, Vivianna R. DeNittis, Matthew T. Davis, Jeanne T. Paz, Kaoru Saijo

**Affiliations:** Department of Molecular and Cell Biology; Helen Wills Neuroscience Institute, University of California, Berkeley, CA 94720; Gladstone Institutes; Department of Neurology, University of California, San Francisco, CA 94158

**Keywords:** Transcription Factor 4 (TCF4), Pitt-Hopkins Syndrome (PTHS), Seizures, Epilepsy, Astrocytes

## Abstract

Transcription factor 4 (TCF4) is a transcription factor that is critical for the normal development and function of the central nervous system. Haploinsufficiency of *TCF4* causes Pitt-Hopkins Syndrome (PTHS), a lifelong neurodevelopmental disorder characterized by seizures and intellectual disability. To expand our currently limited understanding of TCF4 function and PTHS pathophysiology, we created a mouse model of PTHS with largely astrocyte-specific heterozygous knockout of *Tcf4*. These mice developed severe recurrent seizures and had decreased lifespans. In addition, we found that these mice had astrogliosis as well as increased neuronal activity in the cortex, hippocampus, amygdala, and hypothalamus. Furthermore, single nucleus RNA sequencing analysis revealed global changes in the gene expression profiles of excitatory neurons, inhibitory neurons, astrocytes, and oligodendrocytes of PTHS compared to wild-type mice. Overall, this is the first report of a PTHS mouse model with seizures, providing the field with a tool to investigate the mechanisms of PTHS development and progression and develop therapeutics for PTHS and its associated epilepsy.

**HIGHLIGHTS:**

- Novel PTHS mouse model that targets astrocytes develops epilepsy
- PTHS mice exhibit astrogliosis and increased neuron activity
- Changes in gene expression profile observed in PTHS mice across neurons and glial cells

## 1. INTRODUCTION

Transcription factor 4 (TCF4) is a basic helix-loop-helix (bHLH) transcription factor (TF) that binds to Ephrussi box (E-box) DNA motifs either as a homodimer or as a heterodimer with other TFs (Bergqvist et al., 2000; Dennis et al., 2019). Autosomal dominant *de novo* mutations that result in haploinsufficiency of *TCF4* cause a rare but severe neurodevelopmental disorder called Pitt-Hopkins Syndrome (PTHS) (PITT and HOPKINS, 1978; Pontual et al., 2009; Sweatt, 2013). Symptoms of PTHS start to present at early ages and include recurrent seizures, autistic behaviors, intellectual disability, motor delay, absent or limited speech, and craniofacial dysmorphism (Amiel et al., 2007; Pontual et al., 2009; Sweatt, 2013; Whalen et al., 2012). Apart from PTHS, studies have linked other variants of *TCF4* to schizophrenia, post-traumatic stress disorder, depression, and bipolar disorder (Blake et al., 2010; Lennertz et al., 2011; Mossakowska-Wójcik et al., 2018; Program et al., 2019; Quednow et al., 2011; Teixeira et al., 2021). Therefore, there is great interest in understanding the disease-causing mechanisms of TCF4 and developing relevant therapeutic interventions.

In both mouse and human brains, TCF4 is expressed in most major cell types, including excitatory neurons, inhibitory neurons, astrocytes, and oligodendrocytes (Chen et al., 2021; Kim et al., 2020; Schoof et al., 2020; Thaxton et al., 2018). It is also critical for brain development, with its expression levels peaking before birth during neurogenesis, then declining to lower steady-state levels in adulthood (Kim et al., 2020; Page et al., 2018; Rannals et al., 2016). Indeed, TCF4 has been shown to regulate neural progenitor cell proliferation and migration in several brain regions—such as the cortex and hippocampus (Chen et al., 2016; Jung et al., 2018; Li et al., 2019; Page et al., 2018)—and it is involved in modulating neurite length, synapse development, synaptic transmission, and synaptic plasticity (Crux et al., 2018; Kennedy et al., 2016; Schoof et al., 2020; Thaxton et al., 2018). Outside of neurons, TCF4 is required for oligodendrocyte differentiation and normal myelination (Phan et al., 2020; Wedel et al., 2020).

Over the years, both mouse and organoid models of PTHS have been created and have identified a variety of phenotypes related to TCF4 dysfunction. Most of these existing models are heterozygous knockouts of *Tcf4* in neurons, as conventional homozygous deletion results in embryonic lethality (Bergqvist et al., 2000; Jung et al., 2018). These PTHS mouse models replicate behavioral symptoms observed in humans, such as hyperactivity, social interaction deficits, poor vocalizations, reduced anxiety, and impaired learning and memory (Kennedy et al., 2016; Tamberg et al., 2020; Thaxton et al., 2018). However, seizures have yet to be observed in any animal model of PTHS, hampering the development of treatments for patients.

To our knowledge, there is currently no published work on TCF4 function in astrocytes, which are glial cells that maintain brain homeostasis by modulating gliosis, synaptic structure and function, excitatory and inhibitory neurotransmission, and ion and water concentrations (Allen and Eroglu, 2017; Bernardinelli et al., 2014; Clarke and Barres, 2013; Durkee and Araque, 2019; Molofsky and Deneen, 2015). However, since several studies have determined that TCF4 is highly expressed in astrocytes throughout the brain (Chen et al., 2021; Kim et al., 2020; Schoof et al., 2020), and there is evidence that astrocytes play important roles in neurodevelopmental disorders that are similar to PTHS (Devinsky et al., 2013; Patel et al., 2019), we hypothesized that TCF4 could have important astrocytic functions that may contribute to PTHS pathology. To test our hypothesis, we generated a novel mouse strain to model PTHS that has primarily astrocytic heterozygous knockout of *Tcf4*. Strikingly, these mice developed recurrent seizures that happened spontaneously and could be provoked with handling. They also had decreased lifespans, showed evidence of increased neuronal activity and astrogliosis, and exhibited differential gene expression in many cell types. Importantly, through our examination of this model, we have identified potential molecular candidates that could be causing the PTHS seizure phenotype.

## 2. RESULTS

### 2.1. Generation and initial characterization of a novel PTHS mouse model

TCF4 is highly expressed in astrocytes (Kim et al., 2020), so we sought to investigate whether its selective deletion in these cells could recapitulate some of the phenotypes of PTHS. For these studies, we generated a heterozygous *Tcf4* knock-out mouse (*Tcf4^fl/+^*;*Aldh1l1-Cre*, hereafter referred to as Cre+) by crossing mice containing a floxed conditional allele of *Tcf4* with mice constitutively expressing *Cre* recombinase under the control of the *Aldh1l1* promoter (Figure 1A). In this model, *Aldh1l1*-Cre mediates the excision of exon 4 from the floxed allele of *Tcf4*, resulting in a frameshift mutation that prevents the expression of the long isoform of *Tcf4* (Figure 1B) (Skarnes et al., 2011). Such deletion occurs primarily in astrocytes, although some populations of neurons and oligodendrocytes may also be affected (Tien et al., 2012). Reduced expression of *Tcf4* in astrocytes was confirmed in this model, with levels of *Tcf4* being diminished by approximately half in Cre+ mice compared to littermate controls containing a floxed allele but no Cre (*Tcf4^fl/+^*, hereafter referred to as Cre-) (Figure 1C).

**Figure 1.**
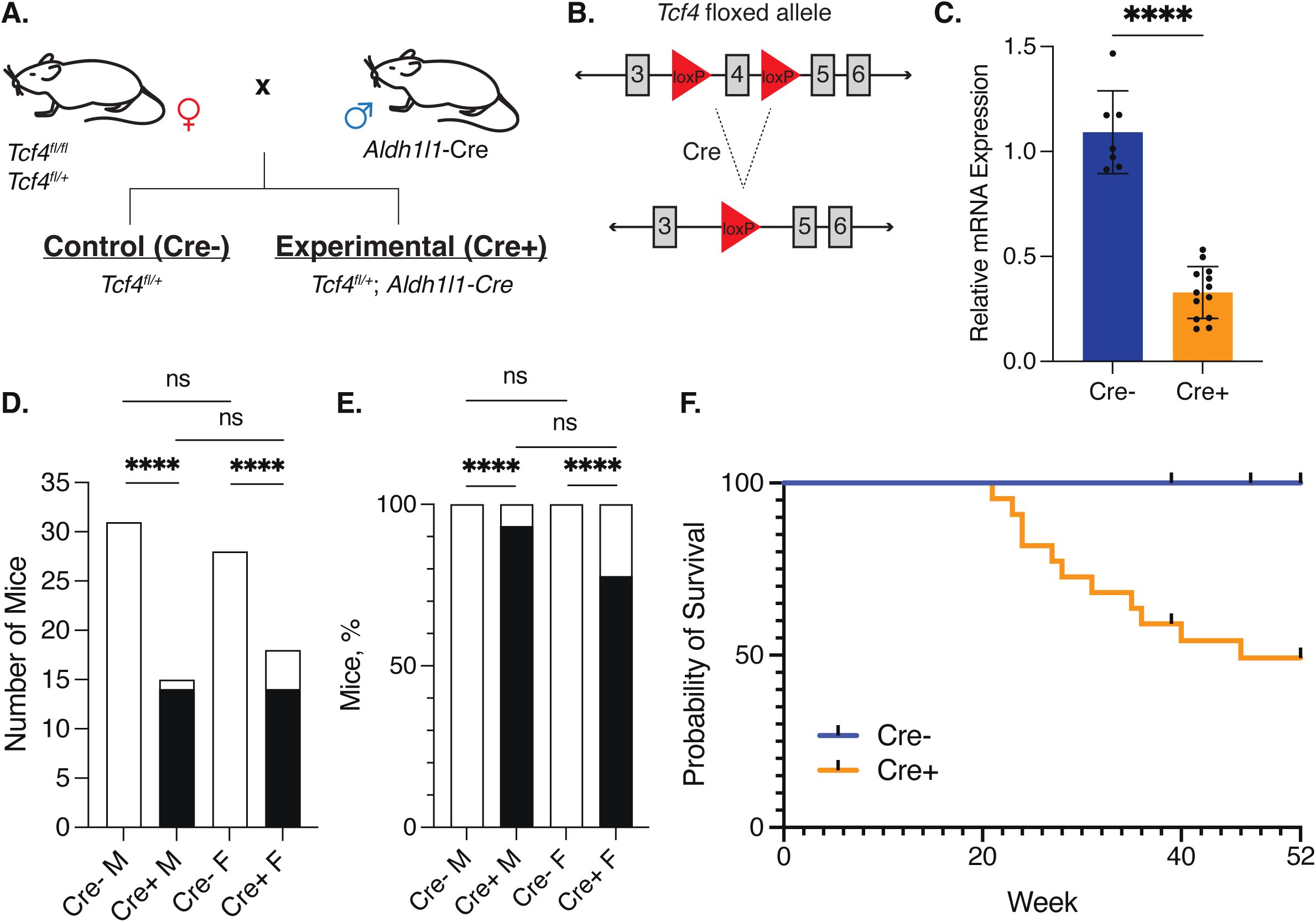
Generation of the PTHS mouse model and characterization of its major phenotypes. **(A)** Schematic of the breeding strategy used to generate our PTHS mouse in which *Aldh1l1*-expressing cells are heterozygous for *Tcf4* expression. **(B)** Schematic showing the floxed allele of *Tcf4* and the deletion of exon 4 following Cre-mediated excision. **(C)** Relative *Tcf4* mRNA levels in astrocytes of Cre- (blue) and Cre+ (orange) mice, as determined by RT-qPCR. \*\*\*\**p*<0.0001, Student’s *t*-test. **(D)** Stacked bar chart showing the numbers of male and female Cre+ mice, as well as their littermate controls, that did (black) and did not (white) have seizures. \*\*\*\**p*<0.0001, Fisher’s exact test. **(E)** Stacked bar chart showing the percentages of male and female Cre+ mice, as well as their littermate controls, that did (black) and did not (white) have seizures. \*\*\*\**p*<0.0001, Fisher’s exact test. **(F)** The survival of Cre- (blue line; *n*=22, grouped males and females) and Cre+ (orange line; *n*=22, grouped males and females) mice was tracked over the course of a year. Chi square=10.66, df=1, *p*=0.0011 (Mantel-Cox test).

As the mice aged (past three months of age), we found that a substantial proportion of male and female *Cre+* mice exhibited recurrent seizure-like behaviors. During our weekly monitoring of these mice in their home cages, we noticed abnormal behaviors that were characterized by stiff muscles followed by a convulsion and falling over for 1–2 minutes, with subsequent wild running and jumping in the cage (examples shown in Supplemental Videos 1 and 2). This phenomenon was observed both with and without handling of the mice when doing routine checks or cleaning their cages. We also observed that these seizures were recurrent in the mice, which would constitute an epilepsy diagnosis in human patients (Fisher et al., 2014). To accurately determine how many mice experienced these seizures, the mice were monitored weekly for a year. The earliest seizures were observed at three months of age and continued to appear as the mice got older. By the end of the year-long monitoring, we found that 14 of the 15 Cre+ male mice exhibited seizures (93.3%) (Figure 1D and E). Of the 18 Cre+ female mice we monitored, 14 had seizures (77.8%) and 4 did not exhibit any seizure activity during the weekly checks (Figure 1D and E). Notably, we did not detect seizures in any of the male or female Cre- mice (Figure 1D and E). Importantly, there were no significant differences in the numbers of Cre+ male and female mice that had seizures (Figure 1D and E); therefore, we grouped males and females together in our subsequent experiments.

At 10–12 months of age, we found that male Cre+ mice were significantly smaller than their Cre- counterparts. A similar difference was not observed between Cre+ and Cre- female mice; however, Cre+ females did trend on the smaller side (Supplemental Figure 1A). Moreover, female Cre+ mouse brains were significantly lighter than their Cre- controls, whereas there was no difference in brain weight between male Cre+ and Cre- controls (Supplemental Figure 1B). While we did not observe sex dimorphism in the seizure phenotype, it is interesting that there were sex-dependent differences in the body and brain weights. However, it is unclear whether these differences correlate to any specific PTHS phenotypes.

Because severe seizures can cause death (Giussani et al., 2023), we closely monitored a cohort of 44 mice (22 Cre- and 22 Cre+ mice) with weekly checks over the course of a year, starting at each mouse’s date of birth (P0). Almost half of the Cre+ mice were found dead in their cages from unknown causes starting at 20 weeks old, while no such premature death was detected in control mice (Figure 1F). Overall, these observations show that our novel mouse model exhibits seizures and a decreased lifespan. Therefore, these mice will be useful for studying not only PTHS-associated seizures, but epilepsy in general.

### 2.2. Characterization of the behavioral seizures of PTHS mice

As shown in Figure 1D and E, we noticed that Cre+ but not Cre- mice exhibited behavioral seizures. To conduct a precise analysis of these seizures, we video recorded mice after placing them in a new cage. The mice were recorded for 10 minutes, which was enough time to capture the rapid seizure onset immediately after being placed in the new cage and the recovery that occurred after the seizure episode. We focused our analysis of the videos on the seizure only, and scored the behavior of the mice using the state-of-the-art Racine scale (scoring criteria is shown in Figure 2A) (Racine, 1972) to quantify the severity of the seizures as well as to determine the duration and latency of each Racine stage. For these analyses, Racine scores of 3–4 were considered moderate seizure behavior and scores of 5–6 were considered severe seizure behavior. We never observed death in these mice at the end of their seizures when video recording. Out of the 19 Cre+ mice that we recorded (7 females and 12 males), 17 (89.5%) had behavioral seizure scores of 4 or higher on the Racine scale, with 47.4% (9 out of 19 mice) experiencing severe seizures (Figure 2B–E). In contrast, the 4 female and 4 male control (Cre-) mice did not have any seizures, as defined by their Racine score (2 or less; Figure 2B–E). The controls that scored a 1 or 2 on the Racine scale showed freezing behavior when they were placed in the new cage, which was followed by exploration of the cage.

**Figure 2.**
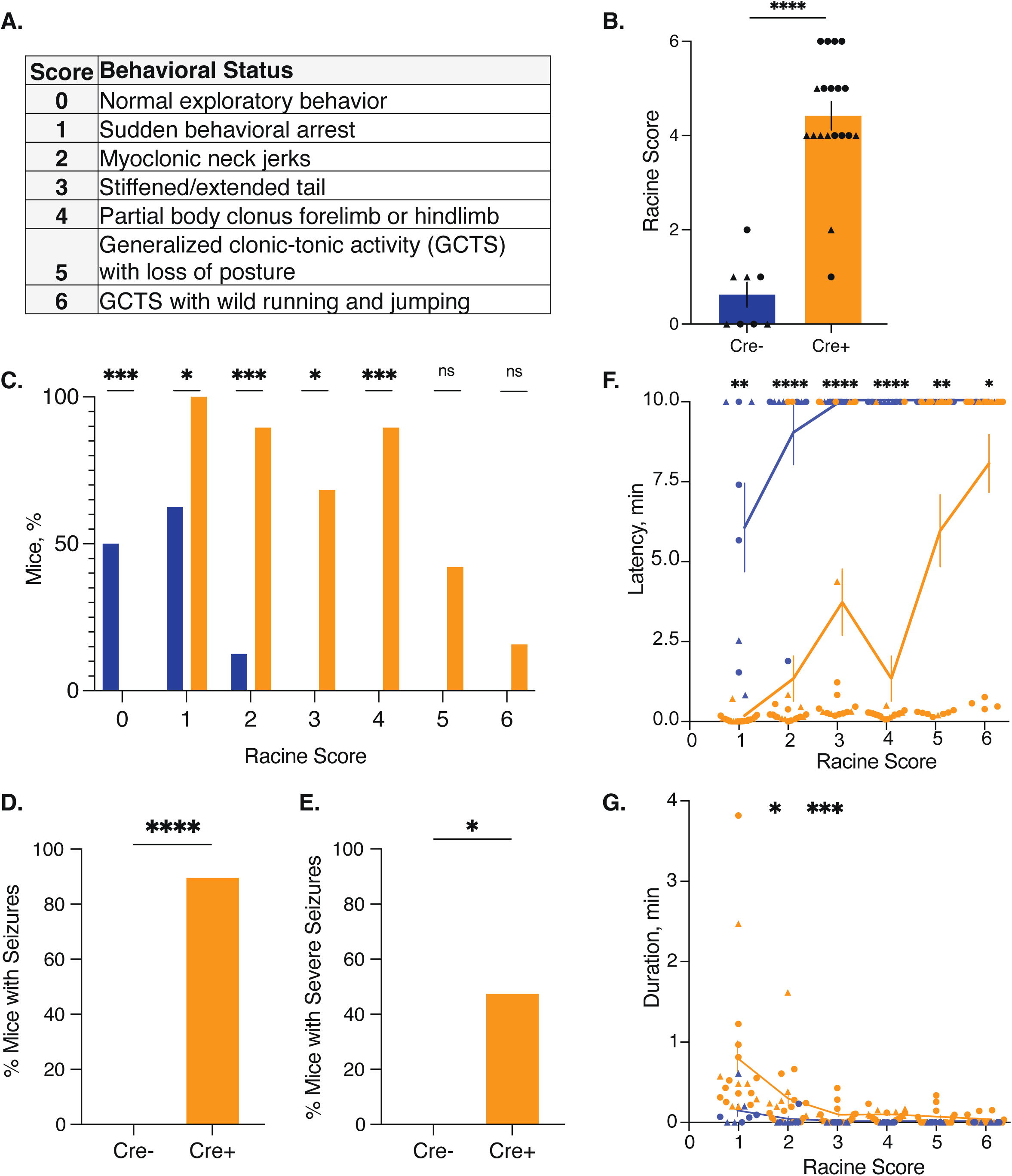
Characterization of the seizure phenotype observed in our PTHS mouse model. **(A)** Table of the modified Racine score criteria. **(B)** Maximum Racine score recorded for each Cre- (blue, *n*=8) and Cre+ (orange, *n*=19) mouse. \*\*\*\**p*<0.0001, Mann-Whitney. **(C)** Percentages of Cre- (blue, *n*=8) and Cre+ (orange, *n*=19) mice that had the modified Racine scores indicated. ns=not significant, \**p*<0.05 and \*\*\**p*<0.001, Fisher’s exact test. **(D)** Percentages of Cre- and Cre+ mice that had Racine scores ≥ 3. \*\*\*\**p*<0.0001, Fisher’s exact test. **(E)** Percentages of Cre- and Cre+ mice that had severe Racine scores (≥5). \**p*<0.05, Fisher’s exact test. **(F)** Latency to the onset of Racine scores is shown. Blue indicates Cre- (*n*=8) and orange indicates Cre+ (*n*=19). \**p*<0.05, \*\**p*<0.01, \*\*\*\**p*<0.0001, 2-way ANOVA. **(G)** Amount of time at the indicated modified Racine score is shown. Blue indicates Cre- (*n*=8) and orange indicates Cre+ (*n*=19). \**p*<0.05, \*\*\**p*<0.001, 2-way ANOVA. **(B-G)** Where applicable: means ± SEM are represented, circles denote males, triangles denote females, blue denotes Cre-, and orange denotes Cre+.

We next analyzed the latency to onset of each Racine score from the beginning of the video (when the mouse is placed in the cage) and found that for each score, Cre+ mice had significantly shorter latencies than Cre- mice (Figure 2F). The amount of time each mouse spent at a specific score number was also calculated and was found to be significantly longer for Cre+ mice at scores of 2 and 3 (Figure 2G). Overall, our data suggest that our PTHS mice have moderate to severe seizures, a symptom that is characteristic of human PTHS.

The genetic background of different mouse strains can have profound impacts on measures of seizure susceptibility (Schauwecker, 2011). The *Aldh1l1*-*Cre* mice used for our breeding scheme were on the C57BL/6J background, while the *Tcf4^fl/fl^* mice were on the C57BL/6N background. Though both of these substrains originated from C57BL/6 mice, they now differ genetically as a result of accumulated mutations over time. Therefore, we sought to exclude the possibility of any substrain-dependent effects on our seizure phenotype. When we genotyped our mice to determine if they were of the J or N background, we found that substrain did not correlate with seizure incidence (Supplemental Figure 2), suggesting that perturbing the function of *Tcf4* in astrocytes is indeed detrimental to the health of Cre+ mice and causes the development of seizures, independent of genetic background.

### 2.3. Neuronal activity and astrogliosis are increased in PTHS mice

During seizures, neurons exhibit a surge of excess electrical activity. Expression of the proto-oncogene c-Fos in neurons has been shown to be transiently induced in the brain in response to neuronal activity (Dragunow and Faull, 1989; Madabhushi et al., 2015; Zhang et al., 2002) as well as after seizures(Morgan et al., 1987). Therefore, to identify areas with increased neuronal activity, we performed unbiased immunostaining analysis for the expression of c-Fos in brain sections from Cre+ mice and their Cre-littermate controls. To capture such expression, PTHS mice were sacrificed within 30 minutes of having a seizure, as standard for the time course of c-Fos expression^41-44^. Brains were immediately processed for coronal sectioning (the frontal plane), and then the sections were subjected to immunostaining. Our results showed a higher number of c-Fos cells in the cortex, hippocampus, amygdala, and hypothalamus of Cre+ mice compared to Cre- mice (Supplemental Figure 3A). To quantify the numbers of c-Fos+ cells in these areas, we selected images of corresponding coronal sections from each mouse brain and digitally magnified them to focus on the areas indicated in the schematic shown in Supplemental Figure 3B. From these magnified images, we counted the numbers of c-Fos+ cells and confirmed that there were significantly more c-Fos+ cells in these areas in Cre+ mouse brains than in Cre-mouse brains (Figure 3). These results indicate that Cre+ mice had increased neuronal activity in their cortex, hippocampus, amygdala, and hypothalamus in response to seizures.

**Figure 3.**
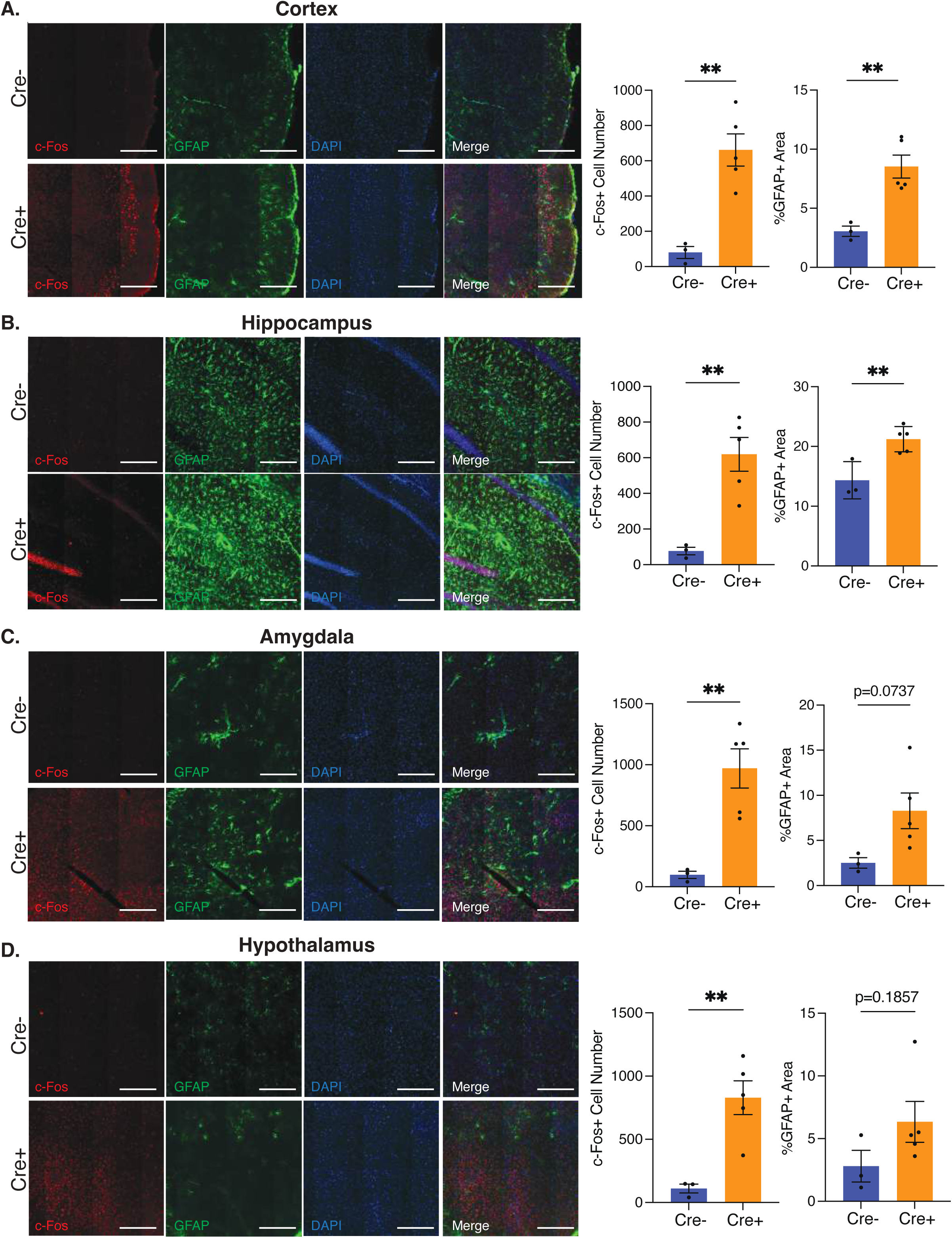
Immunostaining of brain regions to identify c-Fos+ cells and astrogliosis in PTHS mice. Immunostaining was performed on brain sections from Cre+ and Cre- mice. Representative images from the cortex **(A),** hippocampus **(B),** amygdala **(C),** and hypothalamus **(D)** are shown. Sections were immunostained for c-Fos (red) and GFAP (green). DAPI (blue) was used to indicate nuclei. Each scale bar indicates 125 microns. Images were quantified by determining the number of c-Fos+ cells and the percent of GFAP+ area in each region. The results of the quantification performed for each brain region (shown as means ± SEM) of Cre- (blue bars) and Cre+ (orange bars) mice are to the right of the corresponding images. \*\**p*<0.01, unpaired *t*-test. All scale bars indicate

Reactive astrogliosis is the process by which a range of genetic, epigenetic, molecular, morphological, and/or functional changes occur in astrocytes in response to injury or pathology (Liddelow and Barres, 2017; Pekny et al., 2016; Pekny and Pekna, 2014). In both human and animal models of diseases with acquired epileptic seizures, astrogliosis is a hallmark of seizure foci, the neurons in a brain region involved in seizure initiation, spread, or maintenance (Coulter and Steinhäuser, 2015; Devinsky et al., 2013; Patel et al., 2019). A defining feature of reactive astrocytes is their increased expression of intermediate filament proteins, such as glial fibrillary acidic protein (GFAP), which is often used as a marker of reactive astrogliosis (Buffo et al., 2008). To assess astrogliosis in our mouse model, we performed immunostaining for GFAP expression on our c-Fos-stained coronal brain sections of post-seizure Cre+ mice and Cre- controls with no seizures. These studies showed increased staining for GFAP in different brain areas in Cre+ compared to Cre- mice (Supplemental Figure 3A). To evaluate the degree of astrogliosis in the brain regions we identified as having increased neuronal activity in our PTHS mouse model compared to Cre-littermate controls, we examined digitally magnified images of our four regions of interest (Supplemental Figure 3B) and found that GFAP+ cells appeared to develop various degrees of hypertrophy in their cell body and processes, as would be expected of reactive astrocytes. To quantify the GFAP signal, as a measure of astrogliosis, we determined the percentage of GFAP-positive area in our images in each of the identified brain areas. In the cortex and the hippocampus of Cre+ mice, we saw significantly more astrogliosis than in Cre- mice (Figure 3A and B). In the amygdala and the hypothalamus, we did not observe significant differences in astrogliosis between Cre+ mice and controls; however, the amygdala does seem to trend toward more astrogliosis in Cre+ mice (Figure 3C and D). Overall, our data demonstrate that PTHS mice exhibit increased neuronal activity in specific brain regions that, in some cases, is also accompanied by astrogliosis.

### 2.4. Characterization of brain cells in PTHS mice by single nucleus RNA-sequencing

Next, we aimed to investigate the molecular mechanisms underpinning the seizure phenotype in our PTHS mouse model. Because TCF4 is a transcription factor, we expected that transcriptional changes might explain the phenotypes observed in these mice. To test our hypothesis, we used single nucleus RNA-sequencing (snRNA-seq) to identify differentially expressed genes (DEGs) in the brain cells of Cre+ versus Cre- mice. Our experiment included 6 Cre+ females, 6 Cre-females, 6 Cre+ males, and 5 Cre- males (Supplemental Table 1) that were sacrificed at 10 months of age. Whole brains were obtained, and nuclei were isolated and run on the Evercode™ Whole Transcriptome platform, which is based on SPLiT-seq (Rosenberg et al., 2018). The RNA obtained was of good quality (data not shown), with a total of 59,306 cells and 57,010 features recovered (7.4% of the cells were discarded).

With canonical component analysis (CCA) and k-means clustering, we detected a total of 24 clusters (Figure 4A). These clusters were based on the gene expression data of the top five markers, as shown in the heat map in Figure 4B. Using marker annotation approaches, we further identified the major brain cell types of the clusters, which included many neuron and glial cell types (Figure 4C). The only clusters that showed differences in cell numbers between Cre- and Cre+ mice were the excitatory neurons 2 and oligodendrocyte clusters, which had greater cell numbers in Cre+ mice compared to Cre- (Supplemental Figure 4A).

**Figure 4.**
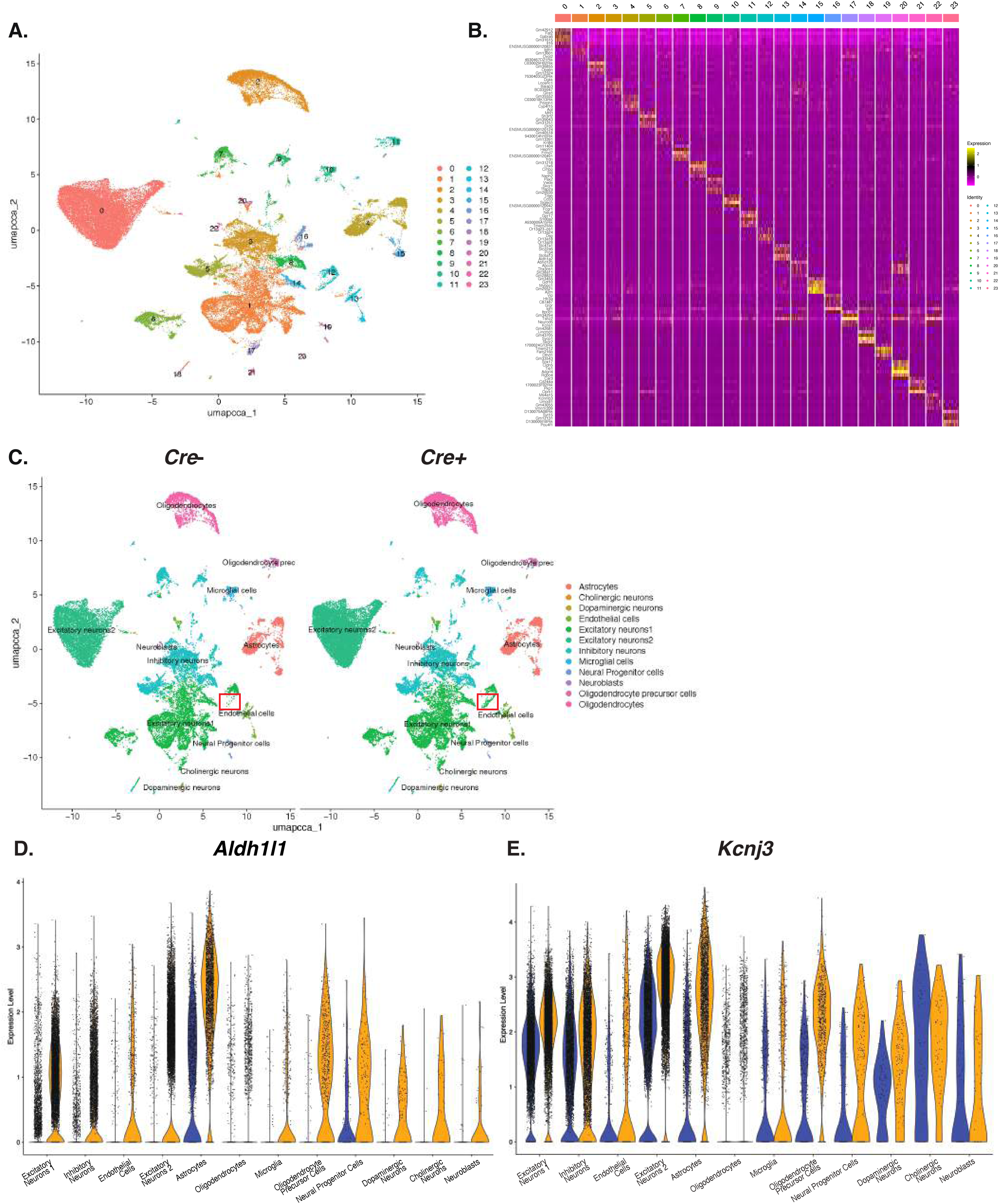
Characterization of brain cells by snRNA-seq. **(A)** A UMAP plot of all the cells from the Cre- and Cre+ males and females is shown. A total of 24 clusters were identified. **(B)** Heatmap showing the top five differentially expressed genes in Cre+ compared to Cre- cells in each cluster. This map was also used to generate the UMAP plot shown in A. Each row indicates a gene, and each column belongs to a cell within a cluster. The cluster IDs are color-coded at the top of the heatmap. **(C)** UMAP plots showing the cell types of the clusters shown in A for both Cre- (left) and Cre+ (right) mice. Combined data from males and females is shown. **(D and E)** Violin plots of *Aldh1l1* (left) and *Kcnj3* (right) expression in different cell types. These genes were the most significantly differentially expressed between Cre- (blue) and Cre+ (orange) cells. The width of each plot represents the number of cells at a given expression level.

No differences in clusters were observed between males and females (Supplemental Figure 4B), which is consistent with our observation that the seizure phenotype was not sex-dependent (Figure 1D and 1E). Therefore, when we looked for changes in gene expression in Cre+ versus Cre- brains, we combined male and female results for each genotype. In this analysis, *Aldh1l1* (aldehyde dehydrogenase 1 family member L1) was one of the most significantly upregulated genes in Cre+ mice compared to Cre- mice across all of the identified cell types (Figure 4D; Supplemental Figure 6A). This is likely because the *Aldh1l1*-Cre mouse line used to establish our PTHS mouse model is a BAC-transgenic mouse that ectopically overexpresses this gene (Tien et al., 2012). Interestingly, we also found that the epilepsy-associated gene *Kcnj3* (potassium inwardly rectifying channel subfamily J member 3) was also significantly upregulated in Cre+ mice compared to Cre- mice in most of the cell types identified (Figure 4E; Supplemental Figure 6B).

### 2.5. Identification of differentially expressed genes (DEGs) in neuron clusters

Seizures are caused by aberrant firing of neuronal populations due to an imbalance of excitatory and inhibitory currents (Coulter and Steinhäuser, 2015); therefore, probing neuron populations could reveal interesting candidate genes to help explain the seizure phenotype in PTHS. When we compared the differences in gene expression in the cell-type specific clusters of Cre+ and Cre- mice, we first focused on the excitatory neurons 1 (EN1) population, since the UMAP plot indicated a segregated cluster within this population that was only present in Cre+ cells (Figure 4C, red box). A comparison of the genes expressed in the Cre+ and Cre- EN1 clusters showed a high number of upregulated genes with some down regulated genes (Figure 5A). Gene ontology (GO) enrichment (Supplemental Figure 7A) and network/pathway analysis (Figure 5B) of the EN1 DEGs indicated that these genes are involved in cellular development, energy metabolism, and learning and memory. A number of the genes that were differentially expressed in Cre+ EN1 cells reached significance (upregulated: log2fold change > 0.5 and Bonferroni adjusted *p-*value (p adj) < 0.05; downregulated: log2fold change < −0.5 and p adj < 0.05) and are listed in Supplemental Table 4. Interestingly, a few of the most significant DEGs in the Cre+ EN1 cluster included *Kcnc4*, *Pcdhgc5*, *Slc2a1*, *Pik3r6*, *Arc*, and *Slc41a3* (Figure 5C), which are linked to epilepsy (Baaij, 2015; Dell’Isola et al., 2022; Egbenya et al., 2023; Gozzelino et al., 2022; Larsen et al., 2015; Otalora et al., 2011).

**Figure 5.**
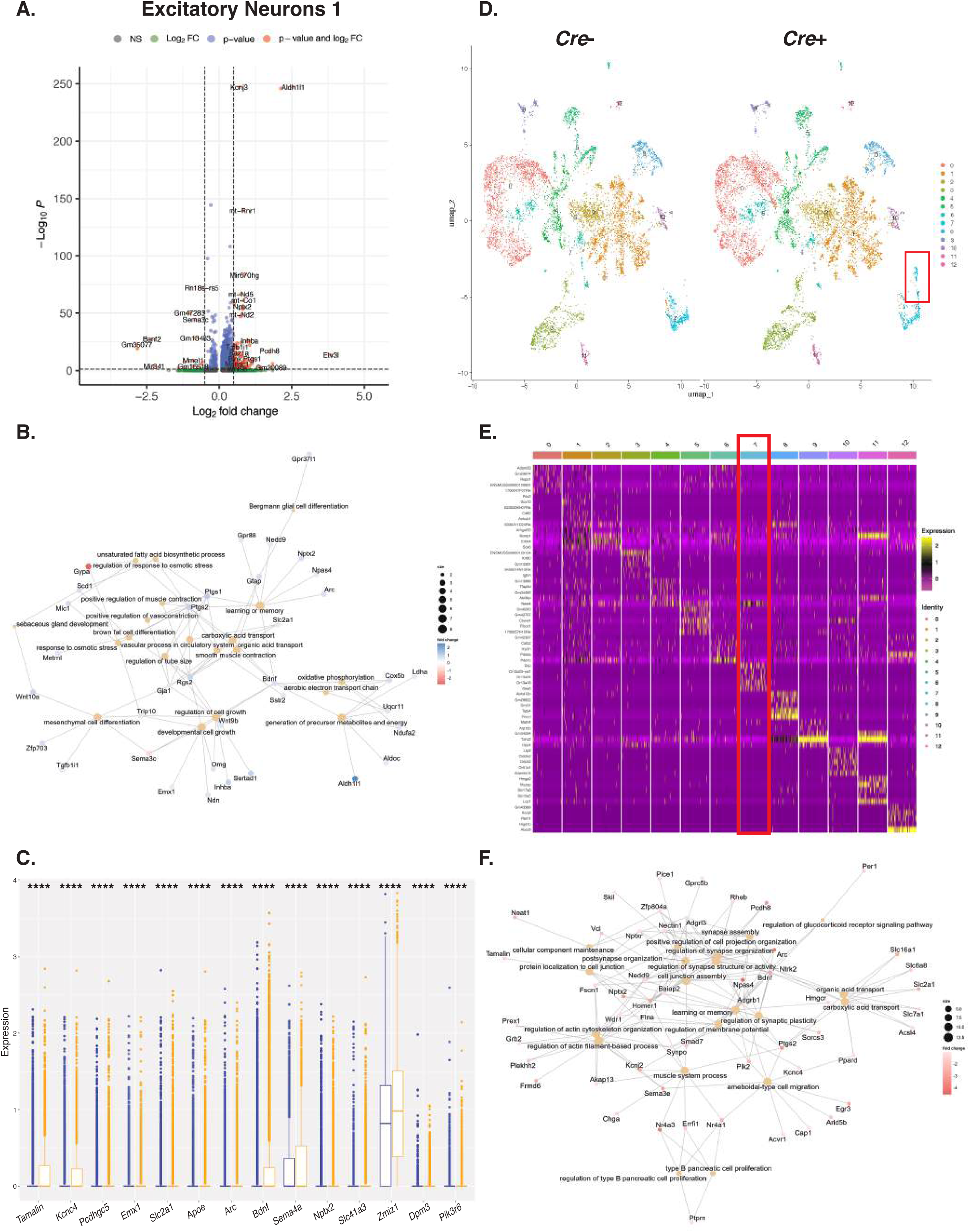
Description of the excitatory neurons 1 (EN1) population in PTHS brains. **(A)** A volcano plot representing the differential gene expression observed in the EN1 population is shown. The upregulated gene cutoff was log2fold change > 0.5 and Bonferroni adjusted *p-*value (p adj) < 0.05. The downregulated gene cutoff was log2fold change < −0.5 and p adj < 0.05. **(B)** Network/pathway analysis of the DEGs identified by comparing the expression profiles of Cre+ and Cre- EN1 clusters. **(C)** Box plots showing the most differentially regulated genes identified in the EN1 population. Blue boxes represent Cre- cells, and orange boxes represent Cre+ cells. Statistical analysis was done using R-suit. \*\*\*\**p*< 0.0001. **(D)** UMAP plots of the EN1 cells of Cre- (left) and Cre+ (right) mice are shown. Each set of data includes both male and female mice. Thirteen subclusters were identified. The red rectangle indicates the unique population present in Cre+ cells. **(E)** Heatmap showing the top five differentially expressed genes identified after comparing the expression profiles of the Cre+ and Cre- EN1 subclusters. Each row indicates a gene, and each column belongs to a cell within a subcluster. The subcluster IDs are color coded at the top of the heatmap. The red rectangle highlights the subcluster in which a new population of cells was found. **(F)** Network/pathway analysis of the DEGs identified by comparing the expression profiles of the cells in Cre+ and Cre- EN1 subcluster 7.

To further characterize the part of the EN1 cluster that was enriched in the Cre+ genotype, additional clustering was performed, resulting in a total of 13 subclusters within this population (Figure 5D and E). Through this analysis, we found that subcluster 7 was only identified in Cre+ EN1 cells (Figure 5D, red rectangle). To confirm the origin of subcluster 7, we performed pseudotime analysis and determined that these cells were from the same lineage as the other EN1 cells but had a unique transcriptional profile (Supplemental Figure 7B). GO (Supplemental Figure 7C) and network/pathway (Figure 5F) analysis of EN1 subcluster 7 showed that the differentially expressed genes in this population are broadly associated with synaptic function and learning and memory.

In addition to EN1, we also examined other neuronal populations that might contribute to the seizure phenotype in our PTHS model. Our snRNA-seq analysis indicated that the excitatory neurons 2 (EN2) and inhibitory neuron populations had increased *Aldh1l1* expression in Cre+ compared to Cre- brains (Figure 4D), suggesting that deletion of *Tcf4* might affect gene expression in these cell types. We found that there were more cells in the EN2 population in Cre+ brains (Supplemental Figure 4B) and that these cells had high numbers of DEGs when compared to Cre- EN2 cells (Figure 6A; Supplemental Table 5). GO enrichment (Supplemental Figure 8A) and network/pathway analysis (Figure 6B) of the genes that were differentially expressed in Cre+ EN2 cells compared to Cre- EN2 cells indicated that these genes are involved in ion transport, angiogenesis, and membrane functions. Interestingly, we identified the epilepsy-associated gene *Slc41s3* as one of the top DEGs in Cre+ EN2 cells (Figure 6C). Additional DEGs that we identified (*Sytl2*, *Cdh13,* and *Cacnb2)* are correlated with intellectual disability and neurodevelopmental disorders (Kessi et al., 2021; Rafi et al., 2019; Rivero et al., 2015). Intellectual disability has been characterized in all patients with PTHS (Blake et al., 2010; Brockschmidt et al., 2007; Forrest et al., 2018; Sepp et al., 2017) and in other PTHS mouse models (Li et al., 2019; Phan et al., 2020; Thaxton et al., 2018); therefore, EN2 cells could be contributing more to the development of this phenotype.

**Figure 6.**
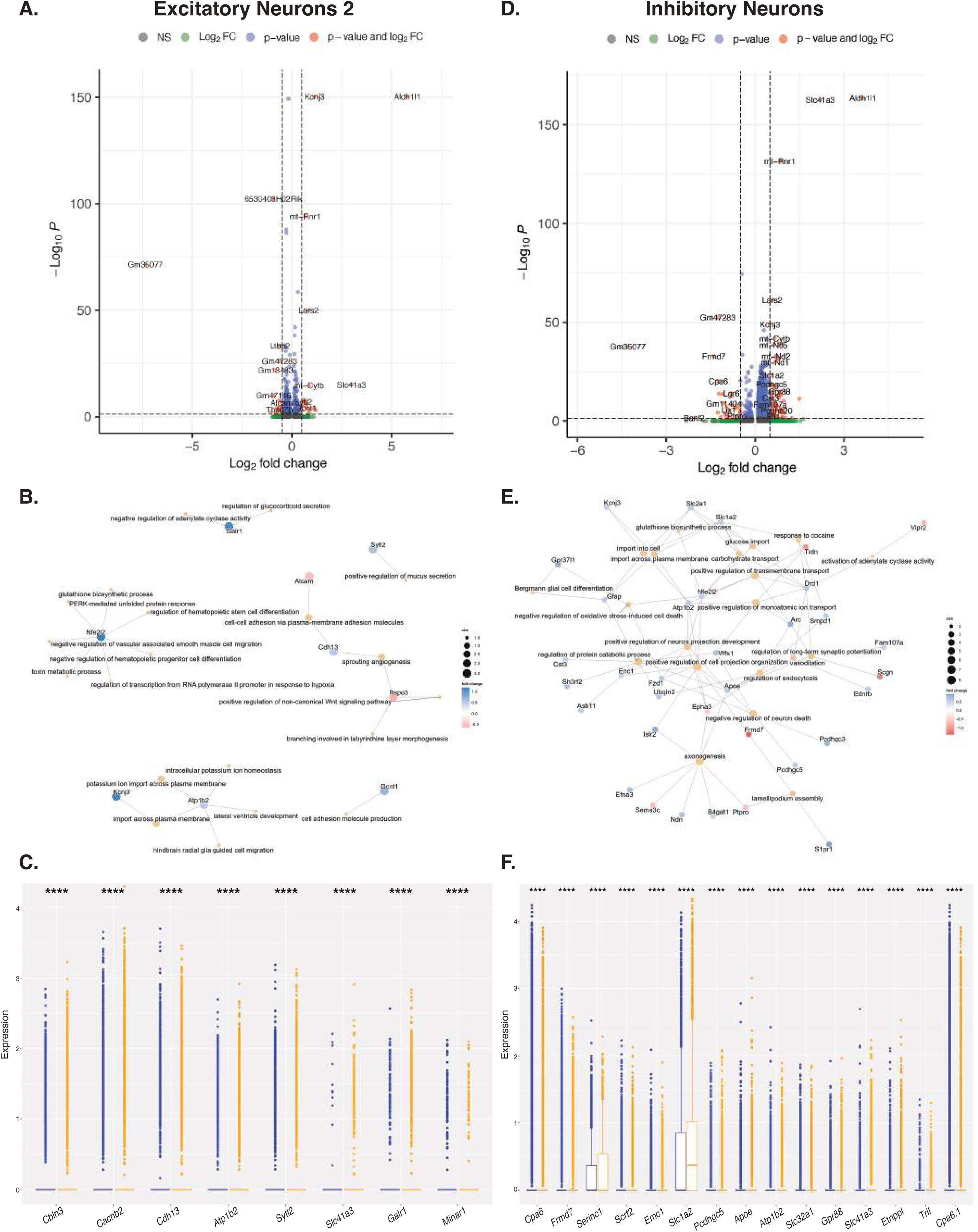
Analysis of the DEGs in the excitatory neurons 2 (EN2) and inhibitory neuron clusters in PTHS brains. **(A and D)** Volcano plots illustrating the differential gene expression observed between Cre+ and Cre- mice in the EN2 (left) and inhibitory neuron (right) clusters are shown. Dots represent differentially expressed genes. The upregulated gene cutoff was log2fold change > 0.5 and Bonferroni adjusted *p-*value (p adj) < 0.05, and the downregulated gene cutoff was log2fold change < −0.5 and p adj < 0.05. **(B and E)** Network/pathway analysis of the DEGs identified comparing Cre+ to Cre- in EN2 (B) and inhibitory neuron (E) clusters. **(C and F)** Box plots showing the most differentially regulated genes identified in the EN2 (C) and inhibitory neuron (F) populations. Blue boxes represent Cre- cells, and orange boxes represent Cre+ cells. Statistical analysis was done using R-suit. \*\*\*\**p*< 0.0001.

Along with excitatory neurons, inhibitory neurons (INs) are also essential for coordinating the balance between excitatory and inhibitory activities within brain circuits (Chen et al., 2023; Wang et al., 2022), and thus could contribute to the seizure phenotype seen in Cre+ mice. Of the genes that were differentially expressed in Cre+ INs compared to Cre- INs, most of them were upregulated, with a few that were downregulated (Figure 6D and Supplemental Figure 5). GO enrichment (Supplemental Figure 8B) and network/pathway analysis (Figure 6E) of the genes that were differentially expressed in the Cre+ IN cluster versus the Cre- IN cluster indicated connections to neuron projection development and organization, synaptic potentiation, and membrane transport. Similar to the DEGs identified in the EN1 and EN2 clusters, many of the top DEGs in Cre+ INs (Figure 6F and Supplemental Table 6) are associated with epilepsy (*Cpa6*, *Serinc1*, *Slc1a2*, *Pcdhgc5*, *Atp1b2*, *Slc32a1*, *Slc41a3*, *Etnppl*, and *Tril*) (Baaij, 2015; Beamer et al., 2021; Dell’Isola et al., 2022; Heron et al., 2021; Inuzuka et al., 2005; Sapio et al., 2015; Stergachis et al., 2019; White et al., 2021; Zhang et al., 2022) and intellectual disability (*Frmd7*, *Scrt2* and *Emc1)* (Dai et al., 2024; Goes et al., 2024; Huang et al., 2022). These results suggest that IN cells could be contributing to the epilepsy and intellectual disability phenotypes observed in PTHS patients.

Overall, our neuron cluster data analysis led to the identification of a number of genes in the EN1, EN2, and IN cell populations in Cre+ mice that regulate synaptic functions, memory and learning, neuron development, and membrane potential. Given that many of the DEGs identified here have previously been associated with epilepsy and intellectual disability, it is possible that they may also be involved in the development of epilepsy and other phenotypes in PTHS. However, further validation is required.

### 2.6. Identification of DEGs in glial cell clusters

There is growing evidence linking glia—especially astrocytes and oligodendrocytes—to epilepsy pathology (Çarçak et al., 2023; Knowles et al., 2022; Vezzani et al., 2022). Therefore, we examined the gene expression patterns in these cell types in Cre+ versus Cre- mice. In both the astrocyte and oligodendrocyte clusters of Cre+ mice, we identified significant numbers of DEGs (Supplemental Tables 7 and 8, respectively). The DEGs we detected in Cre+ astrocytes compared to Cre-astrocytes were primarily upregulated (Figure 7A and Supplemental Figure 5). GO enrichment (Supplemental Figure 9A) and network/pathway analysis (Figure 7B) further indicated that the DEGs identified in our Cre+ astrocyte cluster were broadly associated with synaptic function, membrane organization, and memory. Interestingly, we also found some DEGs, such as *lqsec1*, *Kif5a*, and *Slc41a3*, that are associated with various forms of epilepsy (Figure 7C)(Ansar et al., 2019; Baaij, 2015; Fukuoka et al., 2021).

**Figure 7.**
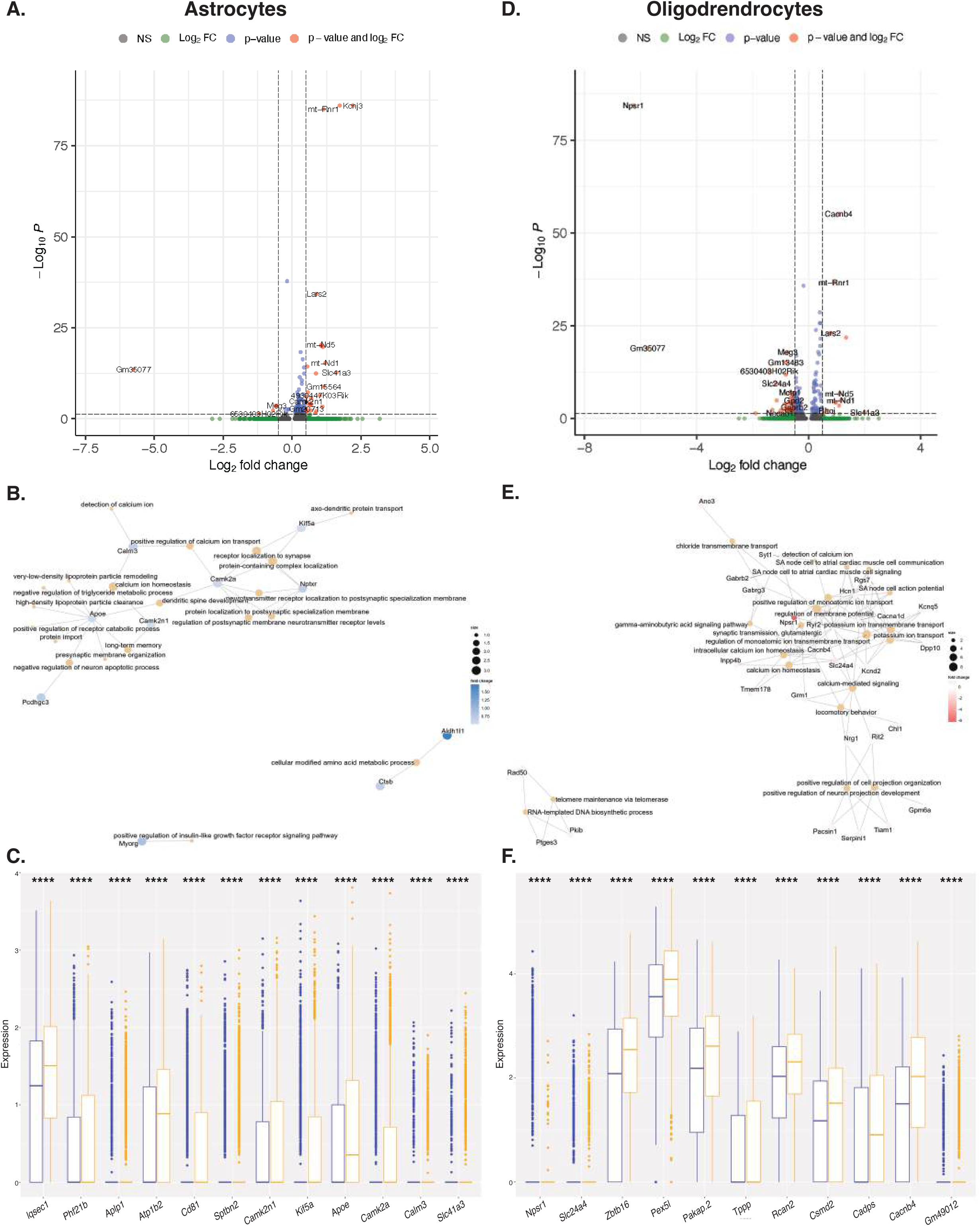
Analysis of the DEGs in astrocytes and oligodendrocytes in PTHS brains. **(A and D)** Volcano plots representing the genes that were differentially expressed in Cre+ compared to Cre-astrocytes (left) and oligodendrocytes (right) are shown. Dots represent differentially expressed genes. The upregulated gene cutoff was log2fold change > 0.5 and Bonferroni adjusted *p*-value (p adj) < 0.05. The downregulated gene cutoff was log2fold change < −0.5 and p adj < 0.05. **(B and E)** Network/pathway analysis of the DEGs identified in Cre+ versus Cre-astrocyte (B) and oligodendrocyte (E) clusters. **(C and F)** Box plots showing the most differentially regulated genes identified in astrocytes (C) and oligodendrocytes (F). Blue boxes represent Cre- cells, and orange boxes represent Cre+ cells. Statistical analysis was done using R-suit. \*\*\*\**p*<0.0001.

Previous work on other models of PTHS have found that TCF4 is necessary for the normal maturation and function of oligodendrocytes(Phan et al., 2020; Wedel et al., 2020). The potential contribution of oligodendrocytes to seizures in animal models has also been reported(Knowles et al., 2022). Moreover, our snRNA-seq data showed that *Aldh1l1* is expressed in oligodendrocytes in our Cre+ mice, so there may also be reduced expression of *Tcf4* that impacts gene expression in this cell type (Figure 4D). As expected, we found that oligodendrocytes showed significant differences in gene expression between Cre+ and Cre- cells (Figure 7D). GO enrichment (Supplemental Figure 9B) and network/pathway analysis (Figure 7E) of the identified DEGs indicated associations with ion transport, action potential signaling, and membrane potential regulation. Many of the top DEGs are also associated with epilepsy (*Cacnb4*, *Rcan2*, and *Pakap.2*)(Delgado-Escueta et al., 2013; Lee and Ahnn, 2020; Silva et al., 2024) or intellectual disability (*Zbtb16*, *Tppp*, *Csmd2* and *Cadps)* (Gutierrez et al., 2019; Oláh et al., 2017; Sitbon et al., 2022; Usui et al., 2021) (Figure 7F). These data suggest that the gene expression changes in astrocytes and oligodendrocytes may also contribute to seizures and other phenotypes in PTHS.

## 3. DISCUSSION

To date, there are many reports of PTHS mouse models that have abnormalities in local field potential recordings and EEGs (Kim et al., 2022), intrinsic excitability (Rannals et al., 2016), inhibitory network development (Chen et al., 2023), and synaptic transmission (Cleary et al., 2021; Li et al., 2019; Papes et al., 2022). Therefore, it is surprising that no other work has seen evidence of seizures. To the best of our knowledge, our largely astrocytic heterozygous *Tcf4* knockout mice are the first PTHS model to show behavioral seizures in almost all male and female Cre+ mice (93.4% and 77.8%, respectively; Figure 1E). In human PTHS patients, the prevalence of seizures is estimated to be between 30–50% (Matricardi et al., 2022; Winter et al., 2016), and PTHS patients without seizures sometimes show abnormal EEG activity without distinctive characterization as well (Matricardi et al., 2022; Rosenfeld et al., 2009). Moreover, similar to our PTHS mice in which we saw seizure onset occur between 3 and 12 months of age, seizure onset in humans is very variable, ranging from the first year of life to as late as early adulthood (Marangi et al., 2011; Pontual et al., 2009). The types of seizures experienced are also variable among patients, ranging from epileptic spasms (Pontual et al., 2009), to tonic-clonic (Matricardi et al., 2022; Zweier et al., 2008), absence (Rosenfeld et al., 2009; Winter et al., 2016), and atonic (Taddeucci et al., 2010) seizures. Since some of these seizures can be difficult to detect by eye—such as absence seizures—they may have been missed in the analysis of our model. Based on the muscle stiffening and twitching/jerking movements that we see in our mice, the seizures seem to align with descriptions of tonic-clonic seizures. However, without electroencephalogram data, we cannot say for sure. Nevertheless, the clear seizures that we observe make our PTHS model a good tool for future studies of both PTHS pathology and epilepsy.

Expression of c-Fos is representative of neuronal activity (Dragunow and Faull, 1989; Madabhushi et al., 2015; Zhang et al., 2002). Since c-Fos staining was seen in multiple brain regions, it was hard to pinpoint the exact seizure foci in our PTHS mice. However, it was unsurprising that we saw significant numbers of c-Fos+ cells in the cortex and hippocampus, as it is known that TCF4 is expressed at high levels in these brain regions, both embryonically and postnatally (Kim et al., 2020; Mesman et al., 2020; Page et al., 2018; Papes et al., 2022; Zhang et al., 2021). In addition, other models of PTHS have reported abnormal development and maturation of the cortex and hippocampus, as well as deficiencies in learning and memory behaviors that are associated with these brain regions (Forrest et al., 2018, 2014; Page et al., 2018; Papes et al., 2022; Schoof et al., 2020; Zhang et al., 2021). Though less is known about TCF4 functioning in the amygdala and hypothalamus, it is important to note that there are severe types of seizures, similar to those that we observed in our PTHS mice (limb stiffening and shaking), that implicate these brain regions (Aroniadou-Anderjaska et al., 2008; Bian et al., 2023). Therefore, more work is needed to clarify the roles of these brain areas in PTHS, as this information could be important for improving our understanding of disease pathogenesis.

Astrogliosis is often observed in human patients and in nearly every mouse model of epilepsy (Coulter and Steinhäuser, 2015; Patel et al., 2019; Robel et al., 2015). Many neurodevelopmental disorders that are similar to PTHS, such as Rett Syndrome, also report finding reactive astrogliosis in mouse models (Lioy et al., 2011; Maezawa et al., 2009; Petrelli et al., 2016; Rakela et al., 2018). Therefore, there is currently much interest in determining whether astrogliosis is a pathology that occurs before or after the onset of seizures. It has been traditionally thought that astrogliosis is a consequence of epileptic activity, as astrocytes are direct sensors of changes in the brain milieu, such as hyperexcitability (Robel and Sontheimer, 2016; Verhoog et al., 2020). However, there is growing evidence that direct changes in astrocytes can cause seizures or increased seizure risk (Robel et al., 2015, 2009; Uhlmann et al., 2002). Since we only examined one time point and observed both increased neuronal activity and astrogliosis, it is unclear whether astrogliosis was a cause or a consequence of the seizures observed in our PTHS model. One potential mechanism for PTHS pathogenesis is that astrocytes adopt reactive states due to the decreased expression of *Tcf4*, which then cause seizures. However, it is also possible that there are other changes that happen in PTHS mice that cause seizures, thus indirectly leading to astrogliosis. For example, it is known that high levels of neuronal activity, which are observed in our PTHS mice (via increased c-Fos staining (Figure 3)) and also in epilepsy (Devinsky et al., 2013; Rannals et al., 2016), can cause astrogliosis. Overall, improving our understanding of astrogliosis in PTHS epilepsy could be important for increasing our mechanistic knowledge of how the disease develops, and, ultimately, how to treat it.

It is now understood that not only neurons but also numerous other cell types are involved in seizure development, including astrocytes and oligodendrocytes (Knowles et al., 2022; Vezzani et al., 2022). Thus, we sought to identify potential molecular mechanisms for seizure development both globally and in specific cell types. Most notably, we found that *Kcnj3* (potassium inwardly rectifying channel subfamily J member 3) was differentially expressed across cell types (Figure 4E). This gene encodes GIRK1 (G-protein coupled inwardly rectifying potassium channel, also known as Kir3.1), which is an integral membrane protein that is broadly distributed in multiple cell types in the brain and controls membrane potential by inhibiting action potential firing by hyperpolarizing the plasma membrane (Chan et al., 1996; Signorini et al., 1997; Zhang et al., 2014). Interestingly, mutations in *Kcnj3*, and subsequent impairment of GIRK1 channel function, have been associated with epilepsy susceptibility (Li et al., 2023; Yamada et al., 2012). Therefore, further mechanistic work on this gene could reveal important casual information on seizure development and progression in patients with PTHS. Moreover, in our global snRNA-seq analysis, we identified a new population of primarily Cre+ cells within the EN1 cluster of excitatory neurons. While examination of the expression profile of this population did not reveal the exact genes that are involved in epilepsy, GO analysis determined that the genes that were differentially expressed in Cre+ mice were related to learning and memory and could therefore explain the intellectual disability observed in PTHS. It also identified genes related to synaptic function, which is notable since TCF4 has been shown to regulate synaptic development, structure, and density in other PTHS mouse and fly models (Badowska et al., 2020; Chen et al., 2016; Crux et al., 2018; Davis et al., 2024; D’Rozario et al., 2016; Kennedy et al., 2016; Schoof et al., 2020; Sepp et al., 2017; Thaxton et al., 2018). We also found many genes that were differentially expressed in the Cre+ EN2, inhibitory neuron, astrocyte, and oligodendrocyte clusters that we identified. Consistent with our phenotype, many of these genes are associated with epilepsy and neurodevelopmental disorders. Given that TCF4 is a transcription factor, previous work on other models of PTHS sought to identify TCF4-regulated genes and found differential expression of genes associated with neurodevelopment, synaptic functions, and myelination (Dennis et al., 2019; Jung et al., 2018; Mary et al., 2018; Phan et al., 2020; Torshizi et al., 2019; Wittmann and Häberle, 2018). Interestingly, few of these genes are associated with epilepsy, which is in contrast to the genes identified in our PTHS mouse model. However, though we have identified several candidate genes, our data is insufficient to conclude whether they are responsible for the seizure phenotype observed in our PTHS mice.

Overall, the current study is the first to generate a PTHS mouse model in which spontaneous and recurrent seizures are present; however, more work is needed to determine the etiology of this phenotype and completely understand the TCF4-dependent mechanisms that contribute to PTHS pathophysiology. Future mechanistic analyses of TCF4 should provide new insights into PTHS development and progression as well as epilepsy pathogenesis, which should help identify potential therapeutic strategies.

## 4. METHODS & MATERIALS

### 4.1. RESOURCE AVAILABILITY

#### 4.1.1. Lead contact

Further information and requests for resources and reagents should be directed to and will be fulfilled by the lead contact.

#### 4.1.2. Material availability

This study did not generate new unique reagents.

#### 4.1.3. Data and code availability

All snRNA-seq data will be posted on GEO.

### 4.2. SUBJECTS

#### 4.2.1. Animals

All animal care and procedures were approved by the University of California, Berkeley Animal Care and Use Committee and the Gladstone Institutes Institutional Animal Care and Use Committee. To generate mice with a *Tcf4*-knockout allele in astrocytes, we first purchased *Tcf4*tm1a(EUCOMM)Wtsi mice from the EMMA Mouse Repository (Skarnes et al., 2011). These mice contain an inserted cassette at position 69594442 of Chromosome 18 upstream of exon 4 that encodes a critical activation domain of *Tcf4*. This cassette is composed of an FRT-flanked LacZ/neomycin sequence followed by a loxP site. An additional loxP site is inserted downstream of the targeted exon at position 69595214. A Flp-deleter mouse purchased from Taconic was used to breed out the FRT-flanked sequence, resulting in a *Tcf4* conditional allele (*Tcf4^fl/fl^*). *Tcf4^fl/fl^*females were then bred with mice purchased from The Jackson Laboratory that constitutively express Cre recombinase under the control of the *Aldh1l1* promoter (Tg(*Aldh1l1*-Cre)) (Tien et al., 2012), resulting in mice with a heterozygous deletion of the long isoform of *Tcf4* (*Tcf4^fl/+^*^;^*Aldh1l1-Cre*) and littermate controls (*Tcf4^fl/+^*), which were used for our assays. The genotypes of the mice were determined by PCR using the primer sequences listed in Supplemental Table 2. Both male and female mice were used in this study. Mice were bred and housed in a specific pathogen-free animal facility of the Li Ka Shing Building at the University of California, Berkeley. All animals were weaned at around three weeks of age, housed in cages with 2–5 mice of the same sex, and given access to food and water *ad libitum*. As a result of the COVID-19 pandemic, the facility light cycle was changed to a 14-hour/10-hour light/dark cycle (on 6:00am; off 8:00pm).

### 4.3. METHODS

#### 4.3.1. Astrocyte isolation and RT-qPCR analysis

P4 pups were anesthetized on ice and sacrificed by decapitation. Tails were clipped for genotyping. Brains were extracted and put directly into ice cold 1X phosphate buffered saline (PBS; Thermo Fisher Scientific) in a dish. Five milliliters of dounce buffer (DB; 15 mM HEPES, 0.5% Glucose in HBSS(-)) containing 250 μL of 10,000 u/mL DNAse-I were added to the dish, then the brains were minced by feather blade and triturated with a flame-polished Pasteur pipette until the pieces were <1 mm in size. Whenever possible, samples were kept on ice, and centrifugation and sorting were performed at 4°C. Following trituration, samples were strained through a 70 μm cell strainer and washed with 4 mL of DB + DNase-I. Samples were then centrifuged at 600 x *g* for 5 minutes, and resuspended in MACS buffer (1X PBS, 2mM EDTA, 0.5% BSA). The Anti-GLAST MicroBead Kit (Miltenyi Biotec) was used, according to the manufacturer’s protocol, to isolate astrocytes from each sample. Following elution from the column, astrocytes were either processed for purity by flow cytometry (average purity was about 90%) or lysed to obtain RNA. Total astrocytic RNA was isolated using the quick-RNA Miniprep Kit (Zymo Research). Complementary DNA was reverse transcribed using SuperScript III (Invitrogen), and quantitative PCR (qPCR) was performed using SYBR green master mix (Roche) on a Quant Studio6 qPCR machine (ThermoFisher). Primers for qPCR were designed using Primer3 software and are listed in Supplemental Table 2. All data were collected with QuantStudio Real Time PCR Software v1.3 and normalized to the housekeeping gene hypoxanthine phosphoribosyltransferase (*Hprt*) using the delta-delta Ct method.

### 4.3.2. Analysis of spontaneous seizure behavior and lifespan

Mice were monitored weekly over the course of a year by examining them in their home cages for death or seizures. Those mice that had seizures were documented and later video recorded to better analyze the severity of their seizures. For the video recordings, adult mice (> three months of age) were moved from their home cage into a new individual cage and observed for 10 minutes with video recording. Mice were transferred back to their home cage at the end of the recording. Each mouse was recorded 3–5 times over 4 weeks to assess seizure severity, changes in seizure phenotypes, and reproducibility of the behaviors. From these videos, we selected the one that showed the most severe seizure for each mouse for analysis. Videos were analyzed in a double-blinded manner to quantify the time spent at each modified Racine Score, according to published scales (Racine, 1972). The criteria for the behavioral Racine scores used are shown in Figure 2A. Full tonic extinction or respiratory arrest were not observed during any video recordings. If there was a tie between videos, the first video recording collected was selected for analysis. Moderate scores were defined as scores of 3–4, severe scores were defined as scores of 5-6. Statistical analyses were performed using GraphPad Prism 10.

### 4.3.3. Immunohistochemistry of fixed tissue

Adult mice were anesthetized with avertin and transcardially perfused with 1X PBS. Brains were then recovered, weighed, and drop-fixed in 4% PFA (paraformaldehyde) in 1X PBS overnight, followed by transfer into 30% sucrose (in PBS) before freezing in Optimal Cutting Temperature mounting medium (OCT; Tissue Tek). Coronal sections of 30 m thickness were obtained using a cryostat (CM1950; Lecia Biosystmes) and placed directly onto Superfrost slides (ThermoFisher). Sections were washed 3 times with 0.1% Triton-X-100 in PBS (PBST; 10 minutes/wash), blocked by incubating in PBST containing 5% normal goat serum, and then incubated with primary antibodies diluted in PBST (Supplemental Table 3) for 18 h at 4°C. After washing 3 times with PBST (10 minutes/wash), brain sections were incubated with the appropriate secondary antibodies conjugated to Alexa Fluor® dyes (Supplemental Table 3) diluted in PBST, for 1 hour at room temperature. Sections were then stained with iamidino-2-phenylindole (DAPI; 1:10000) to label nuclei and mounted in Fluoromount-G Mounting Medium (SouthernBiotech). Images of whole sections were acquired with a Zeiss Axio ScanZ1 slide scanner using a 20x objective lens and Zen Blue 3.1 software (Carl Zeiss; UC Berkeley Molecular Imaging Center). Image analysis and quantification were performed using FIJI (National Institutes of Health) software.

#### 4.3.4. Nuclei extraction and preparation of snRNA-sequencing libraries

The snRNA-seq libraries were prepared following the protocol provided by the Evercode™ Fixation Kit (Parse Biosciences). Brain cell nuclei were isolated from 10-month-old adult mice (4–6 mice per group: Cre- males, Cre+ males, Cre- females, Cre+ females). First, mice were anesthetized and perfused with ice-cold 1X PBS. Each brain was extracted, weighed, and placed in a dish with ice-cold 1X PBS before being minced with a feather blade (one brain hemisphere excluding olfactory bulb and brain stem was used). Minced tissue was transferred to a Dounce homogenizer and homogenized in buffer (1.5 M NIM1, 1 mM DTT, 40 U/μl RNase-In, 20 U/μl Superase-In, 10% Triton-X-100) with both a loose and a tight pestle. The resulting homogenate was filtered through a 40 μm strainer into a 15 mL falcon tube and centrifuged for 4 minutes at 600 x *g* at 4°C. The supernatant was removed and the pellet was resuspended in 1 mL 1X PBS with RNAse inhibitors—Superase-In (Invitrogen) and enzymatic RNase inhibitor (Enzymatics)—and 10 mL of 1X PBS with 0.5% bovine serum albumin (BSA). Cells were then centrifuged and resuspended once more, before being passed through a 40 μm strainer into a fresh 15mL Falcon tube. Nuclei suspensions were then subjected to the Parse Biosciences Nuclei Fixation protocol starting at step 4 (Evercode™ Fixation User Manual v2.1.1). Samples were frozen for up to four weeks before being sent to the DNA Technologies and Expression Analysis Core at the University of California, Davis Genome Center for library preparation. Reverse transcription, split-pool barcoding, and library prep was performed using the Evercode™ WT Kit (Parse Biosciences), according to the manufacturer’s instructions. Quality control for each library was performed with a bioanalyzer (Agilent 2100). Libraries were run on a NovaSeq 6000 (Illumina) and paired-end sequencing was performed on two lanes of a Flow Cell, according to the instructions provided with the kit. Read one was done at 86 sequence cycles and read two was done at 64 cycles. Demultiplexing of sublibrary composition in sequencing reads was done using Bases2Fastq, with standard parameters for length of read 2 to 86 bp.

#### 4.3.5. Bioinformatic analysis of snRNA-sequencing data

For analysis of the snRNA sequencing data, raw sequencing read data first had to be processed using the split-seq-pipeline v0.9.6p from Parse Biosciences. Raw reads were aligned to the Mouse reference genome GRCm39, resulting in a matrix representing unique molecular identifiers (UMI’s) per cell barcode per gene. The raw UMI matrices for each sample were then processed using R v4.3.1 with the Seurat package v5.0.1. Matrices were filtered to remove cells with less than 200 genes detected and cells with >5% of reads mapping to mitochondrial RNA. Data from all samples were then integrated to combine and account for batch effects using the IntegrateLayers function using CCAIntegration as the integration method.

For the unsupervised clustering, the top 2000 most variably expressed genes were used for dimensionality reduction, first by principal component analysis (PCA) and then by uniform manifold projection (UMAP), selecting PCs 1:50 that explained the majority of the variance observed (assessed by elbow plots). A shared nearest-neighbor graph was constructed in PCA-space using PCs 1:50 with the Seurat FindNeighbors function. Clusters were identified within this graph using the Seurat FindClusters function, optimizing the modularity with the Louvain algorithm. The resolution parameter to control cluster granularity was manually set at 0.1. Cluster marker genes were identified with the FindAllMarkers function, only testing genes that were detected in a minimum fraction of 0.1 cells and using a logFC cutoff of 0.25. Cell cluster annotation was done by comparing the top marker genes per cluster with canonical marker genes from the literature.

For the DEG analysis, genes that were differentially expressed between groups were identified per cluster using Seurat’s FindMarkers function, utilizing the Wilcoxon Rank Sum test with default parameters. For the gene set enrichment analysis, GO term enrichment within clusters was determined using clusterProfiler, an R package. Finally, the trajectory pathway analysis was done via Monocle v3 for trajectory inference using the default parameters. Pseudotime was calculated using Cluster 4 as root cells.

## Supporting information

Supplemental information, figure compressed, table 1-8

## CRediT authorship contribution statement

L.C. designed the project, performed the experiments, analyzed data and wrote the manuscript with insights from all authors. V.R.D. analyzed seizure behavior. M.T.D. assisted in analyzing the sn-RNA-sequencing data. J.T.P. provided knowledge and feedback in designing the experiments. K.S. conceived the project and assisted in writing the manuscript.

## Acknowledgements

The authors appreciate Meik Kunz for the bioinformatical analyses. We thank Joseph Moreno and Cindy Wong for assisting with maintaining the animal colony. We would like to thank Agnieszka Ciesielska and Yuliya Voskobiynyk for guidance during Racine analysis. The authors also thank Dr. Amy Sullivan for editing the manuscript. This work was supported by the U.C. Dissertation Year Fellowship for L.C. and, NIH R21AG073735 and R21HD107388 for K.S.

