## Supplemental information, figure compressed, table 1-8 for "*Tcf4* Deficiency Causes Recurrent Seizures in Mice"

#### SUPPLEMENTAL FIGURE LEGENDS

**Supplemental Figure 1. Comparisons of the body and brain weights of Cre- and Cre+ mice.** **(A)** The average weights of 10–12-month-old male and female Cre- (blue) and Cre+ (orange) mice are shown (Cre- vs Cre+ males: ***p*=0.0089, Sidak's multiple comparisons test; Cre- vs Cre+ females: ns=not significant, *p*=0.2549, Sidak's multiple comparisons test). **(B)** The average weights of brains from 10–12-month-old male and female Cre- (blue) and Cre+ (orange) mice are shown (Cre- vs Cre+ males: ns=not significant, *p*=0.8765, Sidak's multiple comparisons test; Cre- vs Cre+ females: ns=not significant, *p*=0.0281, Sidak's multiple comparisons test). **(A and B)** Black circles denote individual male mice and black triangles denote individual female mice. Means ± SEM are represented.

**Supplemental Figure 2. Analysis of the genetic backgrounds of Cre- and Cre+ mice. (A)** Graph showing the distributions of J/J, J/N and N/N background genotypes for Cre- (blue) and Cre+ (orange) mice. ns=not significant, Fisher's exact test.

**Supplemental Figure 3. Regions analyzed for the immunostaining analysis. (A)** Example of coronal sections selected for analysis. Top row indicates a section from a Cre- mouse, and bottom row indicates a section from a Cre+ mouse, stained for c-Fos (red) and Gfap (green). Each scale bar indicates 1000 microns. **(B)** Schematic of a mouse coronal brain section indicating the areas that were used for the immunostaining analysis shown in Figure 3 (black dotted squares, 500 μm x 500 μm).

**Supplemental Figure 4. A detailed analysis of the brain cell types in our PTHS mouse model. (A)** The numbers of cells of the indicated cell types (x-axis) identified in each sample (23 samples total) are shown as box plots. Blue and orange boxes indicate the spreads of the numbers of cells identified in the Cre- and Cre+ samples, respectively. The *p*-values at the top of the graph indicate whether there is a significant difference between the number of Cre- and Cre+ cells for each cell type. Statistical analysis was done using R-suit. *$p<0.05$. **(B)** UMAP plots showing the cell-type clusters (as in Figure 4C) for cells from females (left) and males (right). Cre- and Cre+ cells were combined for these analyses.

**Supplemental Figure 5. Differential gene regulation in Cre+ mice.** The numbers of up-and downregulated genes identified in Cre+ mice compared to Cre- littermate controls is shown. Blue bars indicate downregulated genes, and orange bars indicate upregulated genes.

**Supplemental Figure 6. UMAP plots of the most significant DEGs in PTHS mice.** UMAP plots for *Aldh1l1* **(A)** and *Kcnj3* **(B)** expression in the Cre- (left) and Cre+ (right) cell clusters are shown.

**Supplemental Figure 7. Further characterization of the excitatory neurons 1 (EN1) cluster. (A)** GO analysis of the genes that were differentially expressed in Cre+ EN1 cells compared to Cre- EN1 cells. **(B)** Pseudotime analysis of EN1. **(C)** GO analysis of the DEGs identified in Cre+ EN1 subcluster 7.

**Supplemental Figure 8. GO analysis of the DEGs identified in the excitatory neurons 2 (EN2) and inhibitory neuron clusters.** GO analysis of the genes that were differentially expressed in the EN2 **(A)** and inhibitory neuron **(B)** clusters in Cre+ compared to Cre- mice.

**Supplemental Figure 9. GO analysis of the DEGs identified in the astrocyte and oligodendrocyte clusters.** GO analysis of the genes that were differentially expressed in the astrocyte **(A)** and oligodendrocyte **(B)** clusters in Cre+ compared to Cre- mice.

#### SUPPLEMENTAL TABLE LEGENDS

**Supplemental Table 1.** List of mice used for sn-RNA sequencing

**Supplemental Table 2.** List of sequences used for RT-qPCR and PCR primers

**Supplemental Table 3.** List of primary and secondary antibodies

**Supplemental Table 4.** Differentially expressed genes in excitatory neuron 1 (EN1) population

**Supplemental Table 5.** Differentially expressed genes in excitatory neuron 2 (EN2) population

**Supplemental Table 6.** Differentially expressed genes in inhibitory neuron (IN) population

**Supplemental Table 7.** Differentially expressed genes in astrocyte population

**Supplemental Table 8.** Differentially expressed genes in oligodendrocyte population

#### SUPPLEMENTAL VIDEO LEGENDS

**Supplemental Video 1.** Video showing one male mouse undergoing severe seizure.

**Supplemental Video 2.** Video showing one female mouse undergoing severe seizure.

**Supplemental Figure 1**

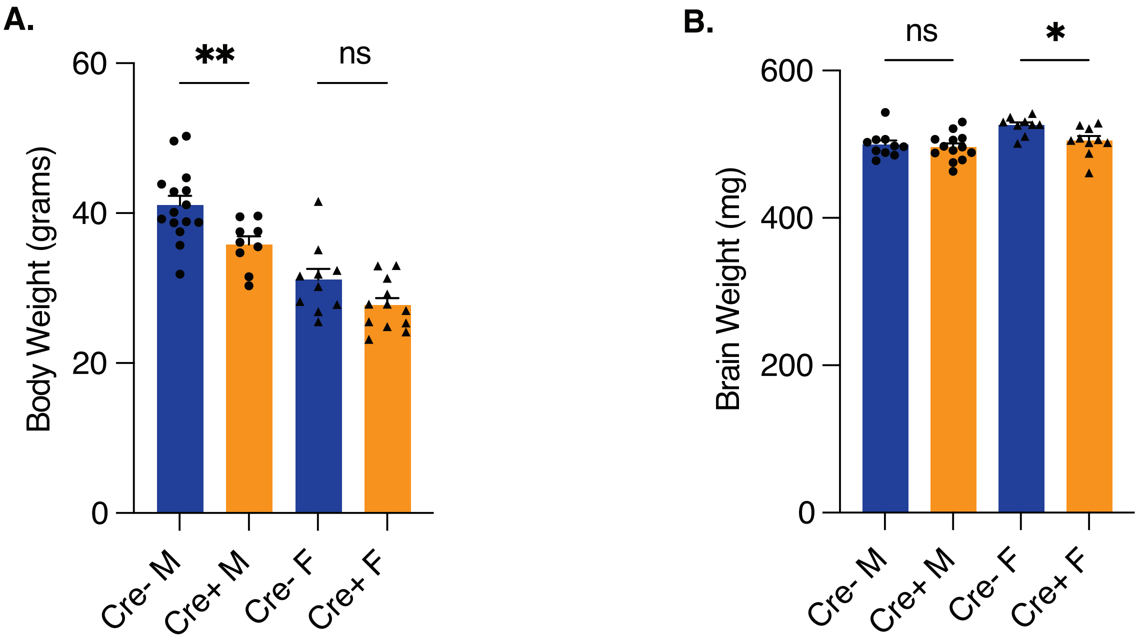

**Supplemental Figure 2**

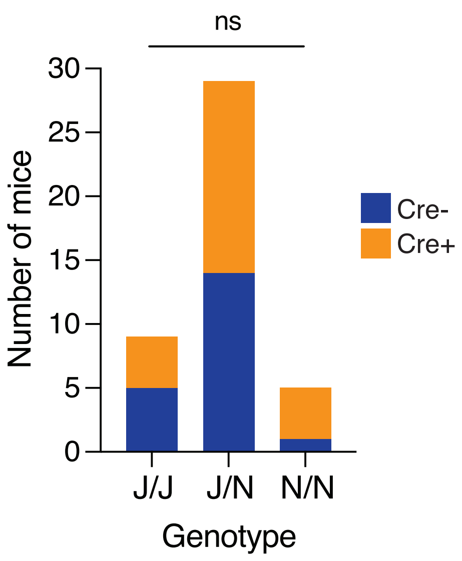

**Supplemental Figure 3**

**A.**

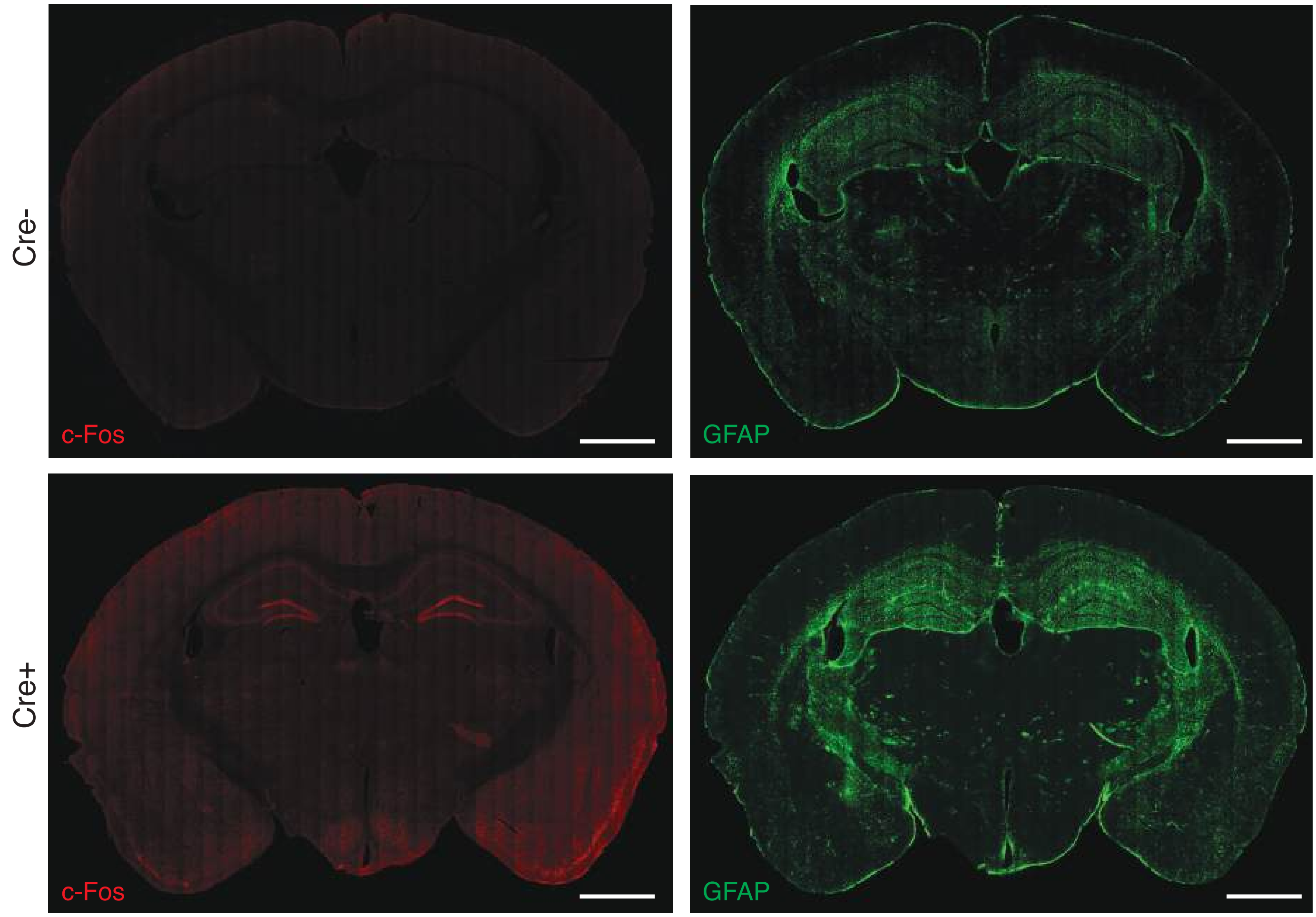


**B.**

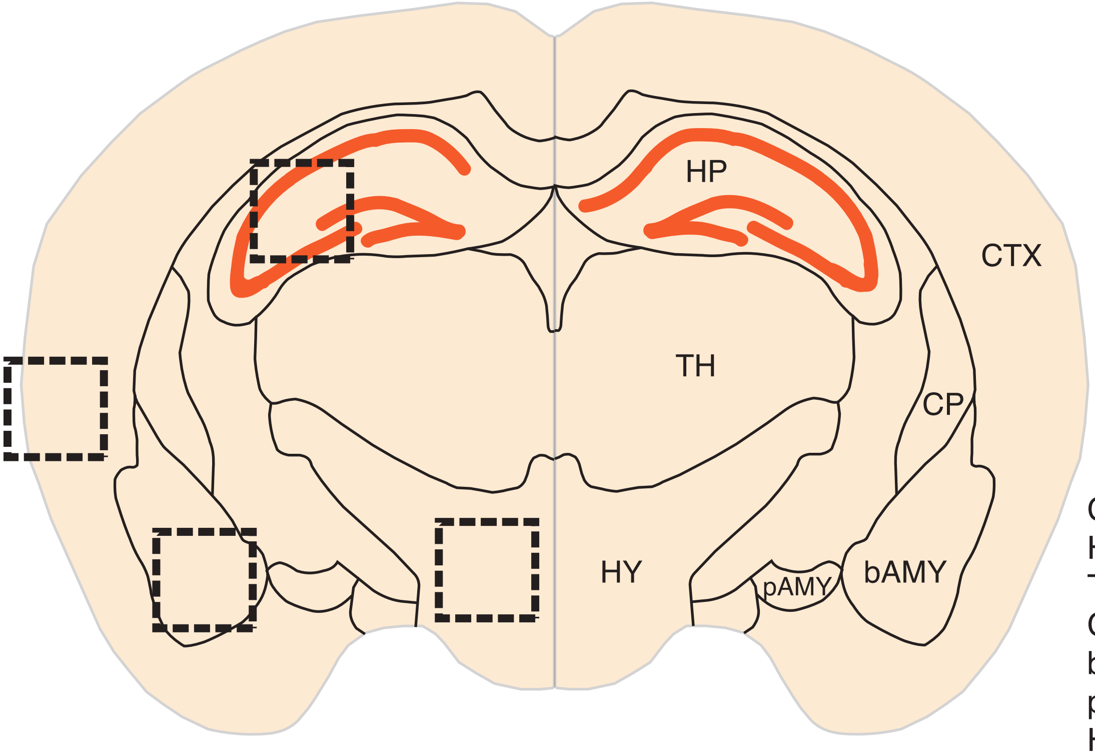


CTX: Cortex
HP: Hippocampus
TH: Thalamus
CP: Caudoputamen
bAMY: basolateral Amygdala
pAMY: posterior Amygdala
HY: Hypothalamus

#### Supplemental Figure 4

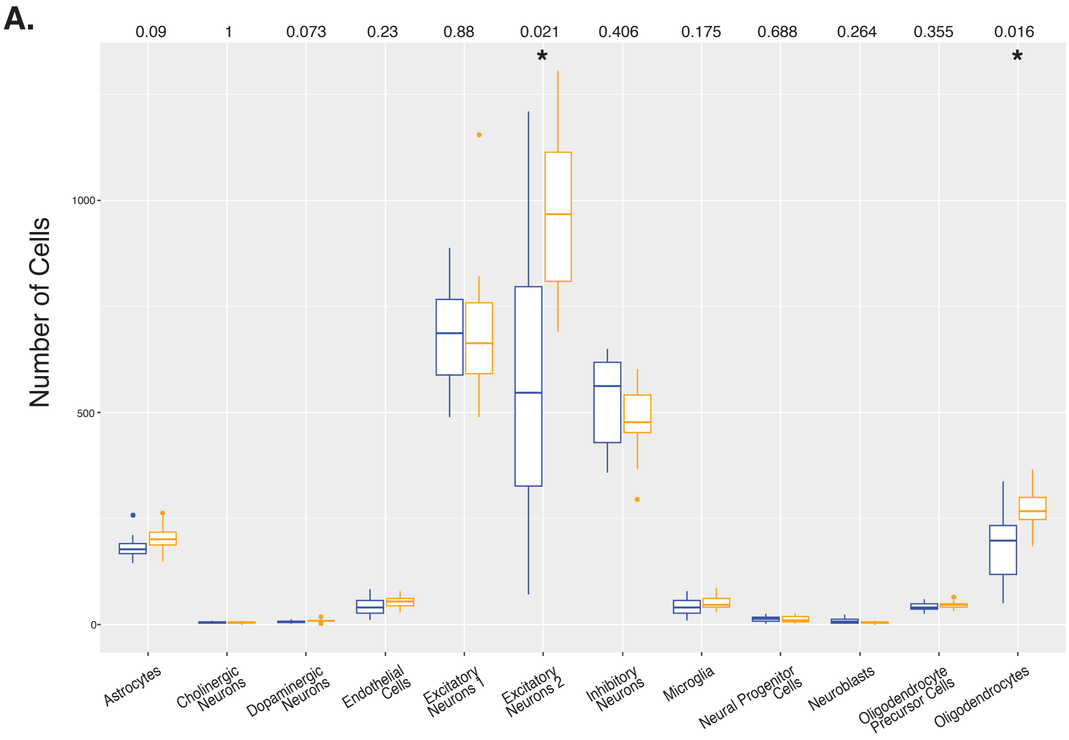


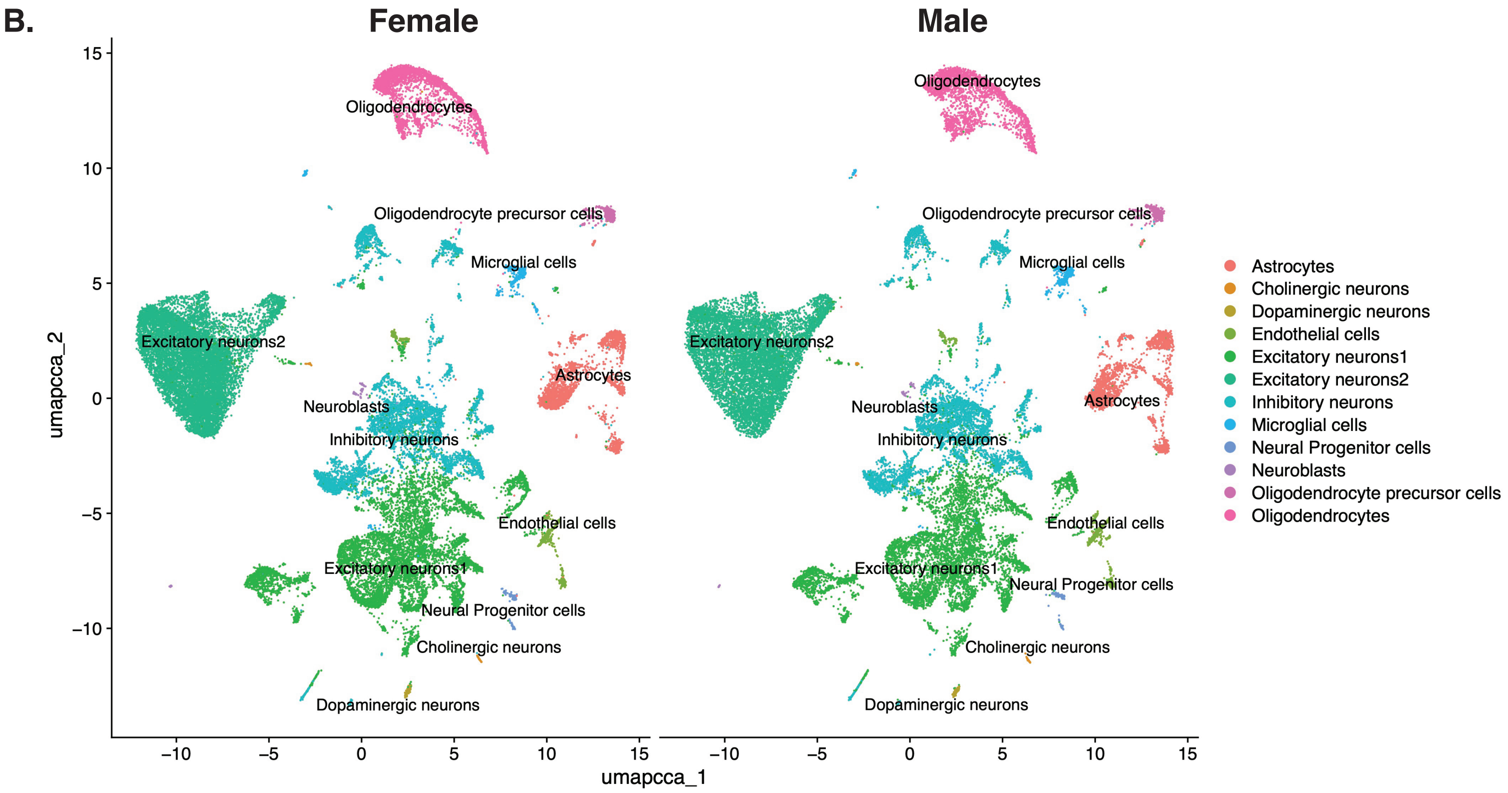

#### Supplemental Figure 5

**A.**

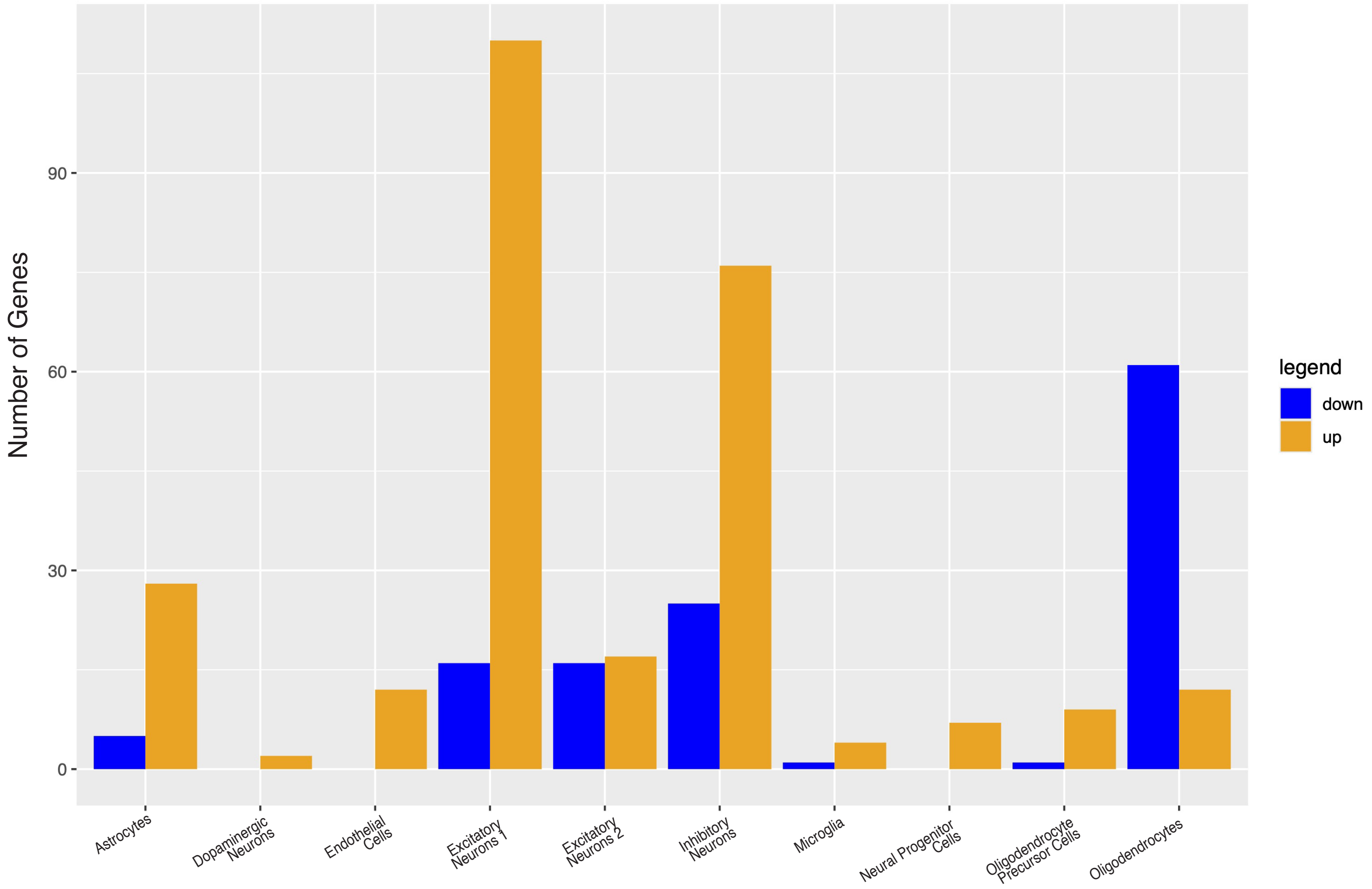

### Supplemental Figure 6

**A.** ***Aldh1l1***

*Cre-* *Cre+*

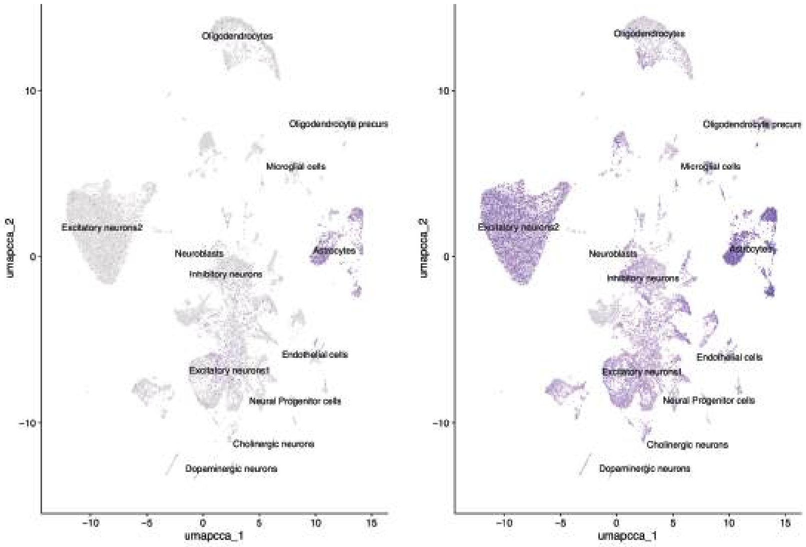


**B.** ***Kcnj3***

*Cre-* *Cre+*

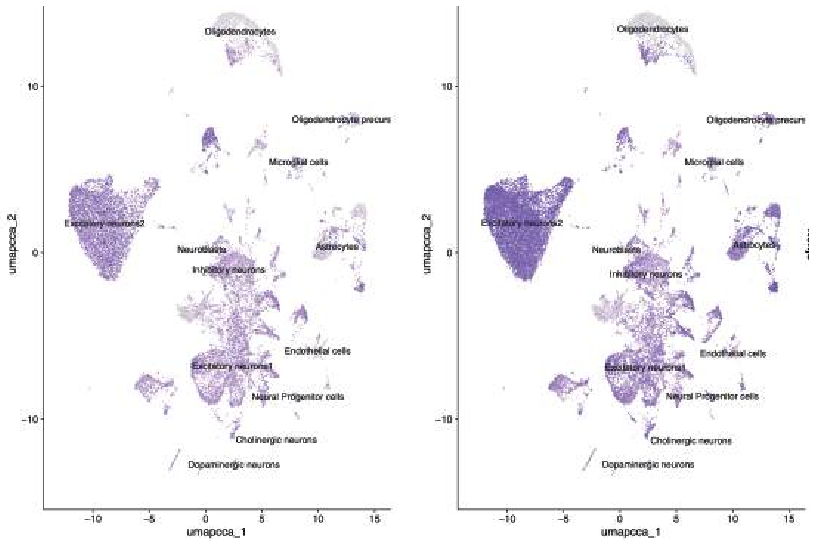

#### Supplemental Figure 7

**A.** **Excitatory Neurons 1**

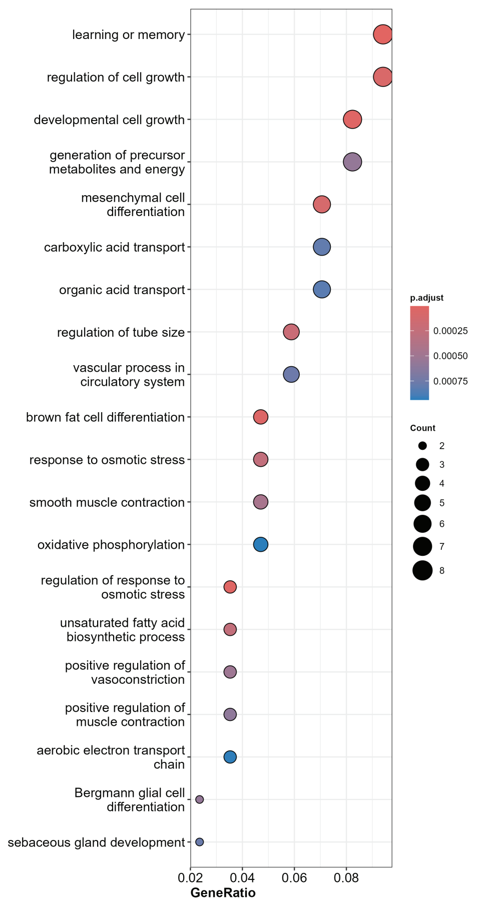


**C.** **Subcluster 7**

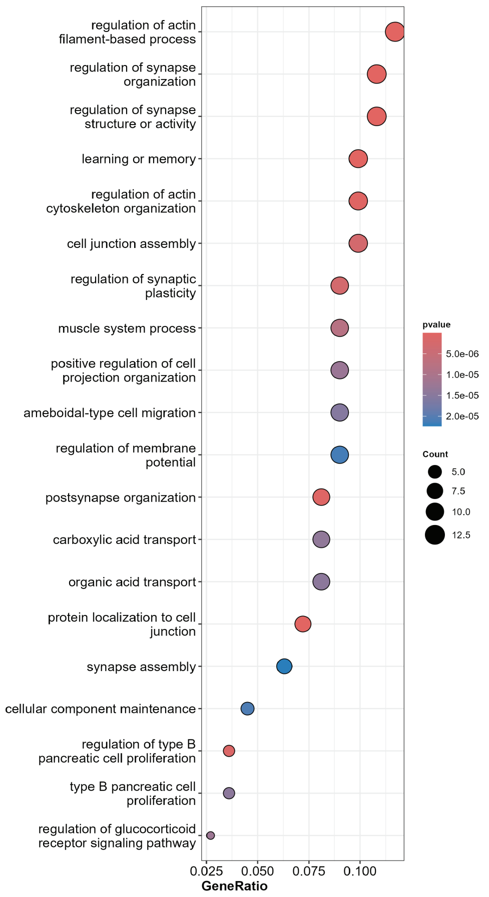


**B.**

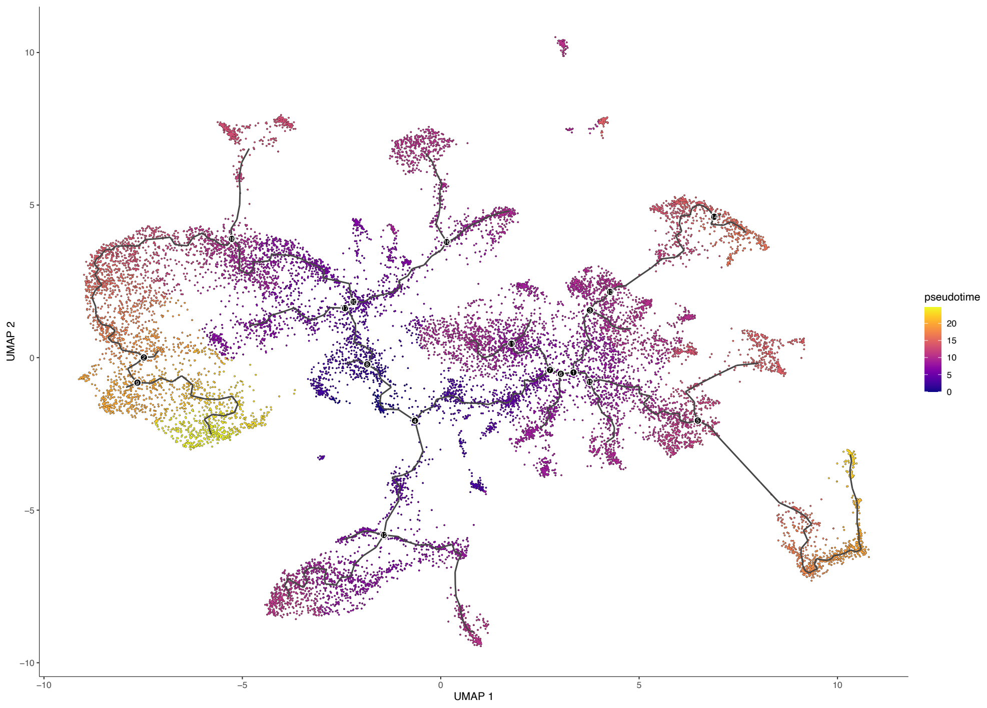

### Supplemental Figure 8

#### A. Excitatory Neurons 2

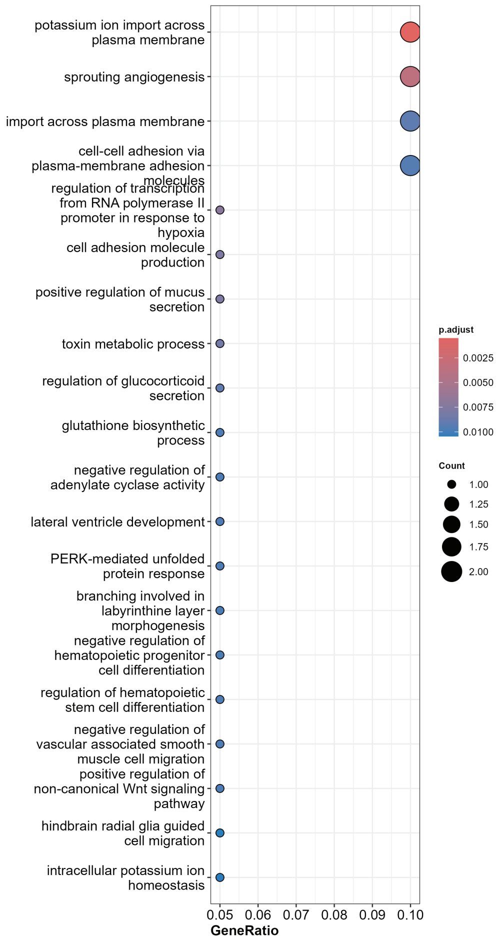


#### B. Inhibitory Neurons

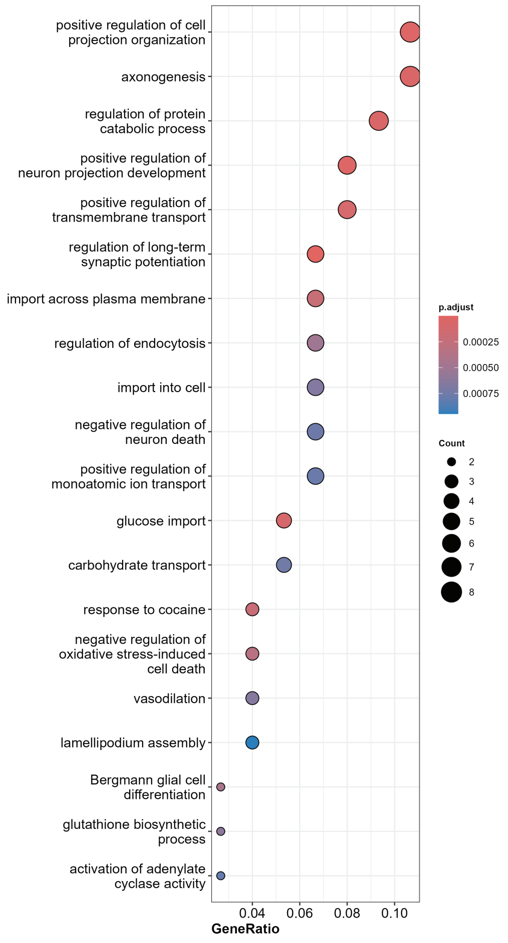

### Supplemental Figure 9

#### A. Astrocytes

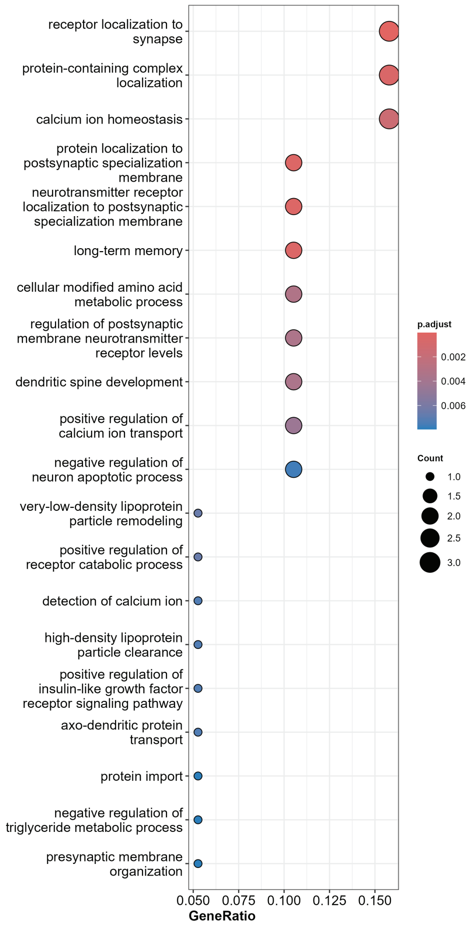


#### B. Oligodendrocytes

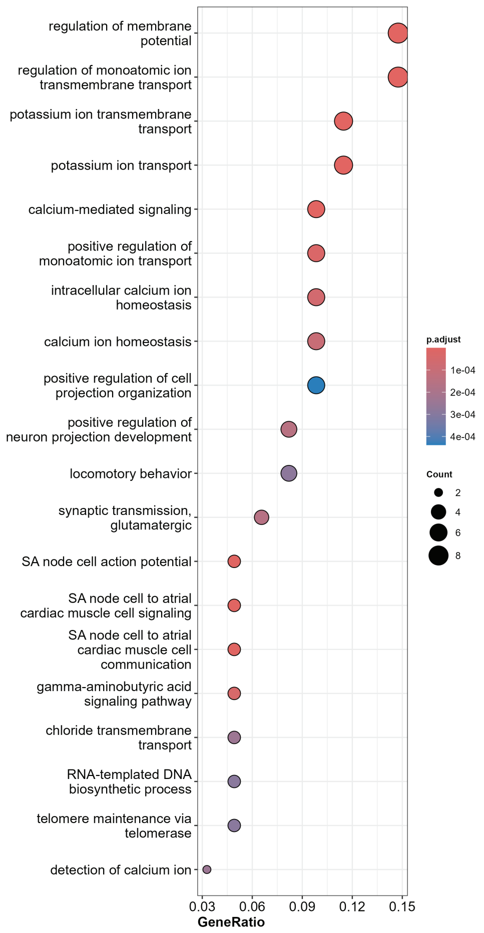

**Supplemental Table 1.** List of mice used for sn-RNA sequencing

| Identification | Sex | Age (months) | Genotype |
|---|---|---|---|
| Mouse 1 | M | 10 | *Tcf4*$^{fl/+}$ |
| Mouse 2 | M | 10 | *Tcf4*$^{fl/+}$;*Aldh1l1-Cre* |
| Mouse 3 | M | 10 | *Tcf4*$^{fl/+}$ |
| Mouse 4 | M | 10 | *Tcf4*$^{fl/+}$;*Aldh1l1-Cre* |
| Mouse 5 | F | 10 | *Tcf4*$^{fl/+}$ |
| Mouse 6 | F | 10 | *Tcf4*$^{fl/+}$ |
| Mouse 7 | M | 10 | *Tcf4*$^{fl/+}$ |
| Mouse 8 | M | 10 | *Tcf4*$^{fl/+}$ |
| Mouse 9 | F | 10 | *Tcf4*$^{fl/+}$ |
| Mouse 10 | F | 10 | *Tcf4*$^{fl/+}$ |
| Mouse 11 | F | 10 | *Tcf4*$^{fl/+}$;*Aldh1l1-Cre* |
| Mouse 12 | F | 10 | *Tcf4*$^{fl/+}$ |
| Mouse 13 | M | 10 | *Tcf4*$^{fl/+}$ |
| Mouse 14 | M | 10 | *Tcf4*$^{fl/+}$;*Aldh1l1-Cre* |
| Mouse 15 | M | 10 | *Tcf4*$^{fl/+}$;*Aldh1l1-Cre* |
| Mouse 16 | F | 10 | *Tcf4*$^{fl/+}$ |
| Mouse 17 | F | 10 | *Tcf4*$^{fl/+}$;*Aldh1l1-Cre* |
| Mouse 18 | F | 10 | *Tcf4*$^{fl/+}$;*Aldh1l1-Cre* |
| Mouse 19 | F | 10 | *Tcf4*$^{fl/+}$;*Aldh1l1-Cre* |
| Mouse 20 | M | 10 | *Tcf4*$^{fl/+}$;*Aldh1l1-Cre* |
| Mouse 21 | M | 10 | *Tcf4*$^{fl/+}$;*Aldh1l1-Cre* |
| Mouse 22 | F | 10 | *Tcf4*$^{fl/+}$;*Aldh1l1-Cre* |
| Mouse 23 | F | 10 | *Tcf4*$^{fl/+}$;*Aldh1l1-Cre* |

**Supplemental Table 2.** List of sequences used for RT-qPCR and PCR primers

| Gene | RT-qPCR Primer Sequences |
|---|---|
| *Hprt* | F: TCAGTCAACGGGGGACATAAA<br>R: GGGGCTGTACTGCTTAACCAG |
| *Tcf4* | F: ACGGACAAAGAGCTGAGTGA<br>R: CTCCAGTTCCCCAGGACC |
| **Gene** | **PCR Primer Sequences** |
| *Aldh1l1-Cre* | F: CCTGTCCCCTTGCACAGTAG<br>R: CGGTTATTCAACTTGCACCA |
| J (*Nnt* wild type) | F: GGGCATAGGAAGCAAATACCAAGTG<br>R: GTAGGGCCAACTGTTTCTGCATGA |
| N (*Nnt* mutation) | F: GTGGAATTCCGCTGAGAGAACTCTT<br>R: GTAGGGCCAACTGTTTCTGCATGA |
| *Tcf4* wild type | F: CCGATGACAGTGATGATGGT<br>R: AAGTTAAGCTGAAGTAAATACCCACA |
| *Tcf4* floxed | F: CCGATGACAGTGATGATGGT<br>R: TCGTGGTATCGTTATGCGCC |

**Supplemental Table 3.** List of primary and secondary antibodies

| Antibody | Concentration | Company |
|---|---|---|
| GFAP | 1:500 | ThermoFisher 13-0300 |
| c-Fos | 1:2000 | Abcam ab190289 |
| Goat anti-Rat Alexa Fluor® 488 | 1:500 | ThermoFisher A-11001 |
| Goat anti-Rabbit Alexa Fluor® 594 | 1:500 | ThermoFisher A-11005 |

Cre+_vs_Cre-_enrichGO

| | ONTOLOG | ID | Description | GeneRati | BgRatio | pvalue | p.adjust | qvalue | geneID | Count |
|---|---|---|---|---|---|---|---|---|---|---|
| GO:000761 | BP | GO:000761 | learning or memory | 8/85 | 326/2856 | ################ | ################ | 0.006387881 | Nptx2/Arc/Bdnf/Ptgs2/Nedd9/Ptgs1/Npas4/Gpr8 | 8 |
| GO:004748 | BP | GO:004748 | regulation of response to osmotic stress | 3/85 | 18/28564 | ################ | ################ | 0.006387881 | Ptgs2/Gypa/Mlc1 | 3 |
| GO:004858 | BP | GO:004858 | developmental cell growth | 7/85 | 287/2856 | ################ | ################ | 0.006387881 | Sema3c/Rgs2/Bdnf/Emx1/Omg/Ndn/Trip10 | 7 |
| GO:005087 | BP | GO:005087 | brown fat cell differentiation | 4/85 | 64/28564 | ################ | ################ | 0.008872596 | Rgs2/Ptgs2/Scd1/Metrnl | 4 |
| GO:000155 | BP | GO:000155 | regulation of cell growth | 8/85 | 464/2856 | ################ | ################ | 0.01382404 | Sema3c/Inhba/Rgs2/Bdnf/Omg/Gja1/Trip10/S | 8 |
| GO:004876 | BP | GO:004876 | mesenchymal cell differentiation | 6/85 | 249/2856 | 0.000101648 | 0.000101648 | 0.016705102 | Sema3c/Tgfb1i1/Gja1/Wnt10a/Zfp703/Trip10 | 6 |
| GO:003515 | BP | GO:003515 | regulation of tube size | 5/85 | 180/2856 | 0.000205101 | 0.000205101 | 0.026965368 | Rgs2/Ptgs2/Ptgs1/Gja1/Wnt9b | 5 |
| GO:000663 | BP | GO:000663 | unsaturated fatty acid biosynthetic process | 3/85 | 41/28564 | 0.000249819 | 0.000249819 | 0.02826643 | Ptgs2/Scd1/Ptgs1 | 3 |
| GO:000697 | BP | GO:000697 | response to osmotic stress | 4/85 | 103/2856 | 0.000257996 | 0.000257996 | 0.02826643 | Ptgs2/Gypa/Mlc1/Trip10 | 4 |
| GO:000693 | BP | GO:000693 | smooth muscle contraction | 4/85 | 119/2856 | 0.00044695 | 0.00044695 | 0.039174799 | Rgs2/Ptgs2/Ptgs1/Sstr2 | 4 |
| GO:004590 | BP | GO:004590 | positive regulation of vasoconstriction | 3/85 | 52/28564 | 0.000505841 | 0.000505841 | 0.041285371 | Ptgs2/Ptgs1/Gja1 | 3 |
| GO:000609 | BP | GO:000609 | generation of precursor metabolites and energy | 7/85 | 478/2856 | 0.000564291 | 0.000564291 | 0.041285371 | Aldh1l1/Bdnf/Aldoc/Ldha/Cox5b/Ndufa2/Uqcr1 | 7 |
| GO:006002 | BP | GO:006002 | Bergmann glial cell differentiation | 2/85 | 12/28564 | 0.000566508 | 0.000566508 | 0.041285371 | Gfap/Gpr37l1 | 2 |
| GO:004593 | BP | GO:004593 | positive regulation of muscle contraction | 3/85 | 55/28564 | 0.000596638 | 0.000596638 | 0.041285371 | Rgs2/Ptgs2/Ptgs1 | 3 |
| GO:000301 | BP | GO:000301 | vascular process in circulatory system | 5/85 | 240/2856 | 0.000762775 | 0.000762775 | 0.047309428 | Rgs2/Ptgs2/Slc2a1/Ptgs1/Gja1 | 5 |
| GO:004873 | BP | GO:004873 | sebaceous gland development | 2/85 | 14/28564 | 0.000778077 | 0.000778077 | 0.047309428 | Scd1/Wnt10a | 2 |
| GO:004694 | BP | GO:004694 | carboxylic acid transport | 6/85 | 367/2856 | 0.000804677 | 0.000804677 | 0.047309428 | Rgs2/Bdnf/Ptgs2/Slc2a1/Gja1/Gfap | 6 |
| GO:001584 | BP | GO:001584 | organic acid transport | 6/85 | 369/2856 | 0.000827631 | 0.000827631 | 0.047309428 | Rgs2/Bdnf/Ptgs2/Slc2a1/Gja1/Gfap | 6 |
| GO:001964 | BP | GO:001964 | aerobic electron transport chain | 3/85 | 64/28564 | 0.00092944 | 0.00092944 | 0.048726132 | Bdnf/Cox5b/Uqcr11 | 3 |
| GO:000611 | BP | GO:000611 | oxidative phosphorylation | 4/85 | 145/2856 | 0.00093782 | 0.00093782 | 0.048726132 | Bdnf/Cox5b/Ndufa2/Uqcr11 | 4 |
| GO:005193 | BP | GO:005193 | synaptic transmission, GABAergic | 3/85 | 68/28564 | 0.001108459 | 0.001108459 | 0.048726132 | Bdnf/Npas4/Hapln4 | 3 |
| GO:009033 | BP | GO:009033 | positive regulation of brown fat cell differentiation | 2/85 | 17/28564 | 0.001156114 | 0.001156114 | 0.048726132 | Ptgs2/Metrnl | 2 |
| GO:004592 | BP | GO:004592 | negative regulation of growth | 5/85 | 267/2856 | 0.001226393 | 0.001226393 | 0.048726132 | Sema3c/Inhba/Rgs2/Gja1/Sertad1 | 5 |
| GO:190352 | BP | GO:190352 | regulation of blood circulation | 5/85 | 270/2856 | 0.001288439 | 0.001288439 | 0.048726132 | Rgs2/Ptgs2/Ptgs1/Gja1/Pebp1 | 5 |
| GO:190486 | BP | GO:190486 | inhibitory synapse assembly | 2/85 | 18/28564 | 0.001298117 | 0.001298117 | 0.048726132 | Npas4/Hapln4 | 2 |
| GO:000719 | BP | GO:000719 | adenylate cyclase-inhibiting G protein-coupled receptor signaling | 3/85 | 72/28564 | 0.001307874 | 0.001307874 | 0.048726132 | Rgs2/Sstr2/Gpr37l1 | 3 |
| GO:004863 | BP | GO:004863 | regulation of developmental growth | 6/85 | 405/2856 | 0.001334214 | 0.001334214 | 0.048726132 | Sema3c/Rgs2/Bdnf/Omg/Gja1/Trip10 | 6 |
| GO:003109 | BP | GO:003109 | regeneration | 4/85 | 161/2856 | 0.001380846 | 0.001380846 | 0.049066184 | Bdnf/Omg/Gja1/Gfap | 4 |
| GO:003563 | BP | GO:003563 | maintenance of blood-brain barrier | 2/85 | 19/28564 | 0.001448036 | 0.001448036 | 0.049084683 | Ptgs2/Ptgs1 | 2 |
| GO:005080 | BP | GO:005080 | negative regulation of synaptic transmission | 3/85 | 77/28564 | 0.001587089 | 0.001587089 | 0.049084683 | Arc/Bdnf/Ptgs2 | 3 |
| GO:006056 | BP | GO:006056 | developmental growth involved in morphogenesis | 5/85 | 290/2856 | 0.001763172 | 0.001763172 | 0.049084683 | Sema3c/Bdnf/Emx1/Omg/Ndn | 5 |
| GO:007139 | BP | GO:007139 | cellular response to cholesterol | 2/85 | 21/28564 | 0.001771432 | 0.001771432 | 0.049084683 | Inhba/Mlc1 | 2 |
| GO:004666 | BP | GO:004666 | male sex differentiation | 4/85 | 174/2856 | 0.001834797 | 0.001834797 | 0.049084683 | Inhba/Spata2/Gja1/Wnt9b | 4 |
| GO:001470 | BP | GO:001470 | striated muscle tissue development | 5/85 | 296/2856 | 0.00192777 | 0.00192777 | 0.049084683 | Sema3c/Rgs2/Bdnf/Gja1/Trip10 | 5 |
| GO:001023 | BP | GO:001023 | vascular transport | 2/85 | 22/28564 | 0.001944816 | 0.001944816 | 0.049084683 | Slc2a1/Gja1 | 2 |
| GO:005502 | BP | GO:005502 | regulation of cardiac muscle tissue growth | 3/85 | 84/28564 | 0.002036639 | 0.002036639 | 0.049084683 | Rgs2/Gja1/Trip10 | 3 |
| GO:001620 | BP | GO:001620 | regulation of striated muscle tissue development | 2/85 | 23/28564 | 0.002125928 | 0.002125928 | 0.049084683 | Bdnf/Gja1 | 2 |
| GO:003072 | BP | GO:003072 | ovulation | 2/85 | 23/28564 | 0.002125928 | 0.002125928 | 0.049084683 | Inhba/Ptgs2 | 2 |
| GO:004867 | BP | GO:004867 | regulation of collateral sprouting | 2/85 | 23/28564 | 0.002125928 | 0.002125928 | 0.049084683 | Bdnf/Omg | 2 |
| GO:005080 | BP | GO:005080 | regulation of synapse organization | 5/85 | 305/2856 | 0.002195404 | 0.002195404 | 0.049084683 | Arc/Bdnf/Pcdh8/Nedd9/Sparc | 5 |
| GO:004358 | BP | GO:004358 | tongue development | 2/85 | 24/28564 | 0.002314722 | 0.002314722 | 0.049084683 | Bdnf/Wnt10a | 2 |
| GO:004669 | BP | GO:004669 | decidualization | 2/85 | 24/28564 | 0.002314722 | 0.002314722 | 0.049084683 | Ptgs2/Junb | 2 |
| GO:006008 | BP | GO:006008 | inhibitory postsynaptic potential | 2/85 | 24/28564 | 0.002314722 | 0.002314722 | 0.049084683 | Bdnf/Npas4 | 2 |
| GO:005080 | BP | GO:005080 | regulation of synapse structure or activity | 5/85 | 313/2856 | 0.002455225 | 0.002455225 | 0.051237697 | Arc/Bdnf/Pcdh8/Nedd9/Sparc | 5 |
| GO:003505 | BP | GO:003505 | cardiocyte differentiation | 4/85 | 196/2856 | 0.002822887 | 0.002822887 | 0.054578723 | Sema3c/Inhba/Rgs2/Trip10 | 4 |
| GO:007072 | BP | GO:007072 | response to cholesterol | 2/85 | 27/28564 | 0.002926732 | 0.002926732 | 0.054969756 | Inhba/Mlc1 | 2 |
| GO:009873 | BP | GO:009873 | import across plasma membrane | 4/85 | 208/2856 | 0.003491628 | 0.003491628 | 0.061185794 | Kcnj3/Rgs2/Slc2a1/Gfap | 4 |
| GO:005115 | BP | GO:005115 | regulation of striated muscle cell differentiation | 3/85 | 106/2856 | 0.003934611 | 0.003934611 | 0.064110778 | Rgs2/Bdnf/Trip10 | 3 |
| GO:000193 | BP | GO:000193 | negative regulation of protein phosphorylation | 5/85 | 354/2856 | 0.004149464 | 0.004149464 | 0.0641818 | Pmepa1/Rgs2/Smpd1/Gadd45b/Pebp1 | 5 |
| GO:003526 | BP | GO:003526 | organ growth | 4/85 | 220/2856 | 0.004260856 | 0.004260856 | 0.064987903 | Rgs2/Gja1/Trip10/Col27a1 | 4 |
| GO:000821 | BP | GO:000821 | regulation of blood pressure | 4/85 | 222/2856 | 0.004399301 | 0.004399301 | 0.064987903 | Ptgs2/Ptgs1/Gja1/Gpr37l1 | 4 |
| GO:006105 | BP | GO:006105 | regulation of cell growth involved in cardiac muscle cell developr | 2/85 | 34/28564 | 0.004615083 | 0.004615083 | 0.067418001 | Rgs2/Trip10 | 2 |
| GO:000206 | BP | GO:000206 | chondrocyte development | 2/85 | 35/28564 | 0.004885369 | 0.004885369 | 0.067610258 | Col27a1/Acan | 2 |
| GO:001404 | BP | GO:001404 | glutamate secretion | 2/85 | 35/28564 | 0.004885369 | 0.004885369 | 0.067610258 | Bdnf/Gja1 | 2 |
| GO:004290 | BP | GO:004290 | xenobiotic transport | 2/85 | 36/28564 | 0.005162795 | 0.005162795 | 0.069976456 | Slc2a1/Gja1 | 2 |
| GO:000194 | BP | GO:000194 | hair follicle development | 3/85 | 118/2856 | 0.00530655 | 0.00530655 | 0.071190986 | Inhba/Ptgs2/Wnt10a | 3 |
| GO:002240 | BP | GO:002240 | molting cycle process | 3/85 | 121/2856 | 0.005688934 | 0.005688934 | 0.073253703 | Inhba/Ptgs2/Wnt10a | 3 |
| GO:002240 | BP | GO:002240 | hair cycle process | 3/85 | 121/2856 | 0.005688934 | 0.005688934 | 0.073253703 | Inhba/Ptgs2/Wnt10a | 3 |
| GO:005195 | BP | GO:005195 | negative regulation of amine transport | 2/85 | 38/28564 | 0.005738891 | 0.005738891 | 0.073253703 | Rgs2/Ptgs1 | 2 |
| GO:200017 | BP | GO:200017 | regulation of neural precursor cell proliferation | 3/85 | 126/2856 | 0.006362193 | 0.006362193 | 0.078173921 | Bdnf/Emx1/Gpr37l1 | 3 |
| GO:003817 | BP | GO:003817 | neurotrophin signaling pathway | 2/85 | 41/28564 | 0.006655496 | 0.006655496 | 0.081020609 | Bdnf/Ndn | 2 |
| GO:003254 | BP | GO:003254 | mitochondrial translation | 3/85 | 132/2856 | 0.007230391 | 0.007230391 | 0.083469061 | Mrpl41/Mrps2/Mrpl49 | 3 |
| GO:004232 | BP | GO:004232 | negative regulation of phosphorylation | 5/85 | 405/2856 | 0.00725961 | 0.00725961 | 0.083469061 | Pmepa1/Rgs2/Smpd1/Gadd45b/Pebp1 | 5 |
| GO:004222 | BP | GO:004222 | response to cocaine | 2/85 | 44/28564 | 0.00763402 | 0.00763402 | 0.083951624 | Bdnf/Smpd1 | 2 |
| GO:006034 | BP | GO:006034 | bone development | 4/85 | 262/2856 | 0.007838081 | 0.007838081 | 0.083951624 | Gja1/Sparc/Phospho1/Col27a1 | 4 |
| GO:000979 | BP | GO:000979 | post-embryonic development | 3/85 | 136/2856 | 0.007846311 | 0.007846311 | 0.083951624 | Sema3c/Emx1/Ndn | 3 |
| GO:001701 | BP | GO:001701 | regulation of transforming growth factor beta receptor signaling pa | 3/85 | 136/2856 | 0.007846311 | 0.007846311 | 0.083951624 | Pmepa1/Tgfb1i1/Zfp703 | 3 |
| GO:009865 | BP | GO:009865 | import into cell | 4/85 | 264/2856 | 0.008045644 | 0.008045644 | 0.083951624 | Kcnj3/Rgs2/Slc2a1/Gfap | 4 |
| GO:000193 | BP | GO:000193 | negative regulation of endothelial cell proliferation | 2/85 | 46/28564 | 0.008320202 | 0.008320202 | 0.084876795 | Gja1/Sparc | 2 |
| GO:003355 | BP | GO:003355 | unsaturated fatty acid metabolic process | 3/85 | 139/2856 | 0.008327983 | 0.008327983 | 0.084876795 | Ptgs2/Scd1/Ptgs1 | 3 |
| GO:000329 | BP | GO:000329 | physiological muscle hypertrophy | 2/85 | 48/28564 | 0.009033056 | 0.009033056 | 0.086103552 | Rgs2/Trip10 | 2 |
| GO:000803 | BP | GO:000803 | neuron recognition | 2/85 | 48/28564 | 0.009033056 | 0.009033056 | 0.086103552 | Bdnf/Ndn | 2 |
| GO:007138 | BP | GO:007138 | cellular response to glucocorticoid stimulus | 2/85 | 49/28564 | 0.00939938 | 0.00939938 | 0.087643346 | Npas4/Sstr2 | 2 |
| GO:006013 | BP | GO:006013 | maternal process involved in female pregnancy | 2/85 | 50/28564 | 0.009772249 | 0.009772249 | 0.090048275 | Ptgs2/Junb | 2 |
| GO:001923 | BP | GO:001923 | sensory perception of pain | 3/85 | 152/2856 | 0.010613979 | 0.010613979 | 0.094287769 | Ptgs2/Ptgs1/Ndn | 3 |
| GO:009730 | BP | GO:009730 | cellular response to alcohol | 3/85 | 156/2856 | 0.011383292 | 0.011383292 | 0.099112807 | Inhba/Npas4/Mlc1 | 3 |
| GO:000166 | BP | GO:000166 | behavioral fear response | 2/85 | 56/28564 | 0.012144569 | 0.012144569 | 0.101839985 | Bdnf/Gja1 | 2 |
| GO:190186 | BP | GO:190186 | regulation of muscle tissue development | 2/85 | 56/28564 | 0.012144569 | 0.012144569 | 0.101839985 | Bdnf/Gja1 | 2 |
| GO:007169 | BP | GO:007169 | anatomical structure maturation | 4/85 | 305/2856 | 0.013124312 | 0.013124312 | 0.105858994 | Ptgs2/Gja1/Phospho1/Pebp1 | 4 |
| GO:014035 | BP | GO:014035 | lipid export from cell | 2/85 | 59/28564 | 0.013415765 | 0.013415765 | 0.107550001 | Inhba/Ptgs2 | 2 |
| GO:000803 | BP | GO:000803 | cell recognition | 3/85 | 167/2856 | 0.0136616 | 0.0136616 | 0.108201257 | Bdnf/Nedd9/Ndn | 3 |
| GO:014005 | BP | GO:014005 | mitochondrial gene expression | 3/85 | 167/2856 | 0.0136616 | 0.0136616 | 0.108201257 | Mrpl41/Mrps2/Mrpl49 | 3 |
| GO:003432 | BP | GO:003432 | cell junction assembly | 5/85 | 477/2856 | 0.014004115 | 0.014004115 | 0.109593605 | Bdnf/Npas4/Gja1/Hapln4/Zfp703 | 5 |
| GO:000197 | BP | GO:000197 | blood vessel remodeling | 2/85 | 61/28564 | 0.01429399 | 0.01429399 | 0.110003247 | Sema3c/Gja1 | 2 |
| GO:003019 | BP | GO:003019 | collagen fibril organization | 2/85 | 61/28564 | 0.01429399 | 0.01429399 | 0.110003247 | Col27a1/Acan | 2 |
| GO:004259 | BP | GO:004259 | fear response | 2/85 | 62/28564 | 0.014742213 | 0.014742213 | 0.111391553 | Bdnf/Gja1 | 2 |
| GO:005076 | BP | GO:005076 | negative regulation of neurogenesis | 3/85 | 173/2856 | 0.0150059 | 0.0150059 | 0.112243491 | Sema3c/Bdnf/Gpr37l1 | 3 |
| GO:003110 | BP | GO:003110 | neuron projection regeneration | 2/85 | 63/28564 | 0.015196457 | 0.015196457 | 0.112243491 | Omg/Gfap | 2 |
| GO:005178 | BP | GO:005178 | positive regulation of nuclear division | 2/85 | 65/28564 | 0.016122853 | 0.016122853 | 0.115621067 | Gja1/Pebp1 | 2 |
| GO:005196 | BP | GO:005196 | negative regulation of nervous system development | 3/85 | 178/2856 | 0.016181395 | 0.016181395 | 0.115621067 | Sema3c/Bdnf/Gpr37l1 | 3 |
| GO:004340 | BP | GO:004340 | negative regulation of MAPK cascade | 3/85 | 187/2856 | 0.018424581 | 0.018424581 | 0.125655721 | Rgs2/Smpd1/Pebp1 | 3 |
| GO:001633 | BP | GO:001633 | morphogenesis of embryonic epithelium | 3/85 | 188/2856 | 0.018683961 | 0.018683961 | 0.125971755 | Tgfb1i1/Pcdh8/Wnt9b | 3 |
| GO:009960 | BP | GO:009960 | regulation of neurotransmitter receptor activity | 2/85 | 72/28564 | 0.019549245 | 0.019549245 | 0.131133228 | Nptx2/Arc | 2 |
| GO:000165 | BP | GO:000165 | temperature homeostasis | 3/85 | 192/2856 | 0.019741804 | 0.019741804 | 0.131752674 | Ptgs2/Scd1/Gja1 | 3 |
| GO:007121 | BP | GO:007121 | cellular response to abiotic stimulus | 4/85 | 349/2856 | 0.020460484 | 0.020460484 | 0.135176643 | Ptgs2/Smpd1/Gja1/Gpr88 | 4 |
| GO:010400 | BP | GO:010400 | cellular response to environmental stimulus | 4/85 | 349/2856 | 0.020460484 | 0.020460484 | 0.135176643 | Ptgs2/Smpd1/Gja1/Gpr88 | 4 |
| GO:000175 | BP | GO:000175 | somitogenesis | 2/85 | 77/28564 | 0.022166624 | 0.022166624 | 0.142859204 | Sema3c/Pcdh8 | 2 |
| GO:000749 | BP | GO:000749 | endoderm development | 2/85 | 77/28564 | 0.022166624 | 0.022166624 | 0.142859204 | Arc/Inhba | 2 |
| GO:004340 | BP | GO:004340 | regulation of MAP kinase activity | 3/85 | 202/2856 | 0.022528899 | 0.022528899 | 0.143784336 | Rgs2/Smpd1/Pik3r6 | 3 |
| GO:004649 | BP | GO:004649 | nicotinamide nucleotide metabolic process | 2/85 | 79/28564 | 0.023252021 | 0.023252021 | 0.146972541 | Aldh1l1/Ldha | 2 |
| GO:007127 | BP | GO:007127 | cellular response to calcium ion | 2/85 | 81/28564 | 0.024358949 | 0.024358949 | 0.151064187 | Junb/Smpd1 | 2 |
| GO:006135 | BP | GO:006135 | neural precursor cell proliferation | 3/85 | 212/2856 | 0.025519759 | 0.025519759 | 0.15678396 | Bdnf/Emx1/Gpr37l1 | 3 |
| GO:002170 | BP | GO:002170 | developmental maturation | 4/85 | 381/2856 | 0.027117601 | 0.027117601 | 0.165057914 | Ptgs2/Gja1/Phospho1/Pebp1 | 4 |
| GO:000836 | BP | GO:000836 | regulation of cell size | 3/85 | 218/2856 | 0.027412007 | 0.027412007 | 0.16534795 | Sema3c/Bdnf/Omg | 3 |
| GO:199054 | BP | GO:199054 | mitochondrial transmembrane transport | 2/85 | 88/28564 | 0.028398625 | 0.028398625 | 0.16534795 | Slc41a3/Dnlz | 2 |
| GO:000854 | BP | GO:000854 | epidermis development | 4/85 | 388/2856 | 0.028727649 | 0.028727649 | 0.16534795 | Inhba/Ptgs2/Ptgs1/Wnt10a | 4 |
| GO:000325 | BP | GO:000325 | cardiac neural crest cell migration involved in outflow tract morph | 1/85 | 10/28564 | 0.029366962 | 0.029366962 | 0.16534795 | Sema3c | 1 |
| GO:001049 | BP | GO:001049 | intercellular transport | 1/85 | 10/28564 | 0.029366962 | 0.029366962 | 0.16534795 | Arc | 1 |
| GO:004666 | BP | GO:004666 | retinal cell programmed cell death | 1/85 | 10/28564 | 0.029366962 | 0.029366962 | 0.16534795 | Bdnf | 1 |
| GO:004688 | BP | GO:004688 | follicle-stimulating hormone secretion | 1/85 | 10/28564 | 0.029366962 | 0.029366962 | 0.16534795 | Inhba | 1 |
| GO:006031 | BP | GO:006031 | regulation of blood vessel remodeling | 1/85 | 10/28564 | 0.029366962 | 0.029366962 | 0.16534795 | Gja1 | 1 |
| GO:007217 | BP | GO:007217 | metanephric tubule morphogenesis | 1/85 | 10/28564 | 0.029366962 | 0.029366962 | 0.16534795 | Wnt9b | 1 |
| GO:009052 | BP | GO:009052 | actin filament reorganization | 1/85 | 10/28564 | 0.029366962 | 0.029366962 | 0.16534795 | Nedd9 | 1 |
| GO:009953 | BP | GO:009953 | synaptic signaling via neuropeptide | 1/85 | 10/28564 | 0.029366962 | 0.029366962 | 0.16534795 | Bdnf | 1 |
| GO:200098 | BP | GO:200098 | positive regulation of behavioral fear response | 1/85 | 10/28564 | 0.029366962 | 0.029366962 | 0.16534795 | Gja1 | 1 |
| GO:005138 | BP | GO:005138 | response to glucocorticoid | 2/85 | 90/28564 | 0.02959892 | 0.02959892 | 0.16534795 | Npas4/Sstr2 | 2 |
| GO:000341 | BP | GO:000341 | growth plate cartilage chondrocyte differentiation | 1/85 | 11/28564 | 0.032256358 | 0.032256358 | 0.16534795 | Col27a1 | 1 |
| GO:002194 | BP | GO:002194 | positive regulation of cerebellar granule cell precursor proliferatio | 1/85 | 11/28564 | 0.032256358 | 0.032256358 | 0.16534795 | Gpr37l1 | 1 |
| GO:003162 | BP | GO:003162 | regulation of fever generation | 1/85 | 11/28564 | 0.032256358 | 0.032256358 | 0.16534795 | Ptgs2 | 1 |
| GO:006017 | BP | GO:006017 | limb bud formation | 1/85 | 11/28564 | 0.032256358 | 0.032256358 | 0.16534795 | Sema3c | 1 |
| GO:006060 | BP | GO:006060 | dichotomous subdivision of an epithelial terminal unit | 1/85 | 11/28564 | 0.032256358 | 0.032256358 | 0.16534795 | Sema3c | 1 |
| GO:007137 | BP | GO:007137 | cellular response to follicle-stimulating hormone stimulus | 1/85 | 11/28564 | 0.032256358 | 0.032256358 | 0.16534795 | Inhba | 1 |
| GO:009026 | BP | GO:009026 | fibroblast growth factor production | 1/85 | 11/28564 | 0.032256358 | 0.032256358 | 0.16534795 | Ptgs2 | 1 |
| GO:009027 | BP | GO:009027 | regulation of fibroblast growth factor production | 1/85 | 11/28564 | 0.032256358 | 0.032256358 | 0.16534795 | Ptgs2 | 1 |
| GO:006105 | BP | GO:006105 | somite development | 2/85 | 96/28564 | 0.033318594 | 0.033318594 | 0.16534795 | Sema3c/Pcdh8 | 2 |
| GO:000697 | BP | GO:000697 | response to oxidative stress | 4/85 | 408/2856 | 0.033637177 | 0.033637177 | 0.16534795 | Ptgs2/Ptgs1/Srxn1/Gpr37l1 | 4 |
| GO:000333 | BP | GO:000333 | mesenchymal to epithelial transition involved in metanephros mo | 1/85 | 12/28564 | 0.035137254 | 0.035137254 | 0.16534795 | Wnt9b | 1 |
| GO:000604 | BP | GO:000604 | UDP-N-acetylglucosamine metabolic process | 1/85 | 12/28564 | 0.035137254 | 0.035137254 | 0.16534795 | Uap1l1 | 1 |
| GO:000688 | BP | GO:000688 | intracellular sequestering of iron ion | 1/85 | 12/28564 | 0.035137254 | 0.035137254 | 0.16534795 | Ftl1 | 1 |
| GO:000718 | BP | GO:000718 | SMAD protein complex assembly | 1/85 | 12/28564 | 0.035137254 | 0.035137254 | 0.16534795 | Pmepa1 | 1 |
| GO:001064 | BP | GO:001064 | regulation of cell communication by electrical coupling | 1/85 | 12/28564 | 0.035137254 | 0.035137254 | 0.16534795 | Gja1 | 1 |
| GO:003038 | BP | GO:003038 | fructose 1,6-bisphosphate metabolic process | 1/85 | 12/28564 | 0.035137254 | 0.035137254 | 0.16534795 | Aldoc | 1 |
| GO:003235 | BP | GO:003235 | response to follicle-stimulating hormone | 1/85 | 12/28564 | 0.035137254 | 0.035137254 | 0.16534795 | Inhba | 1 |
| GO:003433 | BP | GO:003433 | adherens junction assembly | 1/85 | 12/28564 | 0.035137254 | 0.035137254 | 0.16534795 | Zfp703 | 1 |
| GO:003543 | BP | GO:003543 | maintenance of protein localization in endoplasmic reticulum | 1/85 | 12/28564 | 0.035137254 | 0.035137254 | 0.16534795 | Gja1 | 1 |
| GO:004824 | BP | GO:004824 | epinephrine transport | 1/85 | 12/28564 | 0.035137254 | 0.035137254 | 0.16534795 | Ptgs1 | 1 |
| GO:006168 | BP | GO:006168 | chaperone-mediated autophagy | 1/85 | 12/28564 | 0.035137254 | 0.035137254 | 0.16534795 | Gfap | 1 |
| GO:007258 | BP | GO:007258 | caveolin-mediated endocytosis | 1/85 | 12/28564 | 0.035137254 | 0.035137254 | 0.16534795 | Mlc1 | 1 |
| GO:010604 | BP | GO:010604 | regulation of cellular response to osmotic stress | 1/85 | 12/28564 | 0.035137254 | 0.035137254 | 0.16534795 | Ptgs2 | 1 |
| GO:006004 | BP | GO:006004 | heart contraction | 3/85 | 242/2856 | 0.03570983 | 0.03570983 | 0.16534795 | Rgs2/Gja1/Pebp1 | 3 |
| GO:012016 | BP | GO:012016 | positive regulation of cold-induced thermogenesis | 2/85 | 100/2856 | 0.035894661 | 0.035894661 | 0.16534795 | Scd1/Gja1 | 2 |
| GO:003150 | BP | GO:003150 | heterochromatin formation | 2/85 | 101/2856 | 0.036550407 | 0.036550407 | 0.16534795 | Baz1a/Ndn | 2 |
| GO:006002 | BP | GO:006002 | roof of mouth development | 2/85 | 101/2856 | 0.036550407 | 0.036550407 | 0.16534795 | Inhba/Wnt9b | 2 |
| GO:007155 | BP | GO:007155 | response to transforming growth factor beta | 3/85 | 246/2856 | 0.037205276 | 0.037205276 | 0.16534795 | Pmepa1/Tgfb1i1/Zfp703 | 3 |
| GO:009730 | BP | GO:009730 | response to alcohol | 3/85 | 246/2856 | 0.037205276 | 0.037205276 | 0.16534795 | Inhba/Npas4/Mlc1 | 3 |
| GO:000215 | BP | GO:000215 | osteoclast proliferation | 1/85 | 13/28564 | 0.038009674 | 0.038009674 | 0.16534795 | Junb | 1 |
| GO:000309 | BP | GO:000309 | regulation of glomerular filtration | 1/85 | 13/28564 | 0.038009674 | 0.038009674 | 0.16534795 | Gja1 | 1 |
| GO:001809 | BP | GO:001809 | protein polyglutamylation | 1/85 | 13/28564 | 0.038009674 | 0.038009674 | 0.16534795 | Ttll6 | 1 |
| GO:003165 | BP | GO:003165 | positive regulation of heat generation | 1/85 | 13/28564 | 0.038009674 | 0.038009674 | 0.16534795 | Ptgs2 | 1 |
| GO:003227 | BP | GO:003227 | regulation of gonadotropin secretion | 1/85 | 13/28564 | 0.038009674 | 0.038009674 | 0.16534795 | Inhba | 1 |
| GO:003324 | BP | GO:003324 | positive regulation of amine metabolic process | 1/85 | 13/28564 | 0.038009674 | 0.038009674 | 0.16534795 | Pebp1 | 1 |
| GO:004455 | BP | GO:004455 | relaxation of smooth muscle | 1/85 | 13/28564 | 0.038009674 | 0.038009674 | 0.16534795 | Rgs2 | 1 |
| GO:000150 | BP | GO:000150 | ossification | 4/85 | 427/2856 | 0.038729366 | 0.038729366 | 0.16534795 | Ptgs2/Junb/Gja1/Phospho1 | 4 |
| GO:000165 | BP | GO:000165 | ureteric bud development | 2/85 | 107/2856 | 0.040580897 | 0.040580897 | 0.16534795 | Bdnf/Wnt9b | 2 |
| GO:005079 | BP | GO:005079 | regulation of behavior | 2/85 | 107/2856 | 0.040580897 | 0.040580897 | 0.16534795 | Gja1/Mchr1 | 2 |
| GO:000166 | BP | GO:000166 | fever generation | 1/85 | 14/28564 | 0.040873643 | 0.040873643 | 0.16534795 | Ptgs2 | 1 |
| GO:000206 | BP | GO:000206 | columnar/cuboidal epithelial cell maturation | 1/85 | 14/28564 | 0.040873643 | 0.040873643 | 0.16534795 | Gja1 | 1 |
| GO:004248 | BP | GO:004248 | regulation of odontogenesis of dentin-containing tooth | 1/85 | 14/28564 | 0.040873643 | 0.040873643 | 0.16534795 | Wnt10a | 1 |
| GO:004659 | BP | GO:004659 | positive regulation of viral entry into host cell | 1/85 | 14/28564 | 0.040873643 | 0.040873643 | 0.16534795 | Smpd1 | 1 |
| GO:007217 | BP | GO:007217 | mesonephric tubule formation | 1/85 | 14/28564 | 0.040873643 | 0.040873643 | 0.16534795 | Wnt9b | 1 |
| GO:007529 | BP | GO:007529 | positive regulation by symbiont of entry into host | 1/85 | 14/28564 | 0.040873643 | 0.040873643 | 0.16534795 | Smpd1 | 1 |
| GO:009891 | BP | GO:009891 | retrograde trans-synaptic signaling | 1/85 | 14/28564 | 0.040873643 | 0.040873643 | 0.16534795 | Bdnf | 1 |
| GO:190494 | BP | GO:190494 | midbrain dopaminergic neuron differentiation | 1/85 | 14/28564 | 0.040873643 | 0.040873643 | 0.16534795 | Wnt9b | 1 |
| GO:200124 | BP | GO:200124 | positive regulation of extrinsic apoptotic signaling pathway in abs | 1/85 | 14/28564 | 0.040873643 | 0.040873643 | 0.16534795 | Inhba | 1 |
| GO:009706 | BP | GO:009706 | dendritic spine organization | 2/85 | 108/2856 | 0.041268346 | 0.041268346 | 0.165765842 | Arc/Nedd9 | 2 |
| GO:000715 | BP | GO:000715 | homophilic cell adhesion via plasma membrane adhesion molec | 2/85 | 109/2856 | 0.041960197 | 0.041960197 | 0.165765842 | Pcdh8/Pcdhgc3 | 2 |
| GO:200024 | BP | GO:200024 | positive regulation of reproductive process | 2/85 | 110/2856 | 0.042656416 | 0.042656416 | 0.165765842 | Inhba/Gja1 | 2 |
| GO:001059 | BP | GO:001059 | positive regulation of endothelial cell migration | 2/85 | 111/2856 | 0.043356974 | 0.043356974 | 0.165765842 | Ptgs2/Sparc | 2 |
| GO:007082 | BP | GO:007082 | heterochromatin organization | 2/85 | 111/2856 | 0.043356974 | 0.043356974 | 0.165765842 | Baz1a/Ndn | 2 |
| GO:001567 | BP | GO:001567 | carbon dioxide transport | 1/85 | 15/28564 | 0.043729185 | 0.043729185 | 0.165765842 | Car12 | 1 |
| GO:001581 | BP | GO:001581 | glycine transport | 1/85 | 15/28564 | 0.043729185 | 0.043729185 | 0.165765842 | Rgs2 | 1 |
| GO:004254 | BP | GO:004254 | hemoglobin biosynthetic process | 1/85 | 15/28564 | 0.043729185 | 0.043729185 | 0.165765842 | Inhba | 1 |
| GO:003166 | BP | GO:003166 | cellular response to extracellular stimulus | 3/85 | 264/2856 | 0.044326381 | 0.044326381 | 0.165765842 | Ldha/Slc2a1/Wnt9b | 3 |
| GO:004581 | BP | GO:004581 | negative regulation of gene expression, epigenetic | 2/85 | 113/2856 | 0.044770977 | 0.044770977 | 0.165765842 | Baz1a/Ndn | 2 |
| GO:003134 | BP | GO:003134 | positive regulation of defense response | 4/85 | 452/2856 | 0.046069068 | 0.046069068 | 0.165765842 | Ptgs2/Ptgs2os/Gja1/Tril | 4 |
| GO:005134 | BP | GO:005134 | negative regulation of transferase activity | 3/85 | 269/2856 | 0.04641688 | 0.04641688 | 0.165765842 | Rgs2/Smpd1/Gadd45b | 3 |
| GO:000751 | BP | GO:000751 | adult heart development | 1/85 | 16/28564 | 0.046576326 | 0.046576326 | 0.165765842 | Gja1 | 1 |
| GO:000922 | BP | GO:000922 | nucleotide-sugar biosynthetic process | 1/85 | 16/28564 | 0.046576326 | 0.046576326 | 0.165765842 | Uap1l1 | 1 |
| GO:001569 | BP | GO:001569 | magnesium ion transport | 1/85 | 16/28564 | 0.046576326 | 0.046576326 | 0.165765842 | Slc41a3 | 1 |
| GO:002197 | BP | GO:002197 | telencephalon regionalization | 1/85 | 16/28564 | 0.046576326 | 0.046576326 | 0.165765842 | Emx1 | 1 |
| GO:004364 | BP | GO:004364 | dicarboxylic acid catabolic process | 1/85 | 16/28564 | 0.046576326 | 0.046576326 | 0.165765842 | Aldh1l1 | 1 |
| GO:005511 | BP | GO:005511 | relaxation of cardiac muscle | 1/85 | 16/28564 | 0.046576326 | 0.046576326 | 0.165765842 | Rgs2 | 1 |
| GO:006102 | BP | GO:006102 | eyelid development in camera-type eye | 1/85 | 16/28564 | 0.046576326 | 0.046576326 | 0.165765842 | Inhba | 1 |
| GO:008610 | BP | GO:008610 | G protein-coupled receptor signaling pathway involved in heart p | 1/85 | 16/28564 | 0.046576326 | 0.046576326 | 0.165765842 | Rgs2 | 1 |
| GO:190295 | BP | GO:190295 | regulation of dendritic spine maintenance | 1/85 | 16/28564 | 0.046576326 | 0.046576326 | 0.165765842 | Nedd9 | 1 |
| GO:004563 | BP | GO:004563 | positive regulation of myeloid cell differentiation | 2/85 | 116/2856 | 0.046923738 | 0.046923738 | 0.165765842 | Inhba/Nedd9 | 2 |
| GO:007148 | BP | GO:007148 | cellular response to light stimulus | 2/85 | 117/2856 | 0.04764967 | 0.04764967 | 0.165765842 | Smpd1/Gpr88 | 2 |
| GO:010602 | BP | GO:010602 | neuron projection organization | 2/85 | 117/2856 | 0.04764967 | 0.04764967 | 0.165765842 | Arc/Nedd9 | 2 |
| GO:001605 | BP | GO:001605 | Wnt signaling pathway | 4/85 | 458/2856 | 0.047938796 | 0.047938796 | 0.165765842 | Tgfb1i1/Wnt10a/Zfp703/Wnt9b | 4 |
| GO:001095 | BP | GO:001095 | regulation of metal ion transport | 4/85 | 460/2856 | 0.048571333 | 0.048571333 | 0.165765842 | Ptgs2/Gja1/Kcns2/Mchr1 | 4 |
| GO:019873 | BP | GO:019873 | cell-cell signaling by wnt | 4/85 | 460/2856 | 0.048571333 | 0.048571333 | 0.165765842 | Tgfb1i1/Wnt10a/Zfp703/Wnt9b | 4 |
| GO:000177 | BP | GO:000177 | immunological synapse formation | 1/85 | 17/28564 | 0.049415089 | 0.049415089 | 0.165765842 | Nedd9 | 1 |
| GO:000335 | BP | GO:000335 | positive regulation of cilium movement | 1/85 | 17/28564 | 0.049415089 | 0.049415089 | 0.165765842 | Ttll6 | 1 |
| GO:000335 | BP | GO:000335 | regulation of cilium beat frequency | 1/85 | 17/28564 | 0.049415089 | 0.049415089 | 0.165765842 | Ttll6 | 1 |
| GO:001586 | BP | GO:001586 | ATP transport | 1/85 | 17/28564 | 0.049415089 | 0.049415089 | 0.165765842 | Gja1 | 1 |
| GO:005138 | BP | GO:005138 | kinetochore assembly | 1/85 | 17/28564 | 0.049415089 | 0.049415089 | 0.165765842 | Cenpa | 1 |
| GO:007149 | BP | GO:007149 | cellular response to fluid shear stress | 1/85 | 17/28564 | 0.049415089 | 0.049415089 | 0.165765842 | Ptgs2 | 1 |
| GO:200069 | BP | GO:200069 | regulation of epithelial cell differentiation involved in kidney dev | 1/85 | 17/28564 | 0.049415089 | 0.049415089 | 0.165765842 | Wnt9b | 1 |
| GO:009879 | CC | GO:009879 | mitochondrial protein-containing complex | 6/84 | 300/2858 | 0.000260131 | 0.000260131 | 0.023240321 | Mrpl41/Cox5b/Mrps2/Ndufa2/Uqcr11/Mrpl49 | 6 |
| GO:009953 | CC | GO:009953 | synapse-associated extracellular matrix | 2/84 | 10/28585 | 0.000378149 | 0.000378149 | 0.023240321 | Hapln4/Acan | 2 |
| GO:004320 | CC | GO:004320 | myelin sheath | 5/84 | 218/2858 | 0.000466443 | 0.000466443 | 0.023240321 | Omg/Cox5b/Tuba1b/Gfap/Pebp1 | 5 |
| GO:004512 | CC | GO:004512 | membrane raft | 6/84 | 390/2858 | 0.001030607 | 0.001030607 | 0.026014447 | Arc/Ptgs2/Slc2a1/Mlc1/Gja1/Tuba1b | 6 |
| GO:009744 | CC | GO:009744 | astrocyte projection | 2/84 | 22/28585 | 0.001897203 | 0.001897203 | 0.034013321 | Mlc1/Gfap | 2 |
| GO:000574 | CC | GO:000574 | mitochondrial respirasome | 3/84 | 90/28585 | 0.002392082 | 0.002392082 | 0.034013321 | Cox5b/Ndufa2/Uqcr11 | 3 |
| GO:009880 | CC | GO:009880 | respiratory chain complex | 3/84 | 91/28585 | 0.002468426 | 0.002468426 | 0.034013321 | Cox5b/Ndufa2/Uqcr11 | 3 |
| GO:000031 | CC | GO:000031 | organellar ribosome | 3/84 | 93/28585 | 0.002625606 | 0.002625606 | 0.034013321 | Mrpl41/Mrps2/Mrpl49 | 3 |
| GO:000576 | CC | GO:000576 | mitochondrial ribosome | 3/84 | 93/28585 | 0.002625606 | 0.002625606 | 0.034013321 | Mrpl41/Mrps2/Mrpl49 | 3 |
| GO:000590 | CC | GO:000590 | caveola | 3/84 | 97/28585 | 0.002958201 | 0.002958201 | 0.034013321 | Ptgs2/Slc2a1/Mlc1 | 3 |
| GO:007046 | CC | GO:007046 | respirasome | 3/84 | 97/28585 | 0.002958201 | 0.002958201 | 0.034013321 | Cox5b/Ndufa2/Uqcr11 | 3 |
| GO:000572 | CC | GO:000572 | pericentric heterochromatin | 2/84 | 33/28585 | 0.004246581 | 0.004246581 | 0.045339437 | Baz1a/Cenpa | 2 |
| GO:199020 | CC | GO:199020 | oxidoreductase complex | 3/84 | 123/2858 | 0.005748964 | 0.005748964 | 0.055044764 | Ldha/Ndufa2/Uqcr11 | 3 |
| GO:007006 | CC | GO:007006 | cytochrome complex | 2/84 | 39/28585 | 0.005892116 | 0.005892116 | 0.055044764 | Cox5b/Uqcr11 | 2 |
| GO:009773 | CC | GO:009773 | non-motile cilium | 3/84 | 168/2858 | 0.01342149 | 0.01342149 | 0.091189072 | Ptgs1/Mchr1/Ttll6 | 3 |
| GO:001470 | CC | GO:001470 | intercalated disc | 2/84 | 67/28585 | 0.016673122 | 0.016673122 | 0.108356214 | Slc2a1/Gja1 | 2 |
| GO:000807 | CC | GO:000807 | voltage-gated potassium channel complex | 2/84 | 77/28585 | 0.021652804 | 0.021652804 | 0.129095169 | Kcnj3/Kcns2 | 2 |
| GO:001986 | CC | GO:001986 | outer membrane | 3/84 | 216/2858 | 0.025920338 | 0.025920338 | 0.129095169 | Ptgs2/Gja1/Pebp1 | 3 |
| GO:003196 | CC | GO:003196 | organelle outer membrane | 3/84 | 216/2858 | 0.025920338 | 0.025920338 | 0.129095169 | Ptgs2/Gja1/Pebp1 | 3 |
| GO:190249 | CC | GO:190249 | transmembrane transporter complex | 4/84 | 392/2858 | 0.028509006 | 0.028509006 | 0.129095169 | Kcnj3/Kcns2/Ndufa2/Uqcr11 | 4 |
| GO:000193 | CC | GO:000193 | female pronucleus | 1/84 | 10/28585 | 0.029004985 | 0.029004985 | 0.129095169 | Slc2a1 | 1 |
| GO:001632 | CC | GO:001632 | apical plasma membrane | 4/84 | 406/2858 | 0.031838526 | 0.031838526 | 0.129095169 | Slc2a1/Car12/Mlc1/Gja1 | 4 |
| GO:199035 | CC | GO:199035 | transporter complex | 4/84 | 415/2858 | 0.034095095 | 0.034095095 | 0.129095169 | Kcnj3/Kcns2/Ndufa2/Uqcr11 | 4 |
| GO:000558 | CC | GO:000558 | fibrillar collagen trimer | 1/84 | 12/28585 | 0.03470543 | 0.03470543 | 0.129095169 | Col27a1 | 1 |
| GO:003601 | CC | GO:003601 | endolysosome | 1/84 | 12/28585 | 0.03470543 | 0.03470543 | 0.129095169 | Smpd1 | 1 |
| GO:009857 | CC | GO:009857 | cytoplasmic side of lysosomal membrane | 1/84 | 12/28585 | 0.03470543 | 0.03470543 | 0.129095169 | Gfap | 1 |
| GO:009864 | CC | GO:009864 | banded collagen fibril | 1/84 | 12/28585 | 0.03470543 | 0.03470543 | 0.129095169 | Col27a1 | 1 |
| GO:001636 | CC | GO:001636 | nuclear matrix | 2/84 | 107/2858 | 0.039668491 | 0.039668491 | 0.129095169 | Sparc/Zfp703 | 2 |
| GO:003101 | CC | GO:003101 | ISWI-type complex | 1/84 | 14/28585 | 0.040372807 | 0.040372807 | 0.129095169 | Baz1a | 1 |
| GO:003568 | CC | GO:003568 | sperm fibrous sheath | 1/84 | 14/28585 | 0.040372807 | 0.040372807 | 0.129095169 | Ldha | 1 |
| GO:004475 | CC | GO:004475 | autolysosome | 1/84 | 14/28585 | 0.040372807 | 0.040372807 | 0.129095169 | Ftl1 | 1 |
| GO:001632 | CC | GO:001632 | basolateral plasma membrane | 3/84 | 258/2858 | 0.040592249 | 0.040592249 | 0.129095169 | Slc2a1/Car12/Mlc1 | 3 |
| GO:004259 | CC | GO:004259 | lamellar body | 1/84 | 16/28585 | 0.046007303 | 0.046007303 | 0.137537621 | Smpd1 | 1 |
| GO:000554 | MF | GO:000554 | hyaluronic acid binding | 2/85 | 24/28171 | 0.002378346 | 0.002378346 | 0.105084521 | Hapln4/Acan | 2 |
| GO:001620 | MF | GO:001620 | antioxidant activity | 3/85 | 88/28171 | 0.002418167 | 0.002418167 | 0.105084521 | Ptgs2/Ptgs1/Srxn1 | 3 |
| GO:001670 | MF | GO:001670 | oxidoreductase activity, acting on single donors with incorporatior | 2/85 | 26/28171 | 0.002789651 | 0.002789651 | 0.105084521 | Ptgs2/Ptgs1 | 2 |
| GO:003361 | MF | GO:003361 | receptor serine/threonine kinase binding | 2/85 | 27/28171 | 0.003006937 | 0.003006937 | 0.105084521 | Inhba/Pebp1 | 2 |
| GO:001545 | MF | GO:001545 | oxidoreduction-driven active transmembrane transporter activity | 2/85 | 40/28171 | 0.006514586 | 0.006514586 | 0.139073391 | Cox5b/Ndufa2 | 2 |
| GO:199078 | MF | GO:199078 | protein tyrosine kinase binding | 3/85 | 127/2817 | 0.006753371 | 0.006753371 | 0.139073391 | Nedd9/Blnk/Gja1 | 3 |
| GO:004292 | MF | GO:004292 | neuropeptide binding | 2/85 | 41/28171 | 0.006835327 | 0.006835327 | 0.139073391 | Sstr2/Mchr1 | 2 |
| GO:000510 | MF | GO:000510 | frizzled binding | 2/85 | 42/28171 | 0.007163117 | 0.007163117 | 0.139073391 | Wnt10a/Wnt9b | 2 |
| GO:004848 | MF | GO:004848 | beta-tubulin binding | 2/85 | 45/28171 | 0.008188336 | 0.008188336 | 0.14308039 | Rgs2/Gja1 | 2 |
| GO:000520 | MF | GO:000520 | extracellular matrix structural constituent | 3/85 | 148/2817 | 0.010250959 | 0.010250959 | 0.153001984 | Sparc/Hapln4/Col27a1 | 3 |
| GO:000905 | MF | GO:000905 | electron transfer activity | 2/85 | 56/28171 | 0.012467753 | 0.012467753 | 0.153001984 | Cox5b/Ndufa2 | 2 |
| GO:000373 | MF | GO:000373 | structural constituent of ribosome | 3/85 | 167/2817 | 0.014172664 | 0.014172664 | 0.153001984 | Mrpl41/Mrps2/Mrpl49 | 3 |
| GO:000460 | MF | GO:000460 | peroxidase activity | 2/85 | 60/28171 | 0.014218942 | 0.014218942 | 0.153001984 | Ptgs2/Ptgs1 | 2 |
| GO:000520 | MF | GO:000520 | structural constituent of cytoskeleton | 2/85 | 72/28171 | 0.020061023 | 0.020061023 | 0.153001984 | Tuba1b/Gfap | 2 |
| GO:003559 | MF | GO:003559 | signaling adaptor activity | 2/85 | 74/28171 | 0.02111718 | 0.02111718 | 0.153001984 | Blnk/Spata2 | 2 |
| GO:001682 | MF | GO:001682 | lyase activity | 3/85 | 195/2817 | 0.021308394 | 0.021308394 | 0.153001984 | Aldoc/Car12/Etnppl | 3 |
| GO:004633 | MF | GO:004633 | SMAD binding | 2/85 | 85/28171 | 0.027324268 | 0.027324268 | 0.153001984 | Pmepa1/Tgfb1i1 | 2 |
| GO:004256 | MF | GO:004256 | hormone binding | 2/85 | 87/28171 | 0.028522803 | 0.028522803 | 0.153001984 | Inhba/Mchr1 | 2 |
| GO:000524 | MF | GO:000524 | voltage-gated potassium channel activity | 2/85 | 89/28171 | 0.029742135 | 0.029742135 | 0.153001984 | Kcnj3/Kcns2 | 2 |
| GO:000802 | MF | GO:000802 | G protein-coupled photoreceptor activity | 1/85 | 10/28171 | 0.029771162 | 0.029771162 | 0.153001984 | Gpr88 | 1 |
| GO:001674 | MF | GO:001674 | hydroxymethyl-, formyl- and related transferase activity | 1/85 | 10/28171 | 0.029771162 | 0.029771162 | 0.153001984 | Aldh1l1 | 1 |
| GO:007073 | MF | GO:007073 | protein-glutamic acid ligase activity | 1/85 | 10/28171 | 0.029771162 | 0.029771162 | 0.153001984 | Ttll6 | 1 |
| GO:007074 | MF | GO:007074 | tubulin-glutamic acid ligase activity | 1/85 | 10/28171 | 0.029771162 | 0.029771162 | 0.153001984 | Ttll6 | 1 |
| GO:001670 | MF | GO:001670 | oxidoreductase activity, acting on paired donors, with incorporatic | 3/85 | 230/2817 | 0.032532278 | 0.032532278 | 0.153001984 | Ptgs2/Scd1/Ptgs1 | 3 |
| GO:000849 | MF | GO:000849 | translation activator activity | 1/85 | 11/28171 | 0.032699661 | 0.032699661 | 0.153001984 | Pcbp1 | 1 |
| GO:004228 | MF | GO:004228 | xylosyltransferase activity | 1/85 | 11/28171 | 0.032699661 | 0.032699661 | 0.153001984 | Gxylt2 | 1 |
| GO:004849 | MF | GO:004849 | Roundabout binding | 1/85 | 11/28171 | 0.032699661 | 0.032699661 | 0.153001984 | Tgfb1i1 | 1 |
| GO:000512 | MF | GO:000512 | cytokine activity | 3/85 | 231/2817 | 0.032890459 | 0.032890459 | 0.153001984 | Inhba/Wnt10a/Wnt9b | 3 |
| GO:000988 | MF | GO:000988 | photoreceptor activity | 1/85 | 12/28171 | 0.035619424 | 0.035619424 | 0.153001984 | Gpr88 | 1 |
| GO:003500 | MF | GO:003500 | phosphatidylinositol 3-kinase activity | 1/85 | 12/28171 | 0.035619424 | 0.035619424 | 0.153001984 | Pik3r6 | 1 |
| GO:003970 | MF | GO:003970 | co-receptor binding | 1/85 | 12/28171 | 0.035619424 | 0.035619424 | 0.153001984 | Wnt9b | 1 |
| GO:000412 | MF | GO:000412 | cytochrome-c oxidase activity | 1/85 | 13/28171 | 0.038530477 | 0.038530477 | 0.153001984 | Cox5b | 1 |
| GO:001683 | MF | GO:001683 | aldehyde-lyase activity | 1/85 | 13/28171 | 0.038530477 | 0.038530477 | 0.153001984 | Aldoc | 1 |
| GO:001667 | MF | GO:001667 | oxidoreductase activity, acting on a heme group of donors | 1/85 | 14/28171 | 0.041432846 | 0.041432846 | 0.153001984 | Cox5b | 1 |
| GO:000408 | MF | GO:000408 | carbonate dehydratase activity | 1/85 | 15/28171 | 0.044326557 | 0.044326557 | 0.153001984 | Car12 | 1 |
| GO:001085 | MF | GO:001085 | cyclase regulator activity | 1/85 | 15/28171 | 0.044326557 | 0.044326557 | 0.153001984 | Rgs2 | 1 |
| GO:004693 | MF | GO:004693 | 1-phosphatidylinositol-3-kinase regulator activity | 1/85 | 15/28171 | 0.044326557 | 0.044326557 | 0.153001984 | Pik3r6 | 1 |
| GO:009884 | MF | GO:009884 | sequence-specific single stranded DNA binding | 1/85 | 15/28171 | 0.044326557 | 0.044326557 | 0.153001984 | Pcbp1 | 1 |
| GO:000516 | MF | GO:000516 | neurotrophin receptor binding | 1/85 | 16/28171 | 0.047211635 | 0.047211635 | 0.153001984 | Bdnf | 1 |
| GO:000524 | MF | GO:000524 | gap junction channel activity | 1/85 | 16/28171 | 0.047211635 | 0.047211635 | 0.153001984 | Gja1 | 1 |
| GO:001664 | MF | GO:001664 | oxidoreductase activity, acting on the CH-NH group of donors, NA | 1/85 | 16/28171 | 0.047211635 | 0.047211635 | 0.153001984 | Aldh1l1 | 1 |

Cre+_vs_Cre-_enrichGO

| | ONTOLOGY | ID | Description | GeneRatio | BgRatio | pvalue | p.adjust | qvalue | geneID | Count |
|---|---|---|---|---|---|---|---|---|---|---|
| GO:1990573 | BP | GO:1990573 | potassium ion import across plasma membrane | 2/20 | 48/28564 | 0.000515323 | 0.000515323 | 0.057381221 | Kcnj3/Atp1b2 | 2 |
| GO:0002040 | BP | GO:0002040 | sprouting angiogenesis | 2/20 | 128/28564 | 0.003590769 | 0.003590769 | 0.057381221 | Cdh13/Rspo3 | 2 |
| GO:0061418 | BP | GO:0061418 | regulation of transcription from RNA polymerase II promoter in response to hypoxia | 1/20 | 10/28564 | 0.006980897 | 0.006980897 | 0.057381221 | Nfe2l2 | 1 |
| GO:0060352 | BP | GO:0060352 | cell adhesion molecule production | 1/20 | 11/28564 | 0.007676434 | 0.007676434 | 0.057381221 | Gcnt1 | 1 |
| GO:0070257 | BP | GO:0070257 | positive regulation of mucus secretion | 1/20 | 11/28564 | 0.007676434 | 0.007676434 | 0.057381221 | Sytl2 | 1 |
| GO:0009404 | BP | GO:0009404 | toxin metabolic process | 1/20 | 12/28564 | 0.008371509 | 0.008371509 | 0.057381221 | Nfe2l2 | 1 |
| GO:2000849 | BP | GO:2000849 | regulation of glucocorticoid secretion | 1/20 | 13/28564 | 0.009066121 | 0.009066121 | 0.057381221 | Galr1 | 1 |
| GO:0098739 | BP | GO:0098739 | import across plasma membrane | 2/20 | 208/28564 | 0.009197564 | 0.009197564 | 0.057381221 | Kcnj3/Atp1b2 | 2 |
| GO:0098742 | BP | GO:0098742 | cell-cell adhesion via plasma-membrane adhesion molecules | 2/20 | 211/28564 | 0.009453583 | 0.009453583 | 0.057381221 | Alcam/Cdh13 | 2 |
| GO:0006750 | BP | GO:0006750 | glutathione biosynthetic process | 1/20 | 14/28564 | 0.009760271 | 0.009760271 | 0.057381221 | Nfe2l2 | 1 |
| GO:0007194 | BP | GO:0007194 | negative regulation of adenylate cyclase activity | 1/20 | 14/28564 | 0.009760271 | 0.009760271 | 0.057381221 | Galr1 | 1 |
| GO:0021670 | BP | GO:0021670 | lateral ventricle development | 1/20 | 14/28564 | 0.009760271 | 0.009760271 | 0.057381221 | Atp1b2 | 1 |
| GO:0036499 | BP | GO:0036499 | PERK-mediated unfolded protein response | 1/20 | 14/28564 | 0.009760271 | 0.009760271 | 0.057381221 | Nfe2l2 | 1 |
| GO:0060670 | BP | GO:0060670 | branching involved in labyrinthine layer morphogenesis | 1/20 | 14/28564 | 0.009760271 | 0.009760271 | 0.057381221 | Rspo3 | 1 |
| GO:1901533 | BP | GO:1901533 | negative regulation of hematopoietic progenitor cell differentiation | 1/20 | 14/28564 | 0.009760271 | 0.009760271 | 0.057381221 | Nfe2l2 | 1 |
| GO:1902036 | BP | GO:1902036 | regulation of hematopoietic stem cell differentiation | 1/20 | 14/28564 | 0.009760271 | 0.009760271 | 0.057381221 | Nfe2l2 | 1 |
| GO:1904753 | BP | GO:1904753 | negative regulation of vascular associated smooth muscle cell migration | 1/20 | 14/28564 | 0.009760271 | 0.009760271 | 0.057381221 | Nfe2l2 | 1 |
| GO:2000052 | BP | GO:2000052 | positive regulation of non-canonical Wnt signaling pathway | 1/20 | 14/28564 | 0.009760271 | 0.009760271 | 0.057381221 | Rspo3 | 1 |
| GO:0021932 | BP | GO:0021932 | hindbrain radial glia guided cell migration | 1/20 | 15/28564 | 0.010453959 | 0.010453959 | 0.057381221 | Atp1b2 | 1 |
| GO:0030007 | BP | GO:0030007 | intracellular potassium ion homeostasis | 1/20 | 15/28564 | 0.010453959 | 0.010453959 | 0.057381221 | Atp1b2 | 1 |
| GO:0035933 | BP | GO:0035933 | glucocorticoid secretion | 1/20 | 15/28564 | 0.010453959 | 0.010453959 | 0.057381221 | Galr1 | 1 |
| GO:0036376 | BP | GO:0036376 | sodium ion export across plasma membrane | 1/20 | 15/28564 | 0.010453959 | 0.010453959 | 0.057381221 | Atp1b2 | 1 |
| GO:1903238 | BP | GO:1903238 | positive regulation of leukocyte tethering or rolling | 1/20 | 15/28564 | 0.010453959 | 0.010453959 | 0.057381221 | Gcnt1 | 1 |
| GO:0015693 | BP | GO:0015693 | magnesium ion transport | 1/20 | 16/28564 | 0.011147186 | 0.011147186 | 0.057381221 | Slc41a3 | 1 |
| GO:0043649 | BP | GO:0043649 | dicarboxylic acid catabolic process | 1/20 | 16/28564 | 0.011147186 | 0.011147186 | 0.057381221 | Aldh1l1 | 1 |
| GO:1902176 | BP | GO:1902176 | negative regulation of oxidative stress-induced intrinsic apoptotic signaling pathway | 1/20 | 16/28564 | 0.011147186 | 0.011147186 | 0.057381221 | Nfe2l2 | 1 |
| GO:0043542 | BP | GO:0043542 | endothelial cell migration | 2/20 | 230/28564 | 0.011148364 | 0.011148364 | 0.057381221 | Cdh13/Nfe2l2 | 2 |
| GO:0010976 | BP | GO:0010976 | positive regulation of neuron projection development | 2/20 | 237/28564 | 0.01180426 | 0.01180426 | 0.057381221 | Nfe2l2/Atp1b2 | 2 |
| GO:0071498 | BP | GO:0071498 | cellular response to fluid shear stress | 1/20 | 17/28564 | 0.011839951 | 0.011839951 | 0.057381221 | Nfe2l2 | 1 |
| GO:1901522 | BP | GO:1901522 | positive regulation of transcription from RNA polymerase II promoter involved in cellular response to chemical stim | 1/20 | 17/28564 | 0.011839951 | 0.011839951 | 0.057381221 | Nfe2l2 | 1 |
| GO:0019430 | BP | GO:0019430 | removal of superoxide radicals | 1/20 | 18/28564 | 0.012532255 | 0.012532255 | 0.057381221 | Nfe2l2 | 1 |
| GO:0071280 | BP | GO:0071280 | cellular response to copper ion | 1/20 | 18/28564 | 0.012532255 | 0.012532255 | 0.057381221 | Nfe2l2 | 1 |
| GO:0006813 | BP | GO:0006813 | potassium ion transport | 2/20 | 245/28564 | 0.012574301 | 0.012574301 | 0.057381221 | Kcnj3/Atp1b2 | 2 |
| GO:0140467 | BP | GO:0140467 | integrated stress response signaling | 1/20 | 20/28564 | 0.013915481 | 0.013915481 | 0.057381221 | Nfe2l2 | 1 |
| GO:0098657 | BP | GO:0098657 | import into cell | 2/20 | 264/28564 | 0.01448919 | 0.01448919 | 0.057381221 | Kcnj3/Atp1b2 | 2 |
| GO:0006740 | BP | GO:0006740 | NADPH regeneration | 1/20 | 22/28564 | 0.015296866 | 0.015296866 | 0.057381221 | Aldh1l1 | 1 |
| GO:0034764 | BP | GO:0034764 | positive regulation of transmembrane transport | 2/20 | 278/28564 | 0.015976178 | 0.015976178 | 0.057381221 | Nfe2l2/Atp1b2 | 2 |
| GO:0000303 | BP | GO:0000303 | response to superoxide | 1/20 | 23/28564 | 0.015986868 | 0.015986868 | 0.057381221 | Nfe2l2 | 1 |
| GO:1902307 | BP | GO:1902307 | positive regulation of sodium ion transmembrane transport | 1/20 | 23/28564 | 0.015986868 | 0.015986868 | 0.057381221 | Atp1b2 | 1 |
| GO:1904385 | BP | GO:1904385 | cellular response to angiotensin | 1/20 | 23/28564 | 0.015986868 | 0.015986868 | 0.057381221 | Nfe2l2 | 1 |
| GO:0006760 | BP | GO:0006760 | folic acid-containing compound metabolic process | 1/20 | 24/28564 | 0.016676412 | 0.016676412 | 0.057381221 | Aldh1l1 | 1 |
| GO:0034405 | BP | GO:0034405 | response to fluid shear stress | 1/20 | 24/28564 | 0.016676412 | 0.016676412 | 0.057381221 | Nfe2l2 | 1 |
| GO:1901032 | BP | GO:1901032 | negative regulation of response to reactive oxygen species | 1/20 | 24/28564 | 0.016676412 | 0.016676412 | 0.057381221 | Nfe2l2 | 1 |
| GO:1903206 | BP | GO:1903206 | negative regulation of hydrogen peroxide-induced cell death | 1/20 | 24/28564 | 0.016676412 | 0.016676412 | 0.057381221 | Nfe2l2 | 1 |
| GO:0006730 | BP | GO:0006730 | one-carbon metabolic process | 1/20 | 26/28564 | 0.018054122 | 0.018054122 | 0.057381221 | Aldh1l1 | 1 |
| GO:0030194 | BP | GO:0030194 | positive regulation of blood coagulation | 1/20 | 26/28564 | 0.018054122 | 0.018054122 | 0.057381221 | Nfe2l2 | 1 |
| GO:0050927 | BP | GO:0050927 | positive regulation of positive chemotaxis | 1/20 | 26/28564 | 0.018054122 | 0.018054122 | 0.057381221 | Cdh13 | 1 |
| GO:0061744 | BP | GO:0061744 | motor behavior | 1/20 | 26/28564 | 0.018054122 | 0.018054122 | 0.057381221 | Atp1b2 | 1 |
| GO:1900048 | BP | GO:1900048 | positive regulation of hemostasis | 1/20 | 26/28564 | 0.018054122 | 0.018054122 | 0.057381221 | Nfe2l2 | 1 |
| GO:1903975 | BP | GO:1903975 | regulation of glial cell migration | 1/20 | 26/28564 | 0.018054122 | 0.018054122 | 0.057381221 | Atp1b2 | 1 |
| GO:1990776 | BP | GO:1990776 | response to angiotensin | 1/20 | 26/28564 | 0.018054122 | 0.018054122 | 0.057381221 | Nfe2l2 | 1 |
| GO:1901018 | BP | GO:1901018 | positive regulation of potassium ion transmembrane transporter activity | 1/20 | 27/28564 | 0.01874229 | 0.01874229 | 0.057381221 | Atp1b2 | 1 |
| GO:1904996 | BP | GO:1904996 | positive regulation of leukocyte adhesion to vascular endothelial cell | 1/20 | 27/28564 | 0.01874229 | 0.01874229 | 0.057381221 | Gcnt1 | 1 |
| GO:0045765 | BP | GO:0045765 | regulation of angiogenesis | 2/20 | 305/28564 | 0.019021011 | 0.019021011 | 0.057381221 | Nfe2l2/Minar1 | 2 |
| GO:0046688 | BP | GO:0046688 | response to copper ion | 1/20 | 29/28564 | 0.02011725 | 0.02011725 | 0.057381221 | Nfe2l2 | 1 |
| GO:0050820 | BP | GO:0050820 | positive regulation of coagulation | 1/20 | 29/28564 | 0.02011725 | 0.02011725 | 0.057381221 | Nfe2l2 | 1 |
| GO:0001919 | BP | GO:0001919 | regulation of receptor recycling | 1/20 | 30/28564 | 0.020804043 | 0.020804043 | 0.057381221 | Gria3 | 1 |
| GO:0098869 | BP | GO:0098869 | cellular oxidant detoxification | 1/20 | 30/28564 | 0.020804043 | 0.020804043 | 0.057381221 | Nfe2l2 | 1 |
| GO:0090132 | BP | GO:0090132 | epithelium migration | 2/20 | 329/28564 | 0.021917356 | 0.021917356 | 0.057381221 | Cdh13/Nfe2l2 | 2 |
| GO:0045454 | BP | GO:0045454 | cell redox homeostasis | 1/20 | 32/28564 | 0.022176259 | 0.022176259 | 0.057381221 | Nfe2l2 | 1 |
| GO:2000352 | BP | GO:2000352 | negative regulation of endothelial cell apoptotic process | 1/20 | 32/28564 | 0.022176259 | 0.022176259 | 0.057381221 | Nfe2l2 | 1 |
| GO:2000737 | BP | GO:2000737 | negative regulation of stem cell differentiation | 1/20 | 32/28564 | 0.022176259 | 0.022176259 | 0.057381221 | Nfe2l2 | 1 |
| GO:0008045 | BP | GO:0008045 | motor neuron axon guidance | 1/20 | 33/28564 | 0.022861681 | 0.022861681 | 0.057381221 | Alcam | 1 |
| GO:0071404 | BP | GO:0071404 | cellular response to low-density lipoprotein particle stimulus | 1/20 | 34/28564 | 0.023546647 | 0.023546647 | 0.057381221 | Cdh13 | 1 |
| GO:0090322 | BP | GO:0090322 | regulation of superoxide metabolic process | 1/20 | 34/28564 | 0.023546647 | 0.023546647 | 0.057381221 | Nfe2l2 | 1 |
| GO:0055075 | BP | GO:0055075 | potassium ion homeostasis | 1/20 | 35/28564 | 0.024231157 | 0.024231157 | 0.057381221 | Atp1b2 | 1 |
| GO:0050878 | BP | GO:0050878 | regulation of body fluid levels | 2/20 | 354/28564 | 0.025117535 | 0.025117535 | 0.057381221 | Sytl2/Nfe2l2 | 2 |
| GO:0010667 | BP | GO:0010667 | negative regulation of cardiac muscle cell apoptotic process | 1/20 | 37/28564 | 0.02559881 | 0.02559881 | 0.057381221 | Nfe2l2 | 1 |
| GO:0040013 | BP | GO:0040013 | negative regulation of locomotion | 2/20 | 360/28564 | 0.025912651 | 0.025912651 | 0.057381221 | Nfe2l2/Atp1b2 | 2 |
| GO:0050901 | BP | GO:0050901 | leukocyte tethering or rolling | 1/20 | 38/28564 | 0.026281953 | 0.026281953 | 0.057381221 | Gcnt1 | 1 |
| GO:0071402 | BP | GO:0071402 | cellular response to lipoprotein particle stimulus | 1/20 | 38/28564 | 0.026281953 | 0.026281953 | 0.057381221 | Cdh13 | 1 |
| GO:1904738 | BP | GO:1904738 | vascular associated smooth muscle cell migration | 1/20 | 38/28564 | 0.026281953 | 0.026281953 | 0.057381221 | Nfe2l2 | 1 |
| GO:0006767 | BP | GO:0006767 | water-soluble vitamin metabolic process | 1/20 | 39/28564 | 0.026964641 | 0.026964641 | 0.057381221 | Aldh1l1 | 1 |
| GO:0031589 | BP | GO:0031589 | cell-substrate adhesion | 2/20 | 369/28564 | 0.027124593 | 0.027124593 | 0.057381221 | Cdh13/Atp1b2 | 2 |
| GO:0090162 | BP | GO:0090162 | establishment of epithelial cell polarity | 1/20 | 40/28564 | 0.027646874 | 0.027646874 | 0.057381221 | Frmd4b | 1 |
| GO:0006418 | BP | GO:0006418 | tRNA aminoacylation for protein translation | 1/20 | 41/28564 | 0.028328653 | 0.028328653 | 0.057381221 | Lars2 | 1 |
| GO:0006984 | BP | GO:0006984 | ER-nucleus signaling pathway | 1/20 | 43/28564 | 0.029690849 | 0.029690849 | 0.057381221 | Nfe2l2 | 1 |
| GO:0010765 | BP | GO:0010765 | positive regulation of sodium ion transport | 1/20 | 43/28564 | 0.029690849 | 0.029690849 | 0.057381221 | Atp1b2 | 1 |
| GO:0045761 | BP | GO:0045761 | regulation of adenylate cyclase activity | 1/20 | 43/28564 | 0.029690849 | 0.029690849 | 0.057381221 | Galr1 | 1 |
| GO:0008631 | BP | GO:0008631 | intrinsic apoptotic signaling pathway in response to oxidative stress | 1/20 | 44/28564 | 0.030371266 | 0.030371266 | 0.057381221 | Nfe2l2 | 1 |
| GO:0043039 | BP | GO:0043039 | tRNA aminoacylation | 1/20 | 44/28564 | 0.030371266 | 0.030371266 | 0.057381221 | Lars2 | 1 |
| GO:0044331 | BP | GO:0044331 | cell-cell adhesion mediated by cadherin | 1/20 | 44/28564 | 0.030371266 | 0.030371266 | 0.057381221 | Cdh13 | 1 |
| GO:0050850 | BP | GO:0050850 | positive regulation of calcium-mediated signaling | 1/20 | 44/28564 | 0.030371266 | 0.030371266 | 0.057381221 | Cdh13 | 1 |
| GO:0006904 | BP | GO:0006904 | vesicle docking involved in exocytosis | 1/20 | 45/28564 | 0.03105123 | 0.03105123 | 0.057381221 | Sytl2 | 1 |
| GO:0032781 | BP | GO:0032781 | positive regulation of ATP-dependent activity | 1/20 | 45/28564 | 0.03105123 | 0.03105123 | 0.057381221 | Atp1b2 | 1 |
| GO:0045494 | BP | GO:0045494 | photoreceptor cell maintenance | 1/20 | 45/28564 | 0.03105123 | 0.03105123 | 0.057381221 | Atp1b2 | 1 |
| GO:0050918 | BP | GO:0050918 | positive chemotaxis | 1/20 | 45/28564 | 0.03105123 | 0.03105123 | 0.057381221 | Cdh13 | 1 |
| GO:0086009 | BP | GO:0086009 | membrane repolarization | 1/20 | 45/28564 | 0.03105123 | 0.03105123 | 0.057381221 | Atp1b2 | 1 |
| GO:0046326 | BP | GO:0046326 | positive regulation of glucose import | 1/20 | 46/28564 | 0.031730741 | 0.031730741 | 0.058205772 | Nfe2l2 | 1 |
| GO:0006739 | BP | GO:0006739 | NADP metabolic process | 1/20 | 49/28564 | 0.033766559 | 0.033766559 | 0.060603367 | Aldh1l1 | 1 |
| GO:0016601 | BP | GO:0016601 | Rac protein signal transduction | 1/20 | 51/28564 | 0.035121512 | 0.035121512 | 0.061703471 | Cdh13 | 1 |
| GO:0043268 | BP | GO:0043268 | positive regulation of potassium ion transport | 1/20 | 52/28564 | 0.035798311 | 0.035798311 | 0.061901139 | Atp1b2 | 1 |
| GO:0001881 | BP | GO:0001881 | receptor recycling | 1/20 | 53/28564 | 0.036474659 | 0.036474659 | 0.061901139 | Gria3 | 1 |
| GO:0061756 | BP | GO:0061756 | leukocyte adhesion to vascular endothelial cell | 1/20 | 54/28564 | 0.037150556 | 0.037150556 | 0.062622203 | Gcnt1 | 1 |
| GO:0072577 | BP | GO:0072577 | endothelial cell apoptotic process | 1/20 | 55/28564 | 0.037826003 | 0.037826003 | 0.062791893 | Nfe2l2 | 1 |
| GO:0019748 | BP | GO:0019748 | secondary metabolic process | 1/20 | 57/28564 | 0.039175546 | 0.039175546 | 0.062791893 | Nfe2l2 | 1 |
| GO:0044060 | BP | GO:0044060 | regulation of endocrine process | 1/20 | 57/28564 | 0.039175546 | 0.039175546 | 0.062791893 | Galr1 | 1 |
| GO:0051047 | BP | GO:0051047 | positive regulation of secretion | 2/20 | 453/28564 | 0.039497765 | 0.039497765 | 0.062791893 | Sytl2/Galr1 | 2 |
| GO:0032941 | BP | GO:0032941 | secretion by tissue | 1/20 | 58/28564 | 0.039849643 | 0.039849643 | 0.062791893 | Sytl2 | 1 |
| GO:0034332 | BP | GO:0034332 | adherens junction organization | 1/20 | 58/28564 | 0.039849643 | 0.039849643 | 0.062791893 | Cdh13 | 1 |
| GO:0043536 | BP | GO:0043536 | positive regulation of blood vessel endothelial cell migration | 1/20 | 58/28564 | 0.039849643 | 0.039849643 | 0.062791893 | Nfe2l2 | 1 |
| GO:0031346 | BP | GO:0031346 | positive regulation of cell projection organization | 2/20 | 459/28564 | 0.040451365 | 0.040451365 | 0.062791893 | Nfe2l2/Atp1b2 | 2 |
| GO:0140353 | BP | GO:0140353 | lipid export from cell | 1/20 | 59/28564 | 0.040523291 | 0.040523291 | 0.062791893 | Galr1 | 1 |
| GO:0001974 | BP | GO:0001974 | blood vessel remodeling | 1/20 | 61/28564 | 0.04186924 | 0.04186924 | 0.063172591 | Rspo3 | 1 |
| GO:0043616 | BP | GO:0043616 | keratinocyte proliferation | 1/20 | 62/28564 | 0.042541542 | 0.042541542 | 0.063172591 | Cdh13 | 1 |
| GO:0090303 | BP | GO:0090303 | positive regulation of wound healing | 1/20 | 63/28564 | 0.043213395 | 0.043213395 | 0.063790562 | Nfe2l2 | 1 |
| GO:0042391 | BP | GO:0042391 | regulation of membrane potential | 2/20 | 488/28564 | 0.045183768 | 0.045183768 | 0.063979548 | Atp1b2/Gria3 | 2 |
| GO:0010659 | BP | GO:0010659 | cardiac muscle cell apoptotic process | 1/20 | 66/28564 | 0.04522627 | 0.04522627 | 0.063979548 | Nfe2l2 | 1 |
| GO:0032007 | BP | GO:0032007 | negative regulation of TOR signaling | 1/20 | 66/28564 | 0.04522627 | 0.04522627 | 0.063979548 | Minar1 | 1 |
| GO:0034765 | BP | GO:0034765 | regulation of monoatomic ion transmembrane transport | 2/20 | 492/28564 | 0.045852228 | 0.045852228 | 0.063979548 | Kcnj3/Atp1b2 | 2 |
| GO:0001954 | BP | GO:0001954 | positive regulation of cell-matrix adhesion | 1/20 | 67/28564 | 0.045896334 | 0.045896334 | 0.063979548 | Cdh13 | 1 |
| GO:0140115 | BP | GO:0140115 | export across plasma membrane | 1/20 | 68/28564 | 0.04656595 | 0.04656595 | 0.063979548 | Atp1b2 | 1 |
| GO:0006801 | BP | GO:0006801 | superoxide metabolic process | 1/20 | 69/28564 | 0.047235121 | 0.047235121 | 0.063979548 | Nfe2l2 | 1 |
| GO:0006749 | BP | GO:0006749 | glutathione metabolic process | 1/20 | 70/28564 | 0.047903845 | 0.047903845 | 0.063979548 | Nfe2l2 | 1 |
| GO:0042149 | BP | GO:0042149 | cellular response to glucose starvation | 1/20 | 70/28564 | 0.047903845 | 0.047903845 | 0.063979548 | Nfe2l2 | 1 |
| GO:0098754 | BP | GO:0098754 | detoxification | 1/20 | 71/28564 | 0.048572124 | 0.048572124 | 0.063979548 | Nfe2l2 | 1 |
| GO:0042058 | BP | GO:0042058 | regulation of epidermal growth factor receptor signaling pathway | 1/20 | 72/28564 | 0.049239956 | 0.049239956 | 0.063979548 | Cdh13 | 1 |
| GO:0030032 | BP | GO:0030032 | lamellipodium assembly | 1/20 | 73/28564 | 0.049907344 | 0.049907344 | 0.06417819 | Cdh13 | 1 |
| GO:0098688 | CC | GO:0098688 | parallel fiber to Purkinje cell synapse | 2/20 | 37/28585 | 0.000305223 | 0.000305223 | 0.012208902 | Kcnj3/Gria3 | 2 |
| GO:1902495 | CC | GO:1902495 | transmembrane transporter complex | 3/20 | 392/28585 | 0.002453132 | 0.002453132 | 0.02882308 | Kcnj3/Atp1b2/( | 3 |
| GO:1990351 | CC | GO:1990351 | transporter complex | 3/20 | 415/28585 | 0.002882308 | 0.002882308 | 0.02882308 | Kcnj3/Atp1b2/( | 3 |
| GO:0005912 | CC | GO:0005912 | adherens junction | 2/20 | 153/28585 | 0.005076274 | 0.005076274 | 0.040610194 | Frmd4b/Cdh1 | 2 |
| GO:0019897 | CC | GO:0019897 | extrinsic component of plasma membrane | 2/20 | 180/28585 | 0.006953981 | 0.006953981 | 0.043002515 | Sytl2/Cdh13 | 2 |
| GO:0098802 | CC | GO:0098802 | plasma membrane signaling receptor complex | 2/20 | 201/28585 | 0.008600503 | 0.008600503 | 0.043002515 | Alcam/Gria3 | 2 |
| GO:0042734 | CC | GO:0042734 | presynaptic membrane | 2/20 | 238/28585 | 0.011882709 | 0.011882709 | 0.043209852 | Kcnj3/Gria3 | 2 |
| GO:0097449 | CC | GO:0097449 | astrocyte projection | 1/20 | 22/28585 | 0.015285706 | 0.015285706 | 0.047032942 | Atp1b2 | 1 |
| GO:0043083 | CC | GO:0043083 | synaptic cleft | 1/20 | 25/28585 | 0.017352841 | 0.017352841 | 0.049185315 | Gria3 | 1 |
| GO:0032281 | CC | GO:0032281 | AMPA glutamate receptor complex | 1/20 | 29/28585 | 0.020102608 | 0.020102608 | 0.049185315 | Gria3 | 1 |
| GO:0019898 | CC | GO:0019898 | extrinsic component of membrane | 2/20 | 321/28585 | 0.020903759 | 0.020903759 | 0.049185315 | Sytl2/Cdh13 | 2 |
| GO:0042470 | CC | GO:0042470 | melanosome | 1/20 | 39/28585 | 0.026945081 | 0.026945081 | 0.052645163 | Sytl2 | 1 |
| GO:0048770 | CC | GO:0048770 | pigment granule | 1/20 | 39/28585 | 0.026945081 | 0.026945081 | 0.052645163 | Sytl2 | 1 |
| GO:0044298 | CC | GO:0044298 | cell body membrane | 1/20 | 41/28585 | 0.028308118 | 0.028308118 | 0.052645163 | Atp1b2 | 1 |
| GO:0001772 | CC | GO:0001772 | immunological synapse | 1/20 | 49/28585 | 0.033742147 | 0.033742147 | 0.052645163 | Alcam | 1 |
| GO:0043235 | CC | GO:0043235 | receptor complex | 2/20 | 419/28585 | 0.034219356 | 0.034219356 | 0.052645163 | Alcam/Gria3 | 2 |
| GO:0001917 | CC | GO:0001917 | photoreceptor inner segment | 1/20 | 61/28585 | 0.041839091 | 0.041839091 | 0.061505788 | Atp1b2 | 1 |
| GO:0031985 | CC | GO:0031985 | Golgi cisterna | 1/20 | 68/28585 | 0.046532498 | 0.046532498 | 0.061505788 | Gcnt1 | 1 |
| GO:0030315 | CC | GO:0030315 | T-tubule | 1/20 | 70/28585 | 0.047869455 | 0.047869455 | 0.061505788 | Kcnj3 | 1 |
| GO:0043198 | CC | GO:0043198 | dendritic shaft | 1/20 | 71/28585 | 0.048537265 | 0.048537265 | 0.061505788 | Gria3 | 1 |
| GO:0042562 | MF | GO:0042562 | hormone binding | 2/20 | 87/28171 | 0.00172771 | 0.00172771 | 0.026943226 | Cdh13/Galr1 | 2 |
| GO:0046873 | MF | GO:0046873 | metal ion transmembrane transporter activity | 3/20 | 430/28171 | 0.00331896 | 0.00331896 | 0.026943226 | Kcnj3/Slc41a3 | 3 |
| GO:0015276 | MF | GO:0015276 | ligand-gated monoatomic ion channel activity | 2/20 | 137/28171 | 0.004211994 | 0.004211994 | 0.026943226 | Kcnj3/Gria3 | 2 |
| GO:0022834 | MF | GO:0022834 | ligand-gated channel activity | 2/20 | 140/28171 | 0.004393574 | 0.004393574 | 0.026943226 | Kcnj3/Gria3 | 2 |
| GO:0008201 | MF | GO:0008201 | heparin binding | 2/20 | 174/28171 | 0.006698931 | 0.006698931 | 0.026943226 | Ltbp2/Rspo3 | 2 |
| GO:0005391 | MF | GO:0005391 | P-type sodium:potassium-exchanging transporter activity | 1/20 | 10/28171 | 0.007077988 | 0.007077988 | 0.026943226 | Atp1b2 | 1 |
| GO:0016742 | MF | GO:0016742 | hydroxymethyl-, formyl- and related transferase activity | 1/20 | 10/28171 | 0.007077988 | 0.007077988 | 0.026943226 | Aldh1l1 | 1 |
| GO:0002161 | MF | GO:0002161 | aminoacyl-tRNA editing activity | 1/20 | 13/28171 | 0.009192087 | 0.009192087 | 0.026943226 | Lars2 | 1 |
| GO:0042043 | MF | GO:0042043 | neurexin family protein binding | 1/20 | 14/28171 | 0.009895836 | 0.009895836 | 0.026943226 | Sytl2 | 1 |
| GO:0022853 | MF | GO:0022853 | active monoatomic ion transmembrane transporter activity | 2/20 | 225/28171 | 0.010976264 | 0.010976264 | 0.026943226 | Slc41a3/Atp1b | 2 |
| GO:0016646 | MF | GO:0016646 | oxidoreductase activity, acting on the CH-NH group of donors, NAD or NADP as acceptor | 1/20 | 16/28171 | 0.011301909 | 0.011301909 | 0.026943226 | Aldh1l1 | 1 |
| GO:0004029 | MF | GO:0004029 | aldehyde dehydrogenase (NAD+) activity | 1/20 | 17/28171 | 0.012004234 | 0.012004234 | 0.026943226 | Aldh1l1 | 1 |
| GO:0099508 | MF | GO:0099508 | voltage-gated monoatomic ion channel activity involved in regulation of presynaptic membrane potential | 1/20 | 17/28171 | 0.012004234 | 0.012004234 | 0.026943226 | Kcnj3 | 1 |
| GO:0030169 | MF | GO:0030169 | low-density lipoprotein particle binding | 1/20 | 18/28171 | 0.012706085 | 0.012706085 | 0.026943226 | Cdh13 | 1 |
| GO:0048038 | MF | GO:0048038 | quinone binding | 1/20 | 18/28171 | 0.012706085 | 0.012706085 | 0.026943226 | Sqor | 1 |
| GO:0140103 | MF | GO:0140103 | catalytic activity, acting on a glycoprotein | 1/20 | 22/28171 | 0.015508756 | 0.015508756 | 0.0274254 | Gcnt1 | 1 |
| GO:0005242 | MF | GO:0005242 | inward rectifier potassium channel activity | 1/20 | 23/28171 | 0.016208241 | 0.016208241 | 0.0274254 | Kcnj3 | 1 |
| GO:1901681 | MF | GO:1901681 | sulfur compound binding | 2/20 | 293/28171 | 0.018104282 | 0.018104282 | 0.0274254 | Ltbp2/Rspo3 | 2 |
| GO:0001671 | MF | GO:0001671 | ATPase activator activity | 1/20 | 26/28171 | 0.018303867 | 0.018303867 | 0.0274254 | Atp1b2 | 1 |
| GO:0008066 | MF | GO:0008066 | glutamate receptor activity | 1/20 | 27/28171 | 0.019001466 | 0.019001466 | 0.0274254 | Gria3 | 1 |
| GO:0015662 | MF | GO:0015662 | P-type ion transporter activity | 1/20 | 30/28171 | 0.021091438 | 0.021091438 | 0.0274254 | Atp1b2 | 1 |
| GO:0140358 | MF | GO:0140358 | P-type transmembrane transporter activity | 1/20 | 33/28171 | 0.02317718 | 0.02317718 | 0.029276438 | Atp1b2 | 1 |
| GO:0140828 | MF | GO:0140828 | metal cation:monoatomic cation antiporter activity | 1/20 | 36/28171 | 0.0252587 | 0.0252587 | 0.030936153 | Slc41a3 | 1 |
| GO:0042277 | MF | GO:0042277 | peptide binding | 2/20 | 361/28171 | 0.026722226 | 0.026722226 | 0.030936153 | Galr1/Gria3 | 2 |
| GO:0004812 | MF | GO:0004812 | aminoacyl-tRNA ligase activity | 1/20 | 41/28171 | 0.028718539 | 0.028718539 | 0.030936153 | Lars2 | 1 |
| GO:0016875 | MF | GO:0016875 | ligase activity, forming carbon-oxygen bonds | 1/20 | 41/28171 | 0.028718539 | 0.028718539 | 0.030936153 | Lars2 | 1 |
| GO:0005109 | MF | GO:0005109 | frizzled binding | 1/20 | 42/28171 | 0.029409105 | 0.029409105 | 0.030936153 | Rspo3 | 1 |
| GO:0008188 | MF | GO:0008188 | neuropeptide receptor activity | 1/20 | 43/28171 | 0.030099205 | 0.030099205 | 0.030936153 | Galr1 | 1 |
| GO:0071949 | MF | GO:0071949 | FAD binding | 1/20 | 44/28171 | 0.030788838 | 0.030788838 | 0.030936153 | Sqor | 1 |
| GO:0008375 | MF | GO:0008375 | acetylglucosaminyltransferase activity | 1/20 | 47/28171 | 0.032854945 | 0.032854945 | 0.031592234 | Gcnt1 | 1 |
| GO:0099529 | MF | GO:0099529 | neurotransmitter receptor activity involved in regulation of postsynaptic membrane potential | 1/20 | 53/28171 | 0.036974614 | 0.036974614 | 0.031592234 | Gria3 | 1 |
| GO:0016667 | MF | GO:0016667 | oxidoreductase activity, acting on a sulfur group of donors | 1/20 | 55/28171 | 0.038344128 | 0.038344128 | 0.031592234 | Sqor | 1 |
| GO:0001786 | MF | GO:0001786 | phosphatidylserine binding | 1/20 | 68/28171 | 0.047200974 | 0.047200974 | 0.03401021 | Sytl2 | 1 |
| GO:0045296 | MF | GO:0045296 | cadherin binding | 1/20 | 70/28171 | 0.048556668 | 0.048556668 | 0.03401021 | Cdh13 | 1 |

Cre+_vs_Cre-_enrichGO

| | ONTOLOGY | ID | Description | GeneRatio | BgRatio | pvalue | p.adjust | qvalue | geneID | Count |
|---|---|---|---|---|---|---|---|---|---|---|
| GO:1900271 | BP | GO:1900271 | regulation of long-term synaptic potentiation | 5/75 | 57/28564 | 4.10238975631642E-07 | 4.10238975631642E-07 | 0.000476741 | Apoe/Arc/Drd1/Fam107a/Scgn | 5 |
| GO:0031346 | BP | GO:0031346 | positive regulation of cell projection organization | 8/75 | 458/28564 | 2.75186283508555E-05 | 2.75186283508555E-05 | 0.011363409 | Frmd7/Atp1b2/Apoe/Enc1/Epha3/Islr2/Nfe2l2/Fzd1 | 8 |
| GO:0010976 | BP | GO:0010976 | positive regulation of neuron projection development | 6/75 | 237/28564 | 3.82329077719565E-05 | 3.82329077719565E-05 | 0.011363409 | Atp1b2/Apoe/Enc1/Epha3/Nfe2l2/Fzd1 | 6 |
| GO:0042176 | BP | GO:0042176 | regulation of protein catabolic process | 7/75 | 364/28564 | 4.86030258629382E-05 | 4.86030258629382E-05 | 0.011363409 | Wfs1/Apoe/Cst3/Sh3rf2/Ubqln2/Asb11/Gfap | 7 |
| GO:0007409 | BP | GO:0007409 | axonogenesis | 8/75 | 498/28564 | 4.89388544532294E-05 | 4.89388544532294E-05 | 0.011363409 | Apoe/Efna3/Sema3c/Epha3/Islr2/Ndn/B4gat1/Ptpro | 8 |
| GO:0046323 | BP | GO:0046323 | glucose import | 4/75 | 87/28564 | 8.27313489565513E-05 | 8.27313489565513E-05 | 0.013418799 | Slc1a2/Slc2a1/Drd1/Nfe2l2 | 4 |
| GO:0034764 | BP | GO:0034764 | positive regulation of transmembrane transport | 6/75 | 278/28564 | 9.23757912216128E-05 | 9.23757912216128E-05 | 0.013418799 | Slc1a2/Atp1b2/Arc/Drd1/Trdn/Nfe2l2 | 6 |
| GO:0042220 | BP | GO:0042220 | response to cocaine | 3/75 | 44/28564 | 0.000213105 | 0.000213105 | 0.022158078 | Slc1a2/Drd1/Smpd1 | 3 |
| GO:0098739 | BP | GO:0098739 | import across plasma membrane | 5/75 | 208/28564 | 0.000222672 | 0.000222672 | 0.022158078 | Kcnj3/Slc1a2/Atp1b2/Slc2a1/Gfap | 5 |
| GO:1903202 | BP | GO:1903202 | negative regulation of oxidative stress-induced cell death | 3/75 | 51/28564 | 0.000330697 | 0.000330697 | 0.02401907 | Gpr37l1/Nfe2l2/Fzd1 | 3 |
| GO:0060020 | BP | GO:0060020 | Bergmann glial cell differentiation | 2/75 | 12/28564 | 0.000441381 | 0.000441381 | 0.026996406 | Gpr37l1/Gfap | 2 |
| GO:0030100 | BP | GO:0030100 | regulation of endocytosis | 5/75 | 248/28564 | 0.000498804 | 0.000498804 | 0.02761993 | Apoe/Arc/Epha3/Smpd1/Ubqln2 | 5 |
| GO:0006750 | BP | GO:0006750 | glutathione biosynthetic process | 2/75 | 14/28564 | 0.000606503 | 0.000606503 | 0.02819283 | Slc1a2/Nfe2l2 | 2 |
| GO:0042311 | BP | GO:0042311 | vasodilation | 3/75 | 64/28564 | 0.000645624 | 0.000645624 | 0.028469557 | Apoe/Drd1/Ednrb | 3 |
| GO:0098657 | BP | GO:0098657 | import into cell | 5/75 | 264/28564 | 0.000661453 | 0.000661453 | 0.028469557 | Kcnj3/Slc1a2/Atp1b2/Slc2a1/Gfap | 5 |
| GO:0008643 | BP | GO:0008643 | carbohydrate transport | 4/75 | 155/28564 | 0.000751516 | 0.000751516 | 0.028946149 | Slc1a2/Slc2a1/Drd1/Nfe2l2 | 4 |
| GO:1901215 | BP | GO:1901215 | negative regulation of neuron death | 5/75 | 273/28564 | 0.000768986 | 0.000768986 | 0.028946149 | Pcdhgc5/Wfs1/Apoe/Pcdhgc3/Fzd1 | 5 |
| GO:0043270 | BP | GO:0043270 | positive regulation of monoatomic ion transport | 5/75 | 274/28564 | 0.000781693 | 0.000781693 | 0.028946149 | Wfs1/Atp1b2/Arc/Drd1/Trdn | 5 |
| GO:0007190 | BP | GO:0007190 | activation of adenylate cyclase activity | 2/75 | 16/28564 | 0.000797068 | 0.000797068 | 0.028946149 | Drd1/Vipr2 | 2 |
| GO:0030032 | BP | GO:0030032 | lamellipodium assembly | 3/75 | 73/28564 | 0.000947617 | 0.000947617 | 0.030179298 | Frmd7/S1pr1/Ptpro | 3 |
| GO:0060560 | BP | GO:0060560 | developmental growth involved in morphogenesis | 5/75 | 290/28564 | 0.001007116 | 0.001007116 | 0.030179298 | Apoe/Sema3c/S1pr1/Islr2/Ndn | 5 |
| GO:0061318 | BP | GO:0061318 | renal filtration cell differentiation | 2/75 | 18/28564 | 0.001012811 | 0.001012811 | 0.030179298 | Ednrb/Ptpro | 2 |
| GO:0072112 | BP | GO:0072112 | podocyte differentiation | 2/75 | 18/28564 | 0.001012811 | 0.001012811 | 0.030179298 | Ednrb/Ptpro | 2 |
| GO:0072311 | BP | GO:0072311 | glomerular epithelial cell differentiation | 2/75 | 18/28564 | 0.001012811 | 0.001012811 | 0.030179298 | Ednrb/Ptpro | 2 |
| GO:1904294 | BP | GO:1904294 | positive regulation of ERAD pathway | 2/75 | 18/28564 | 0.001012811 | 0.001012811 | 0.030179298 | Wfs1/Ubqln2 | 2 |
| GO:0007568 | BP | GO:0007568 | aging | 3/75 | 78/28564 | 0.001148255 | 0.001148255 | 0.032236961 | Slc1a2/Aldoc/Apoe | 3 |
| GO:0015810 | BP | GO:0015810 | aspartate transmembrane transport | 2/75 | 20/28564 | 0.00125347 | 0.00125347 | 0.032236961 | Slc1a2/Gfap | 2 |
| GO:0050807 | BP | GO:0050807 | regulation of synapse organization | 5/75 | 305/28564 | 0.001259339 | 0.001259339 | 0.032236961 | Apoe/Arc/Pcdh8/Ptpro/Fzd1 | 5 |
| GO:0060070 | BP | GO:0060070 | canonical Wnt signaling pathway | 5/75 | 305/28564 | 0.001259339 | 0.001259339 | 0.032236961 | Apoe/Lgr5/Ednrb/Ptpro/Fzd1 | 5 |
| GO:0198738 | BP | GO:0198738 | cell-cell signaling by wnt | 6/75 | 460/28564 | 0.001331527 | 0.001331527 | 0.032236961 | Apoe/Lgr6/Lgr5/Ednrb/Ptpro/Fzd1 | 6 |
| GO:0150104 | BP | GO:0150104 | transport across blood-brain barrier | 2/75 | 21/28564 | 0.001383061 | 0.001383061 | 0.032801278 | Apoe/Slc2a1 | 2 |
| GO:0050803 | BP | GO:0050803 | regulation of synapse structure or activity | 5/75 | 313/28564 | 0.001411565 | 0.001411565 | 0.032807752 | Apoe/Arc/Pcdh8/Ptpro/Fzd1 | 5 |
| GO:0006874 | BP | GO:0006874 | intracellular calcium ion homeostasis | 5/75 | 316/28564 | 0.001472012 | 0.001472012 | 0.033541829 | Wfs1/Apoe/Drd1/Trdn/Scgn | 5 |
| GO:0097581 | BP | GO:0097581 | lamellipodium organization | 3/75 | 91/28564 | 0.001789436 | 0.001789436 | 0.03562141 | Frmd7/S1pr1/Ptpro | 3 |
| GO:1901032 | BP | GO:1901032 | negative regulation of response to reactive oxygen species | 2/75 | 24/28564 | 0.001808496 | 0.001808496 | 0.03562141 | Gpr37l1/Nfe2l2 | 2 |
| GO:0043524 | BP | GO:0043524 | negative regulation of neuron apoptotic process | 4/75 | 199/28564 | 0.001888559 | 0.001888559 | 0.035633441 | Pcdhgc5/Wfs1/Apoe/Pcdhgc3 | 4 |
| GO:0009410 | BP | GO:0009410 | response to xenobiotic stimulus | 5/75 | 335/28564 | 0.001900193 | 0.001900193 | 0.035633441 | Slc1a2/Drd1/Nfe2l2/Fzd1/Hadh | 5 |
| GO:0007413 | BP | GO:0007413 | axonal fasciculation | 2/75 | 25/28564 | 0.001962421 | 0.001962421 | 0.035633441 | Epha3/Ndn | 2 |
| GO:0106030 | BP | GO:0106030 | neuron projection fasciculation | 2/75 | 25/28564 | 0.001962421 | 0.001962421 | 0.035633441 | Epha3/Ndn | 2 |
| GO:1902074 | BP | GO:1902074 | response to salt | 5/75 | 340/28564 | 0.002026813 | 0.002026813 | 0.036062161 | Slc1a2/Drd1/Cpne6/Smpd1/Ednrb | 5 |
| GO:0045762 | BP | GO:0045762 | positive regulation of adenylate cyclase activity | 2/75 | 26/28564 | 0.00212235 | 0.00212235 | 0.036062161 | Drd1/Vipr2 | 2 |
| GO:1902993 | BP | GO:1902993 | positive regulation of amyloid precursor protein catabolic process | 2/75 | 26/28564 | 0.00212235 | 0.00212235 | 0.036062161 | Apoe/Efna3 | 2 |
| GO:0006469 | BP | GO:0006469 | negative regulation of protein kinase activity | 4/75 | 206/28564 | 0.002141191 | 0.002141191 | 0.036062161 | Apoe/Drd1/Smpd1/Ptpro | 4 |
| GO:0055074 | BP | GO:0055074 | calcium ion homeostasis | 5/75 | 347/28564 | 0.002213805 | 0.002213805 | 0.036734509 | Wfs1/Apoe/Drd1/Trdn/Scgn | 5 |
| GO:0036473 | BP | GO:0036473 | cell death in response to oxidative stress | 3/75 | 101/28564 | 0.002409113 | 0.002409113 | 0.037899121 | Gpr37l1/Nfe2l2/Fzd1 | 3 |
| GO:0050878 | BP | GO:0050878 | regulation of body fluid levels | 5/75 | 354/28564 | 0.002413323 | 0.002413323 | 0.037899121 | Wfs1/Apoe/Ednrb/Ptpro/Nfe2l2 | 5 |
| GO:0015740 | BP | GO:0015740 | C4-dicarboxylate transport | 2/75 | 28/28564 | 0.00246009 | 0.00246009 | 0.038118447 | Slc1a2/Gfap | 2 |
| GO:0040013 | BP | GO:0040013 | negative regulation of locomotion | 5/75 | 360/28564 | 0.002584503 | 0.002584503 | 0.039286066 | Atp1b2/Apoe/Drd1/Sema3c/Nfe2l2 | 5 |
| GO:0003071 | BP | GO:0003071 | renal system process involved in regulation of systemic arterial blood pressure | 2/75 | 29/28564 | 0.002637839 | 0.002637839 | 0.039286066 | Ednrb/Ptpro | 2 |
| GO:0014821 | BP | GO:0014821 | phasic smooth muscle contraction | 2/75 | 29/28564 | 0.002637839 | 0.002637839 | 0.039286066 | Drd1/Ednrb | 2 |
| GO:0031345 | BP | GO:0031345 | negative regulation of cell projection organization | 4/75 | 219/28564 | 0.00267067 | 0.00267067 | 0.039286066 | Apoe/Sema3c/Epha3/Gfap | 4 |
| GO:0050890 | BP | GO:0050890 | cognition | 5/75 | 364/28564 | 0.002720658 | 0.002720658 | 0.03952114 | Gpr88/Apoe/Arc/Drd1/Fam107a | 5 |
| GO:0015849 | BP | GO:0015849 | organic acid transport | 5/75 | 369/28564 | 0.002884545 | 0.002884545 | 0.040234643 | Slc32a1/Slc1a2/Apoe/Slc2a1/Gfap | 5 |
| GO:0007188 | BP | GO:0007188 | adenylate cyclase-modulating G protein-coupled receptor signaling pathway | 4/75 | 225/28564 | 0.002942887 | 0.002942887 | 0.040234643 | Drd1/Vipr2/S1pr1/Gpr37l1 | 4 |
| GO:0033673 | BP | GO:0033673 | negative regulation of kinase activity | 4/75 | 230/28564 | 0.003183783 | 0.003183783 | 0.042623704 | Apoe/Drd1/Smpd1/Ptpro | 4 |
| GO:0051497 | BP | GO:0051497 | negative regulation of stress fiber assembly | 2/75 | 34/28564 | 0.003614117 | 0.003614117 | 0.044476297 | Frmd7/S1pr1 | 2 |
| GO:0003018 | BP | GO:0003018 | vascular process in circulatory system | 4/75 | 240/28564 | 0.003705326 | 0.003705326 | 0.044476297 | Apoe/Slc2a1/Drd1/Ednrb | 4 |
| GO:0051651 | BP | GO:0051651 | maintenance of location in cell | 4/75 | 242/28564 | 0.003816178 | 0.003816178 | 0.044476297 | Apoe/Drd1/Trdn/Ftl1 | 4 |
| GO:0048013 | BP | GO:0048013 | ephrin receptor signaling pathway | 2/75 | 35/28564 | 0.003826662 | 0.003826662 | 0.044476297 | Efna3/Epha3 | 2 |
| GO:1905898 | BP | GO:1905898 | positive regulation of response to endoplasmic reticulum stress | 2/75 | 35/28564 | 0.003826662 | 0.003826662 | 0.044476297 | Wfs1/Ubqln2 | 2 |
| GO:0033555 | BP | GO:0033555 | multicellular organismal response to stress | 3/75 | 119/28564 | 0.003927209 | 0.003927209 | 0.044476297 | Apoe/Drd1/Ednrb | 3 |
| GO:0099560 | BP | GO:0099560 | synaptic membrane adhesion | 2/75 | 36/28564 | 0.004044899 | 0.004044899 | 0.046540583 | Pcdh8/Epha3 | 2 |
| GO:0006979 | BP | GO:0006979 | response to oxidative stress | 5/75 | 408/28564 | 0.004417859 | 0.004417859 | 0.049365546 | Apoe/Cst3/Gpr37l1/Nfe2l2/Fzd1 | 5 |
| GO:0019934 | BP | GO:0019934 | cGMP-mediated signaling | 2/75 | 39/28564 | 0.004733455 | 0.004733455 | 0.050653116 | Apoe/Ednrb | 2 |
| GO:0061387 | BP | GO:0061387 | regulation of extent of cell growth | 3/75 | 129/28564 | 0.004794611 | 0.004794611 | 0.050653116 | Apoe/Sema3c/Islr2 | 3 |
| GO:0032543 | BP | GO:0032543 | mitochondrial translation | 3/75 | 132/28564 | 0.00511099 | 0.00511099 | 0.051412245 | Lars2/Mrpl41/Mrps2 | 3 |
| GO:0031668 | BP | GO:0031668 | cellular response to extracellular stimulus | 4/75 | 264/28564 | 0.00518741 | 0.00518741 | 0.051412245 | Slc1a2/Slc2a1/Fam107a/Nfe2l2 | 4 |
| GO:0051348 | BP | GO:0051348 | negative regulation of transferase activity | 4/75 | 269/28564 | 0.005539548 | 0.005539548 | 0.052842304 | Apoe/Drd1/Smpd1/Ptpro | 4 |
| GO:0003014 | BP | GO:0003014 | renal system process | 3/75 | 136/28564 | 0.005552014 | 0.005552014 | 0.052842304 | Wfs1/Ednrb/Ptpro | 3 |
| GO:0045761 | BP | GO:0045761 | regulation of adenylate cyclase activity | 2/75 | 43/28564 | 0.005729369 | 0.005729369 | 0.052842304 | Drd1/Vipr2 | 2 |
| GO:0051281 | BP | GO:0051281 | positive regulation of release of sequestered calcium ion into cytosol | 2/75 | 43/28564 | 0.005729369 | 0.005729369 | 0.052842304 | Drd1/Trdn | 2 |
| GO:0098801 | BP | GO:0098801 | regulation of renal system process | 2/75 | 43/28564 | 0.005729369 | 0.005729369 | 0.052842304 | Ednrb/Ptpro | 2 |
| GO:0001504 | BP | GO:0001504 | neurotransmitter uptake | 2/75 | 44/28564 | 0.005992049 | 0.005992049 | 0.054188701 | Slc1a2/Gfap | 2 |
| GO:0032411 | BP | GO:0032411 | positive regulation of transporter activity | 3/75 | 140/28564 | 0.00601524 | 0.00601524 | 0.054188701 | Atp1b2/Arc/Trdn | 3 |
| GO:1905039 | BP | GO:1905039 | carboxylic acid transmembrane transport | 3/75 | 143/28564 | 0.006377405 | 0.006377405 | 0.056574165 | Slc1a2/Slc2a1/Gfap | 3 |
| GO:0009991 | BP | GO:0009991 | response to extracellular stimulus | 5/75 | 447/28564 | 0.006460601 | 0.006460601 | 0.05666252 | Slc1a2/Apoe/Slc2a1/Fam107a/Nfe2l2 | 5 |
| GO:0046326 | BP | GO:0046326 | positive regulation of glucose import | 2/75 | 46/28564 | 0.00653364 | 0.00653364 | 0.05666252 | Slc1a2/Nfe2l2 | 2 |
| GO:0048863 | BP | GO:0048863 | stem cell differentiation | 4/75 | 286/28564 | 0.006855745 | 0.006855745 | 0.059015538 | Sema3c/Ednrb/Nfe2l2/Fzd1 | 4 |
| GO:0048588 | BP | GO:0048588 | developmental cell growth | 4/75 | 287/28564 | 0.006939055 | 0.006939055 | 0.059293475 | Apoe/Sema3c/Islr2/Ndn | 4 |
| GO:1990573 | BP | GO:1990573 | potassium ion import across plasma membrane | 2/75 | 48/28564 | 0.00709668 | 0.00709668 | 0.059761513 | Kcnj3/Atp1b2 | 2 |
| GO:0070997 | BP | GO:0070997 | neuron death | 5/75 | 461/28564 | 0.007333417 | 0.007333417 | 0.060872873 | Pcdhgc5/Wfs1/Apoe/Pcdhgc3/Fzd1 | 5 |
| GO:0034248 | BP | GO:0034248 | regulation of amide metabolic process | 5/75 | 473/28564 | 0.008144619 | 0.008144619 | 0.066188145 | Wfs1/Apoe/Enc1/Rplp1/Nfe2l2 | 5 |
| GO:0050767 | BP | GO:0050767 | regulation of neurogenesis | 5/75 | 480/28564 | 0.008645636 | 0.008645636 | 0.068816021 | Apoe/Sema3c/Islr2/Gpr37l1/Gfap | 5 |
| GO:0007617 | BP | GO:0007617 | mating behavior | 2/75 | 54/28564 | 0.008912158 | 0.008912158 | 0.070448235 | Drd1/Ednrb | 2 |
| GO:0030336 | BP | GO:0030336 | negative regulation of cell migration | 4/75 | 312/28564 | 0.009244826 | 0.009244826 | 0.070680661 | Atp1b2/Apoe/Drd1/Nfe2l2 | 4 |
| GO:0001662 | BP | GO:0001662 | behavioral fear response | 2/75 | 56/28564 | 0.00955867 | 0.00955867 | 0.07166568 | Apoe/Drd1 | 2 |
| GO:0140053 | BP | GO:0140053 | mitochondrial gene expression | 3/75 | 167/28564 | 0.009743014 | 0.009743014 | 0.071917782 | Lars2/Mrpl41/Mrps2 | 3 |
| GO:0019228 | BP | GO:0019228 | neuronal action potential | 2/75 | 57/28564 | 0.009889551 | 0.009889551 | 0.071917782 | Gpr88/Drd1 | 2 |
| GO:0051017 | BP | GO:0051017 | actin filament bundle assembly | 3/75 | 173/28564 | 0.010717899 | 0.010717899 | 0.075484003 | Frmd7/Fam107a/S1pr1 | 3 |
| GO:0009416 | BP | GO:0009416 | response to light stimulus | 4/75 | 327/28564 | 0.010842758 | 0.010842758 | 0.075484003 | Slc1a2/Gpr88/Drd1/Smpd1 | 4 |
| GO:0071496 | BP | GO:0071496 | cellular response to external stimulus | 4/75 | 327/28564 | 0.010842758 | 0.010842758 | 0.075484003 | Slc1a2/Slc2a1/Fam107a/Nfe2l2 | 4 |
| GO:0042987 | BP | GO:0042987 | amyloid precursor protein catabolic process | 2/75 | 60/28564 | 0.010912361 | 0.010912361 | 0.075484003 | Apoe/Efna3 | 2 |
| GO:0055078 | BP | GO:0055078 | sodium ion homeostasis | 2/75 | 60/28564 | 0.010912361 | 0.010912361 | 0.075484003 | Atp1b2/Ednrb | 2 |
| GO:0071248 | BP | GO:0071248 | cellular response to metal ion | 3/75 | 176/28564 | 0.011225799 | 0.011225799 | 0.076544406 | Cpne6/Smpd1/Nfe2l2 | 3 |
| GO:0001974 | BP | GO:0001974 | blood vessel remodeling | 2/75 | 61/28564 | 0.01126326 | 0.01126326 | 0.076544406 | Cst3/Sema3c | 2 |
| GO:1902904 | BP | GO:1902904 | negative regulation of supramolecular fiber organization | 3/75 | 178/28564 | 0.011572022 | 0.011572022 | 0.077916261 | Frmd7/Apoe/S1pr1 | 3 |
| GO:0035296 | BP | GO:0035296 | regulation of tube diameter | 3/75 | 179/28564 | 0.011747427 | 0.011747427 | 0.077916261 | Apoe/Drd1/Ednrb | 3 |
| GO:0001755 | BP | GO:0001755 | neural crest cell migration | 2/75 | 63/28564 | 0.011979833 | 0.011979833 | 0.077916261 | Sema3c/Ednrb | 2 |
| GO:0051402 | BP | GO:0051402 | neuron apoptotic process | 4/75 | 337/28564 | 0.012001504 | 0.012001504 | 0.077916261 | Pcdhgc5/Wfs1/Apoe/Pcdhgc3 | 4 |
| GO:0051893 | BP | GO:0051893 | regulation of focal adhesion assembly | 2/75 | 65/28564 | 0.012715924 | 0.012715924 | 0.081193641 | Fam107a/Epha3 | 2 |
| GO:0090109 | BP | GO:0090109 | regulation of cell-substrate junction assembly | 2/75 | 65/28564 | 0.012715924 | 0.012715924 | 0.081193641 | Fam107a/Epha3 | 2 |
| GO:0090497 | BP | GO:0090497 | mesenchymal cell migration | 2/75 | 65/28564 | 0.012715924 | 0.012715924 | 0.081193641 | Sema3c/Ednrb | 2 |
| GO:0043409 | BP | GO:0043409 | negative regulation of MAPK cascade | 3/75 | 187/28564 | 0.013205993 | 0.013205993 | 0.083406274 | Apoe/Sh3rf2/Smpd1 | 3 |
| GO:0015909 | BP | GO:0015909 | long-chain fatty acid transport | 2/75 | 67/28564 | 0.013471315 | 0.013471315 | 0.084167131 | Apoe/Slc2a1 | 2 |
| GO:0001659 | BP | GO:0001659 | temperature homeostasis | 3/75 | 192/28564 | 0.014167641 | 0.014167641 | 0.086201691 | Drd1/Ednrb/Hadh | 3 |
| GO:0048660 | BP | GO:0048660 | regulation of smooth muscle cell proliferation | 3/75 | 192/28564 | 0.014167641 | 0.014167641 | 0.086201691 | Apoe/Vipr2/S1pr1 | 3 |
| GO:0006749 | BP | GO:0006749 | glutathione metabolic process | 2/75 | 70/28564 | 0.014640114 | 0.014640114 | 0.087247964 | Slc1a2/Nfe2l2 | 2 |
| GO:0150116 | BP | GO:0150116 | regulation of cell-substrate junction organization | 2/75 | 70/28564 | 0.014640114 | 0.014640114 | 0.087247964 | Fam107a/Epha3 | 2 |
| GO:0006879 | BP | GO:0006879 | intracellular iron ion homeostasis | 2/75 | 71/28564 | 0.015039129 | 0.015039129 | 0.087385253 | Hephl1/Ftl1 | 2 |
| GO:0030193 | BP | GO:0030193 | regulation of blood coagulation | 2/75 | 71/28564 | 0.015039129 | 0.015039129 | 0.087385253 | Apoe/Nfe2l2 | 2 |
| GO:1904036 | BP | GO:1904036 | negative regulation of epithelial cell apoptotic process | 2/75 | 71/28564 | 0.015039129 | 0.015039129 | 0.087385253 | Wfs1/Nfe2l2 | 2 |
| GO:1902075 | BP | GO:1902075 | cellular response to salt | 3/75 | 197/28564 | 0.015168596 | 0.015168596 | 0.087668098 | Slc1a2/Cpne6/Smpd1 | 3 |
| GO:0032956 | BP | GO:0032956 | regulation of actin cytoskeleton organization | 4/75 | 362/28564 | 0.015238337 | 0.015238337 | 0.087668098 | Frmd7/Fam107a/S1pr1/Epha3 | 4 |
| GO:0030968 | BP | GO:0030968 | endoplasmic reticulum unfolded protein response | 2/75 | 72/28564 | 0.015442807 | 0.015442807 | 0.087971409 | Wfs1/Nfe2l2 | 2 |
| GO:0048659 | BP | GO:0048659 | smooth muscle cell proliferation | 3/75 | 200/28564 | 0.015787801 | 0.015787801 | 0.088988757 | Apoe/Vipr2/S1pr1 | 3 |
| GO:1900046 | BP | GO:1900046 | regulation of hemostasis | 2/75 | 73/28564 | 0.015851122 | 0.015851122 | 0.088988757 | Apoe/Nfe2l2 | 2 |
| GO:0050818 | BP | GO:0050818 | regulation of coagulation | 2/75 | 75/28564 | 0.016681557 | 0.016681557 | 0.092754699 | Apoe/Nfe2l2 | 2 |
| GO:0001756 | BP | GO:0001756 | somitogenesis | 2/75 | 77/28564 | 0.017530224 | 0.017530224 | 0.096094175 | Pcdh8/Sema3c | 2 |
| GO:0098742 | BP | GO:0098742 | cell-cell adhesion via plasma-membrane adhesion molecules | 3/75 | 211/28564 | 0.018179132 | 0.018179132 | 0.098569968 | Pcdhgc3/Pcdh8/Epha3 | 3 |
| GO:0007632 | BP | GO:0007632 | visual behavior | 2/75 | 79/28564 | 0.018396912 | 0.018396912 | 0.098569968 | Slc1a2/Drd1 | 2 |
| GO:0031396 | BP | GO:0031396 | regulation of protein ubiquitination | 3/75 | 214/28564 | 0.018864406 | 0.018864406 | 0.100561583 | Wfs1/Fam107a/Sh3rf2 | 3 |
| GO:0008361 | BP | GO:0008361 | regulation of cell size | 3/75 | 218/28564 | 0.019800229 | 0.019800229 | 0.103183634 | Apoe/Sema3c/Islr2 | 3 |
| GO:0002090 | BP | GO:0002090 | regulation of receptor internalization | 2/75 | 83/28564 | 0.02018353 | 0.02018353 | 0.104711545 | Arc/Ubqln2 | 2 |
| GO:0060411 | BP | GO:0060411 | cardiac septum morphogenesis | 2/75 | 85/28564 | 0.02110305 | 0.02110305 | 0.108513119 | Sema3c/Fzd1 | 2 |
| GO:0006836 | BP | GO:0006836 | neurotransmitter transport | 3/75 | 225/28564 | 0.021498855 | 0.021498855 | 0.110061378 | Slc32a1/Slc1a2/Gfap | 3 |
| GO:0034103 | BP | GO:0034103 | regulation of tissue remodeling | 2/75 | 87/28564 | 0.022039773 | 0.022039773 | 0.111358854 | Cst3/S1pr1 | 2 |
| GO:0060415 | BP | GO:0060415 | muscle tissue morphogenesis | 2/75 | 87/28564 | 0.022039773 | 0.022039773 | 0.111358854 | S1pr1/Fzd1 | 2 |
| GO:0048041 | BP | GO:0048041 | focal adhesion assembly | 2/75 | 88/28564 | 0.022514524 | 0.022514524 | 0.112667546 | Fam107a/Epha3 | 2 |
| GO:0030595 | BP | GO:0030595 | leukocyte chemotaxis | 3/75 | 230/28564 | 0.022759694 | 0.022759694 | 0.112667546 | S1pr1/Ednrb/Ptpro | 3 |
| GO:0032970 | BP | GO:0032970 | regulation of actin filament-based process | 4/75 | 411/28564 | 0.0230706 | 0.0230706 | 0.112667546 | Frmd7/Fam107a/S1pr1/Epha3 | 4 |
| GO:0035023 | BP | GO:0035023 | regulation of Rho protein signal transduction | 2/75 | 90/28564 | 0.023476678 | 0.023476678 | 0.112667546 | Apoe/Stard8 | 2 |
| GO:0009952 | BP | GO:0009952 | anterior/posterior pattern specification | 3/75 | 233/28564 | 0.023535219 | 0.023535219 | 0.112667546 | Arc/Pcdh8/Sema3c | 3 |
| GO:0050709 | BP | GO:0050709 | negative regulation of protein secretion | 2/75 | 92/28564 | 0.024455537 | 0.024455537 | 0.112667546 | Apoe/Hadh | 2 |
| GO:0007191 | BP | GO:0007191 | adenylate cyclase-activating dopamine receptor signaling pathway | 1/75 | 10/28564 | 0.02595279 | 0.02595279 | 0.112667546 | Drd1 | 1 |
| GO:0010496 | BP | GO:0010496 | intercellular transport | 1/75 | 10/28564 | 0.02595279 | 0.02595279 | 0.112667546 | Arc | 1 |
| GO:0021826 | BP | GO:0021826 | substrate-independent telencephalic tangential migration | 1/75 | 10/28564 | 0.02595279 | 0.02595279 | 0.112667546 | Drd1 | 1 |
| GO:0021843 | BP | GO:0021843 | substrate-independent telencephalic tangential interneuron migration | 1/75 | 10/28564 | 0.02595279 | 0.02595279 | 0.112667546 | Drd1 | 1 |
| GO:0034372 | BP | GO:0034372 | very-low-density lipoprotein particle remodeling | 1/75 | 10/28564 | 0.02595279 | 0.02595279 | 0.112667546 | Apoe | 1 |
| GO:0060022 | BP | GO:0060022 | hard palate development | 1/75 | 10/28564 | 0.02595279 | 0.02595279 | 0.112667546 | Fzd1 | 1 |
| GO:0060312 | BP | GO:0060312 | regulation of blood vessel remodeling | 1/75 | 10/28564 | 0.02595279 | 0.02595279 | 0.112667546 | Cst3 | 1 |
| GO:0061418 | BP | GO:0061418 | regulation of transcription from RNA polymerase II promoter in response to hypo | 1/75 | 10/28564 | 0.02595279 | 0.02595279 | 0.112667546 | Nfe2l2 | 1 |
| GO:0070294 | BP | GO:0070294 | renal sodium ion absorption | 1/75 | 10/28564 | 0.02595279 | 0.02595279 | 0.112667546 | Ednrb | 1 |
| GO:0098700 | BP | GO:0098700 | neurotransmitter loading into synaptic vesicle | 1/75 | 10/28564 | 0.02595279 | 0.02595279 | 0.112667546 | Slc32a1 | 1 |
| GO:1903054 | BP | GO:1903054 | negative regulation of extracellular matrix organization | 1/75 | 10/28564 | 0.02595279 | 0.02595279 | 0.112667546 | Cst3 | 1 |
| GO:2000646 | BP | GO:2000646 | positive regulation of receptor catabolic process | 1/75 | 10/28564 | 0.02595279 | 0.02595279 | 0.112667546 | Apoe | 1 |
| GO:0055072 | BP | GO:0055072 | iron ion homeostasis | 2/75 | 95/28564 | 0.025954718 | 0.025954718 | 0.112667546 | Hephl1/Ftl1 | 2 |
| GO:0006470 | BP | GO:0006470 | protein dephosphorylation | 3/75 | 243/28564 | 0.026223313 | 0.026223313 | 0.112667546 | Smpd1/Ptpn14/Ptpro | 3 |
| GO:0043535 | BP | GO:0043535 | regulation of blood vessel endothelial cell migration | 2/75 | 96/28564 | 0.026462586 | 0.026462586 | 0.112667546 | Apoe/Nfe2l2 | 2 |
| GO:0061053 | BP | GO:0061053 | somite development | 2/75 | 96/28564 | 0.026462586 | 0.026462586 | 0.112667546 | Pcdh8/Sema3c | 2 |
| GO:0045664 | BP | GO:0045664 | regulation of neuron differentiation | 3/75 | 244/28564 | 0.026500832 | 0.026500832 | 0.112667546 | Efna3/Gpr37l1/Ednrb | 3 |
| GO:0021892 | BP | GO:0021892 | cerebral cortex GABAergic interneuron differentiation | 1/75 | 11/28564 | 0.028511225 | 0.028511225 | 0.112667546 | Drd1 | 1 |
| GO:0021940 | BP | GO:0021940 | positive regulation of cerebellar granule cell precursor proliferation | 1/75 | 11/28564 | 0.028511225 | 0.028511225 | 0.112667546 | Gpr37l1 | 1 |
| GO:0031620 | BP | GO:0031620 | regulation of fever generation | 1/75 | 11/28564 | 0.028511225 | 0.028511225 | 0.112667546 | Ednrb | 1 |
| GO:0034384 | BP | GO:0034384 | high-density lipoprotein particle clearance | 1/75 | 11/28564 | 0.028511225 | 0.028511225 | 0.112667546 | Apoe | 1 |
| GO:0042045 | BP | GO:0042045 | epithelial fluid transport | 1/75 | 11/28564 | 0.028511225 | 0.028511225 | 0.112667546 | Ednrb | 1 |
| GO:0060174 | BP | GO:0060174 | limb bud formation | 1/75 | 11/28564 | 0.028511225 | 0.028511225 | 0.112667546 | Sema3c | 1 |
| GO:0060405 | BP | GO:0060405 | regulation of penile erection | 1/75 | 11/28564 | 0.028511225 | 0.028511225 | 0.112667546 | Ednrb | 1 |
| GO:0060600 | BP | GO:0060600 | dichotomous subdivision of an epithelial terminal unit | 1/75 | 11/28564 | 0.028511225 | 0.028511225 | 0.112667546 | Sema3c | 1 |
| GO:0071372 | BP | GO:0071372 | cellular response to follicle-stimulating hormone stimulus | 1/75 | 11/28564 | 0.028511225 | 0.028511225 | 0.112667546 | Epha3 | 1 |
| GO:0097278 | BP | GO:0097278 | complement-dependent cytotoxicity | 1/75 | 11/28564 | 0.028511225 | 0.028511225 | 0.112667546 | Cd59b | 1 |
| GO:2001243 | BP | GO:2001243 | negative regulation of intrinsic apoptotic signaling pathway | 2/75 | 100/28564 | 0.028534116 | 0.028534116 | 0.112667546 | Wfs1/Nfe2l2 | 2 |
| GO:0042158 | BP | GO:0042158 | lipoprotein biosynthetic process | 2/75 | 104/28564 | 0.030668597 | 0.030668597 | 0.112667546 | Apoe/Zdhhc22 | 2 |
| GO:0009404 | BP | GO:0009404 | toxin metabolic process | 1/75 | 12/28564 | 0.031063029 | 0.031063029 | 0.112667546 | Nfe2l2 | 1 |
| GO:0017038 | BP | GO:0017038 | protein import | 1/75 | 12/28564 | 0.031063029 | 0.031063029 | 0.112667546 | Apoe | 1 |
| GO:0030202 | BP | GO:0030202 | heparin metabolic process | 1/75 | 12/28564 | 0.031063029 | 0.031063029 | 0.112667546 | Ednrb | 1 |
| GO:0030388 | BP | GO:0030388 | fructose 1,6-bisphosphate metabolic process | 1/75 | 12/28564 | 0.031063029 | 0.031063029 | 0.112667546 | Aldoc | 1 |
| GO:0032354 | BP | GO:0032354 | response to follicle-stimulating hormone | 1/75 | 12/28564 | 0.031063029 | 0.031063029 | 0.112667546 | Epha3 | 1 |
| GO:0035810 | BP | GO:0035810 | positive regulation of urine volume | 1/75 | 12/28564 | 0.031063029 | 0.031063029 | 0.112667546 | Ednrb | 1 |
| GO:0061684 | BP | GO:0061684 | chaperone-mediated autophagy | 1/75 | 12/28564 | 0.031063029 | 0.031063029 | 0.112667546 | Gfap | 1 |
| GO:0090209 | BP | GO:0090209 | negative regulation of triglyceride metabolic process | 1/75 | 12/28564 | 0.031063029 | 0.031063029 | 0.112667546 | Apoe | 1 |
| GO:0097090 | BP | GO:0097090 | presynaptic membrane organization | 1/75 | 12/28564 | 0.031063029 | 0.031063029 | 0.112667546 | Apoe | 1 |
| GO:1900102 | BP | GO:1900102 | negative regulation of endoplasmic reticulum unfolded protein response | 1/75 | 12/28564 | 0.031063029 | 0.031063029 | 0.112667546 | Wfs1 | 1 |
| GO:1902947 | BP | GO:1902947 | regulation of tau-protein kinase activity | 1/75 | 12/28564 | 0.031063029 | 0.031063029 | 0.112667546 | Apoe | 1 |
| GO:1905907 | BP | GO:1905907 | negative regulation of amyloid fibril formation | 1/75 | 12/28564 | 0.031063029 | 0.031063029 | 0.112667546 | Apoe | 1 |
| GO:2000253 | BP | GO:2000253 | positive regulation of feeding behavior | 1/75 | 12/28564 | 0.031063029 | 0.031063029 | 0.112667546 | Drd1 | 1 |
| GO:0034976 | BP | GO:0034976 | response to endoplasmic reticulum stress | 3/75 | 260/28564 | 0.031155917 | 0.031155917 | 0.112667546 | Wfs1/Ubqln2/Nfe2l2 | 3 |
| GO:0006816 | BP | GO:0006816 | calcium ion transport | 4/75 | 452/28564 | 0.031209227 | 0.031209227 | 0.112667546 | Wfs1/Drd1/Ednrb/Trdn | 4 |
| GO:0150115 | BP | GO:0150115 | cell-substrate junction organization | 2/75 | 106/28564 | 0.031758972 | 0.031758972 | 0.112667546 | Fam107a/Epha3 | 2 |
| GO:0050795 | BP | GO:0050795 | regulation of behavior | 2/75 | 107/28564 | 0.032309862 | 0.032309862 | 0.112667546 | Apoe/Drd1 | 2 |
| GO:0097061 | BP | GO:0097061 | dendritic spine organization | 2/75 | 108/28564 | 0.032864522 | 0.032864522 | 0.112667546 | Apoe/Arc | 2 |
| GO:0048484 | BP | GO:0048484 | enteric nervous system development | 1/75 | 13/28564 | 0.033608219 | 0.033608219 | 0.112667546 | Ednrb | 1 |
| GO:0090158 | BP | GO:0090158 | endoplasmic reticulum membrane organization | 1/75 | 13/28564 | 0.033608219 | 0.033608219 | 0.112667546 | Trdn | 1 |
| GO:1903430 | BP | GO:1903430 | negative regulation of cell maturation | 1/75 | 13/28564 | 0.033608219 | 0.033608219 | 0.112667546 | Ednrb | 1 |
| GO:0007626 | BP | GO:0007626 | locomotory behavior | 3/75 | 270/28564 | 0.034269739 | 0.034269739 | 0.112667546 | Gpr88/Apoe/Drd1 | 3 |
| GO:0006898 | BP | GO:0006898 | receptor-mediated endocytosis | 3/75 | 273/28564 | 0.035234357 | 0.035234357 | 0.112667546 | Apoe/Arc/Ubqln2 | 3 |
| GO:0001660 | BP | GO:0001660 | fever generation | 1/75 | 14/28564 | 0.036146813 | 0.036146813 | 0.112667546 | Ednrb | 1 |
| GO:0032341 | BP | GO:0032341 | aldosterone metabolic process | 1/75 | 14/28564 | 0.036146813 | 0.036146813 | 0.112667546 | Ednrb | 1 |
| GO:0046598 | BP | GO:0046598 | positive regulation of viral entry into host cell | 1/75 | 14/28564 | 0.036146813 | 0.036146813 | 0.112667546 | Smpd1 | 1 |
| GO:0075294 | BP | GO:0075294 | positive regulation by symbiont of entry into host | 1/75 | 14/28564 | 0.036146813 | 0.036146813 | 0.112667546 | Smpd1 | 1 |
| GO:0090179 | BP | GO:0090179 | planar cell polarity pathway involved in neural tube closure | 1/75 | 14/28564 | 0.036146813 | 0.036146813 | 0.112667546 | Fzd1 | 1 |
| GO:0097113 | BP | GO:0097113 | AMPA glutamate receptor clustering | 1/75 | 14/28564 | 0.036146813 | 0.036146813 | 0.112667546 | Apoe | 1 |
| GO:0097688 | BP | GO:0097688 | glutamate receptor clustering | 1/75 | 14/28564 | 0.036146813 | 0.036146813 | 0.112667546 | Apoe | 1 |
| GO:1901533 | BP | GO:1901533 | negative regulation of hematopoietic progenitor cell differentiation | 1/75 | 14/28564 | 0.036146813 | 0.036146813 | 0.112667546 | Nfe2l2 | 1 |
| GO:1902036 | BP | GO:1902036 | regulation of hematopoietic stem cell differentiation | 1/75 | 14/28564 | 0.036146813 | 0.036146813 | 0.112667546 | Nfe2l2 | 1 |
| GO:1902931 | BP | GO:1902931 | negative regulation of alcohol biosynthetic process | 1/75 | 14/28564 | 0.036146813 | 0.036146813 | 0.112667546 | Apoe | 1 |
| GO:2000822 | BP | GO:2000822 | regulation of behavioral fear response | 1/75 | 14/28564 | 0.036146813 | 0.036146813 | 0.112667546 | Apoe | 1 |
| GO:2001140 | BP | GO:2001140 | positive regulation of phospholipid transport | 1/75 | 14/28564 | 0.036146813 | 0.036146813 | 0.112667546 | Apoe | 1 |
| GO:0043473 | BP | GO:0043473 | pigmentation | 2/75 | 114/28564 | 0.036270384 | 0.036270384 | 0.112667546 | Apoe/Ednrb | 2 |
| GO:0045446 | BP | GO:0045446 | endothelial cell differentiation | 2/75 | 116/28564 | 0.037434802 | 0.037434802 | 0.112667546 | Ednrb/Fzd1 | 2 |
| GO:0071482 | BP | GO:0071482 | cellular response to light stimulus | 2/75 | 117/28564 | 0.038022373 | 0.038022373 | 0.112667546 | Gpr88/Smpd1 | 2 |
| GO:0106027 | BP | GO:0106027 | neuron projection organization | 2/75 | 117/28564 | 0.038022373 | 0.038022373 | 0.112667546 | Apoe/Arc | 2 |
| GO:0006825 | BP | GO:0006825 | copper ion transport | 1/75 | 15/28564 | 0.038678826 | 0.038678826 | 0.112667546 | Hephl1 | 1 |
| GO:0015816 | BP | GO:0015816 | glycine transport | 1/75 | 15/28564 | 0.038678826 | 0.038678826 | 0.112667546 | Slc32a1 | 1 |
| GO:0021932 | BP | GO:0021932 | hindbrain radial glia guided cell migration | 1/75 | 15/28564 | 0.038678826 | 0.038678826 | 0.112667546 | Atp1b2 | 1 |
| GO:0030007 | BP | GO:0030007 | intracellular potassium ion homeostasis | 1/75 | 15/28564 | 0.038678826 | 0.038678826 | 0.112667546 | Atp1b2 | 1 |
| GO:0032372 | BP | GO:0032372 | negative regulation of sterol transport | 1/75 | 15/28564 | 0.038678826 | 0.038678826 | 0.112667546 | Apoe | 1 |
| GO:0032375 | BP | GO:0032375 | negative regulation of cholesterol transport | 1/75 | 15/28564 | 0.038678826 | 0.038678826 | 0.112667546 | Apoe | 1 |
| GO:0035269 | BP | GO:0035269 | protein O-linked mannosylation | 1/75 | 15/28564 | 0.038678826 | 0.038678826 | 0.112667546 | B4gat1 | 1 |
| GO:0035815 | BP | GO:0035815 | positive regulation of renal sodium excretion | 1/75 | 15/28564 | 0.038678826 | 0.038678826 | 0.112667546 | Ednrb | 1 |
| GO:0036376 | BP | GO:0036376 | sodium ion export across plasma membrane | 1/75 | 15/28564 | 0.038678826 | 0.038678826 | 0.112667546 | Atp1b2 | 1 |
| GO:0045916 | BP | GO:0045916 | negative regulation of complement activation | 1/75 | 15/28564 | 0.038678826 | 0.038678826 | 0.112667546 | Cd59b | 1 |
| GO:0070207 | BP | GO:0070207 | protein homotrimerization | 1/75 | 15/28564 | 0.038678826 | 0.038678826 | 0.112667546 | Slc1a2 | 1 |
| GO:0006986 | BP | GO:0006986 | response to unfolded protein | 2/75 | 119/28564 | [illegible] | [illegible] | 0.112667546 | Wfs1/Nfe2l2 | 2 |
| GO:0008212 | BP | GO:0008212 | mineralocorticoid metabolic process | 1/75 | 16/28564 | 0.041204277 | 0.041204277 | 0.112667546 | Ednrb | 1 |
| GO:0015693 | BP | GO:0015693 | magnesium ion transport | 1/75 | 16/28564 | 0.041204277 | 0.041204277 | 0.112667546 | Slc41a3 | 1 |
| GO:0031650 | BP | GO:0031650 | regulation of heat generation | 1/75 | 16/28564 | 0.041204277 | 0.041204277 | 0.112667546 | Ednrb | 1 |
| GO:0032488 | BP | GO:0032488 | Cdc42 protein signal transduction | 1/75 | 16/28564 | 0.041204277 | 0.041204277 | 0.112667546 | Apoe | 1 |
| GO:0033700 | BP | GO:0033700 | phospholipid efflux | 1/75 | 16/28564 | 0.041204277 | 0.041204277 | 0.112667546 | Apoe | 1 |
| GO:0042159 | BP | GO:0042159 | lipoprotein catabolic process | 1/75 | 16/28564 | 0.041204277 | 0.041204277 | 0.112667546 | Apoe | 1 |
| GO:0043649 | BP | GO:0043649 | dicarboxylic acid catabolic process | 1/75 | 16/28564 | 0.041204277 | 0.041204277 | 0.112667546 | Aldh1l1 | 1 |
| GO:0080009 | BP | GO:0080009 | mRNA methylation | 1/75 | 16/28564 | 0.041204277 | 0.041204277 | 0.112667546 | Trmt61a | 1 |
| GO:0097050 | BP | GO:0097050 | type B pancreatic cell apoptotic process | 1/75 | 16/28564 | 0.041204277 | 0.041204277 | 0.112667546 | Wfs1 | 1 |
| GO:1902950 | BP | GO:1902950 | regulation of dendritic spine maintenance | 1/75 | 16/28564 | 0.041204277 | 0.041204277 | 0.112667546 | Apoe | 1 |
| GO:1903019 | BP | GO:1903019 | negative regulation of glycoprotein metabolic process | 1/75 | 16/28564 | 0.041204277 | 0.041204277 | 0.112667546 | Cst3 | 1 |
| GO:0043534 | BP | GO:0043534 | blood vessel endothelial cell migration | 2/75 | 124/28564 | 0.042233378 | 0.042233378 | 0.114672036 | Apoe/Nfe2l2 | 2 |
| GO:0010715 | BP | GO:0010715 | regulation of extracellular matrix disassembly | 1/75 | 17/28564 | 0.043723181 | 0.043723181 | 0.114697379 | Cst3 | 1 |
| GO:0055089 | BP | GO:0055089 | fatty acid homeostasis | 1/75 | 17/28564 | 0.043723181 | 0.043723181 | 0.114697379 | Apoe | 1 |
| GO:0071498 | BP | GO:0071498 | cellular response to fluid shear stress | 1/75 | 17/28564 | 0.043723181 | 0.043723181 | 0.114697379 | Nfe2l2 | 1 |
| GO:0098712 | BP | GO:0098712 | L-glutamate import across plasma membrane | 1/75 | 17/28564 | 0.043723181 | 0.043723181 | 0.114697379 | Slc1a2 | 1 |
| GO:1901522 | BP | GO:1901522 | positive regulation of transcription from RNA polymerase II promoter involved in c | 1/75 | 17/28564 | 0.043723181 | 0.043723181 | 0.114697379 | Nfe2l2 | 1 |
| GO:1904936 | BP | GO:1904936 | interneuron migration | 1/75 | 17/28564 | 0.043723181 | 0.043723181 | 0.114697379 | Drd1 | 1 |
| GO:0001952 | BP | GO:0001952 | regulation of cell-matrix adhesion | 2/75 | 130/28564 | 0.045975373 | 0.045975373 | 0.116347068 | Fam107a/Epha3 | 2 |
| GO:0019430 | BP | GO:0019430 | removal of superoxide radicals | 1/75 | 18/28564 | 0.046235556 | 0.046235556 | 0.116347068 | Nfe2l2 | 1 |
| GO:0042249 | BP | GO:0042249 | establishment of planar polarity of embryonic epithelium | 1/75 | 18/28564 | 0.046235556 | 0.046235556 | 0.116347068 | Fzd1 | 1 |
| GO:1900221 | BP | GO:1900221 | regulation of amyloid-beta clearance | 1/75 | 18/28564 | 0.046235556 | 0.046235556 | 0.116347068 | Apoe | 1 |
| GO:0010038 | BP | GO:0010038 | response to metal ion | 3/75 | 309/28564 | 0.047890335 | 0.047890335 | 0.116347068 | Cpne6/Smpd1/Nfe2l2 | 3 |
| GO:0031647 | BP | GO:0031647 | regulation of protein stability | 3/75 | 311/28564 | 0.048651034 | 0.048651034 | 0.116347068 | Wfs1/Atp1b2/Fam107a | 3 |
| GO:0001991 | BP | GO:0001991 | regulation of systemic arterial blood pressure by circulatory renin-angiotensin | 1/75 | 19/28564 | 0.048741418 | 0.048741418 | 0.116347068 | Ednrb | 1 |
| GO:0002031 | BP | GO:0002031 | G protein-coupled receptor internalization | 1/75 | 19/28564 | 0.048741418 | 0.048741418 | 0.116347068 | Ubqln2 | 1 |
| GO:0010544 | BP | GO:0010544 | negative regulation of platelet activation | 1/75 | 19/28564 | 0.048741418 | 0.048741418 | 0.116347068 | Apoe | 1 |
| GO:0010713 | BP | GO:0010713 | negative regulation of collagen metabolic process | 1/75 | 19/28564 | 0.048741418 | 0.048741418 | 0.116347068 | Cst3 | 1 |
| GO:0042754 | BP | GO:0042754 | negative regulation of circadian rhythm | 1/75 | 19/28564 | 0.048741418 | 0.048741418 | 0.116347068 | Drd1 | 1 |
| GO:0051000 | BP | GO:0051000 | positive regulation of nitric-oxide synthase activity | 1/75 | 19/28564 | 0.048741418 | 0.048741418 | 0.116347068 | Apoe | 1 |
| GO:0071514 | BP | GO:0071514 | genomic imprinting | 1/75 | 19/28564 | 0.048741418 | 0.048741418 | 0.116347068 | Ndn | 1 |
| GO:1902430 | BP | GO:1902430 | negative regulation of amyloid-beta formation | 1/75 | 19/28564 | 0.048741418 | 0.048741418 | 0.116347068 | Apoe | 1 |
| GO:0003158 | BP | GO:0003158 | endothelium development | 2/75 | 135/28564 | 0.049183655 | 0.049183655 | 0.116347068 | Ednrb/Fzd1 | 2 |
| GO:0009791 | BP | GO:0009791 | post-embryonic development | 2/75 | 136/28564 | 0.049834873 | 0.049834873 | 0.116347068 | Sema3c/Ndn | 2 |
| GO:0097449 | CC | GO:0097449 | astrocyte projection | 3/77 | 22/28585 | 2.78829005963571E-05 | 2.78829005963571E-05 | 0.003963728 | Slc1a2/Atp1b2/Gfap | 3 |
| GO:0044306 | CC | GO:0044306 | neuron projection terminus | 5/77 | 224/28585 | 0.000352979 | 0.000352979 | 0.01672007 | Slc32a1/Slc1a2/Drd1/Scgn/Pdyn | 5 |
| GO:0098982 | CC | GO:0098982 | GABA-ergic synapse | 4/77 | 134/28585 | 0.000479381 | 0.000479381 | 0.017030626 | Slc32a1/Drd1/Epha3/Ptpro | 4 |
| GO:0044391 | CC | GO:0044391 | ribosomal subunit | 4/77 | 201/28585 | 0.002151253 | 0.002151253 | 0.04632343 | Mrpl41/Rplp1/Mrps2/Rps4x | 4 |
| GO:0043679 | CC | GO:0043679 | axon terminus | 4/77 | 202/28585 | 0.002190282 | 0.002190282 | 0.04632343 | Slc32a1/Drd1/Scgn/Pdyn | 4 |
| GO:0043197 | CC | GO:0043197 | dendritic spine | 4/77 | 206/28585 | 0.002351294 | 0.002351294 | 0.04632343 | Slc1a2/Arc/Drd1/Ptpro | 4 |
| GO:0044309 | CC | GO:0044309 | neuron spine | 4/77 | 212/28585 | 0.002607838 | 0.002607838 | 0.04632343 | Slc1a2/Arc/Drd1/Ptpro | 4 |
| GO:0045121 | CC | GO:0045121 | membrane raft | 5/77 | 390/28585 | 0.00407649 | 0.00407649 | 0.052009883 | Slc1a2/Arc/Slc2a1/Drd1/Ednrb | 5 |
| GO:0150034 | CC | GO:0150034 | distal axon | 5/77 | 397/28585 | 0.004393559 | 0.004393559 | 0.052009883 | Frmd7/Slc32a1/Drd1/Scgn/Pdyn | 5 |
| GO:0042383 | CC | GO:0042383 | sarcolemma | 3/77 | 166/28585 | 0.010277029 | 0.010277029 | 0.105045149 | Kcnj3/Slc2a1/Cd59b | 3 |
| GO:0005771 | CC | GO:0005771 | multivesicular body | 2/77 | 59/28585 | 0.01109656 | 0.01109656 | 0.105045149 | Apoe/Cst3 | 2 |
| GO:0031253 | CC | GO:0031253 | cell projection membrane | 4/77 | 325/28585 | 0.011589717 | 0.011589717 | 0.105045149 | Slc1a2/Atp1b2/Drd1/Fam107a | 4 |
| GO:0001917 | CC | GO:0001917 | photoreceptor inner segment | 2/77 | 61/28585 | 0.011827306 | 0.011827306 | 0.105045149 | Atp1b2/Slc2a1 | 2 |
| GO:0045211 | CC | GO:0045211 | postsynaptic membrane | 4/77 | 348/28585 | 0.014567912 | 0.014567912 | 0.121225247 | Arc/Drd1/Pcdh8/Ptpro | 4 |
| GO:0043198 | CC | GO:0043198 | dendritic shaft | 2/77 | 71/28585 | 0.015785274 | 0.015785274 | 0.121225247 | Slc1a2/Drd1 | 2 |
| GO:0005761 | CC | GO:0005761 | mitochondrial ribosome | 2/77 | 93/28585 | 0.02616377 | 0.02616377 | 0.124753547 | Mrpl41/Mrps2 | 2 |
| GO:0042734 | CC | GO:0042734 | presynaptic membrane | 3/77 | 238/28585 | 0.02658845 | 0.02658845 | 0.124753547 | Kcnj3/Slc1a2/Drd1 | 3 |
| GO:0001939 | CC | GO:0001939 | female pronucleus | 1/77 | 10/28585 | 0.026617153 | 0.026617153 | 0.124753547 | Slc2a1 | 1 |
| GO:0051286 | CC | GO:0051286 | cell tip | 1/77 | 10/28585 | 0.026617153 | 0.026617153 | 0.124753547 | Slc32a1 | 1 |
| GO:0005901 | CC | GO:0005901 | caveola | 2/77 | 97/28585 | 0.0282847 | 0.0282847 | 0.124753547 | Slc2a1/Drd1 | 2 |
| GO:0005890 | CC | GO:0005890 | sodium:potassium-exchanging ATPase complex | 1/77 | 11/28585 | 0.029240092 | 0.029240092 | 0.124753547 | Atp1b2 | 1 |
| GO:0031965 | CC | GO:0031965 | nuclear membrane | 3/77 | 253/28585 | 0.031034451 | 0.031034451 | 0.124753547 | Cst3/Epha3/Ednrb | 3 |
| GO:0036019 | CC | GO:0036019 | endolysosome | 1/77 | 12/28585 | 0.031858055 | 0.031858055 | 0.124753547 | Smpd1 | 1 |
| GO:0042627 | CC | GO:0042627 | chylomicron | 1/77 | 12/28585 | 0.031858055 | 0.031858055 | 0.124753547 | Apoe | 1 |
| GO:0098574 | CC | GO:0098574 | cytoplasmic side of lysosomal membrane | 1/77 | 12/28585 | 0.031858055 | 0.031858055 | 0.124753547 | Gfap | 1 |
| GO:0008021 | CC | GO:0008021 | synaptic vesicle | 3/77 | 258/28585 | 0.032609161 | 0.032609161 | 0.124753547 | Slc32a1/Wfs1/Pdyn | 3 |
| GO:0034774 | CC | GO:0034774 | secretory granule lumen | 1/77 | 14/28585 | 0.037067124 | 0.037067124 | 0.124753547 | Pdyn | 1 |
| GO:0044754 | CC | GO:0044754 | autolysosome | 1/77 | 14/28585 | 0.037067124 | 0.037067124 | 0.124753547 | Ftl1 | 1 |
| GO:0060205 | CC | GO:0060205 | cytoplasmic vesicle lumen | 1/77 | 14/28585 | 0.037067124 | 0.037067124 | 0.124753547 | Pdyn | 1 |
| GO:0042599 | CC | GO:0042599 | lamellar body | 1/77 | 16/28585 | 0.042250506 | 0.042250506 | 0.124753547 | Smpd1 | 1 |
| GO:0015629 | CC | GO:0015629 | actin cytoskeleton | 4/77 | 500/28585 | 0.046134686 | 0.046134686 | 0.124753547 | Arc/Slc2a1/Fam107a/Epha3 | 4 |
| GO:0098845 | CC | GO:0098845 | postsynaptic endosome | 1/77 | 18/28585 | 0.047406348 | 0.047406348 | 0.124753547 | Arc | 1 |
| GO:0003735 | MF | GO:0003735 | structural constituent of ribosome | 4/74 | 167/28171 | 0.000992495 | 0.000992495 | 0.084272676 | Mrpl41/Rplp1/Mrps2/Rps4x | 4 |
| GO:0016500 | MF | GO:0016500 | protein-hormone receptor activity | 2/74 | 20/28171 | 0.00125432 | 0.00125432 | 0.084272676 | Lgr6/Lgr5 | 2 |
| GO:0001664 | MF | GO:0001664 | G protein-coupled receptor binding | 5/74 | 328/28171 | 0.001734063 | 0.001734063 | 0.084272676 | Drd1/S1pr1/Ednrb/Fzd1/Pdyn | 5 |
| GO:0051117 | MF | GO:0051117 | ATPase binding | 3/74 | 102/28171 | 0.00247936 | 0.00247936 | 0.084272676 | Wfs1/Atp1b2/Drd1 | 3 |
| GO:0042277 | MF | GO:0042277 | peptide binding | 5/74 | 361/28171 | 0.002627239 | 0.002627239 | 0.084272676 | Apoe/Cst3/Vipr2/Gpr37l1/Ednrb | 5 |
| GO:0022853 | MF | GO:0022853 | active monoatomic ion transmembrane transporter activity | 4/74 | 225/28171 | 0.002945049 | 0.002945049 | 0.084272676 | Slc41a3/Slc32a1/Slc1a2/Atp1b2 | 4 |
| GO:0017147 | MF | GO:0017147 | Wnt-protein binding | 2/74 | 31/28171 | 0.003012975 | 0.003012975 | 0.084272676 | Ptpro/Fzd1 | 2 |
| GO:0008528 | MF | GO:0008528 | G protein-coupled peptide receptor activity | 3/74 | 140/28171 | 0.006019864 | 0.006019864 | 0.126334438 | Vipr2/Gpr37l1/Ednrb | 3 |
| GO:0015175 | MF | GO:0015175 | neutral L-amino acid transmembrane transporter activity | 2/74 | 46/28171 | 0.006538051 | 0.006538051 | 0.126334438 | Slc32a1/Slc1a2 | 2 |
| GO:0001653 | MF | GO:0001653 | peptide receptor activity | 3/74 | 149/28171 | 0.007145643 | 0.007145643 | 0.126334438 | Vipr2/Gpr37l1/Ednrb | 3 |
| GO:0015081 | MF | GO:0015081 | sodium ion transmembrane transporter activity | 3/74 | 158/28171 | 0.008388335 | 0.008388335 | 0.126334438 | Slc41a3/Slc1a2/Atp1b2 | 3 |
| GO:0008237 | MF | GO:0008237 | metallopeptidase activity | 3/74 | 201/28171 | 0.01600975 | 0.01600975 | 0.15490281 | Cpa6/Adamts19/Mmel1 | 3 |
| GO:0042562 | MF | GO:0042562 | hormone binding | 2/74 | 87/28171 | 0.022054524 | 0.022054524 | 0.15490281 | Vipr2/Ednrb | 2 |
| GO:0004725 | MF | GO:0004725 | protein tyrosine phosphatase activity | 2/74 | 91/28171 | 0.023980055 | 0.023980055 | 0.15490281 | Ptpn14/Ptpro | 2 |
| GO:0008514 | MF | GO:0008514 | organic anion transmembrane transporter activity | 3/74 | 238/28171 | 0.024878832 | 0.024878832 | 0.15490281 | Slc32a1/Slc1a2/Slc2a1 | 3 |
| GO:0005041 | MF | GO:0005041 | low-density lipoprotein particle receptor activity | 1/74 | 10/28171 | 0.025963905 | 0.025963905 | 0.15490281 | Lrp10 | 1 |
| GO:0005391 | MF | GO:0005391 | P-type sodium:potassium-exchanging transporter activity | 1/74 | 10/28171 | 0.025963905 | 0.025963905 | 0.15490281 | Atp1b2 | 1 |
| GO:0008020 | MF | GO:0008020 | G protein-coupled photoreceptor activity | 1/74 | 10/28171 | 0.025963905 | 0.025963905 | 0.15490281 | Gpr88 | 1 |
| GO:0016742 | MF | GO:0016742 | hydroxymethyl-, formyl- and related transferase activity | 1/74 | 10/28171 | 0.025963905 | 0.025963905 | 0.15490281 | Aldh1l1 | 1 |
| GO:0005005 | MF | GO:0005005 | transmembrane-ephrin receptor activity | 1/74 | 11/28171 | 0.028523426 | 0.028523426 | 0.15490281 | Epha3 | 1 |
| GO:0008174 | MF | GO:0008174 | mRNA methyltransferase activity | 1/74 | 11/28171 | 0.028523426 | 0.028523426 | 0.15490281 | Trmt61a | 1 |
| GO:0009881 | MF | GO:0009881 | photoreceptor activity | 1/74 | 12/28171 | 0.031076312 | 0.031076312 | 0.15490281 | Gpr88 | 1 |
| GO:1901338 | MF | GO:1901338 | catecholamine binding | 1/74 | 12/28171 | 0.031076312 | 0.031076312 | 0.15490281 | Drd1 | 1 |
| GO:0002161 | MF | GO:0002161 | aminoacyl-tRNA editing activity | 1/74 | 13/28171 | 0.03362258 | 0.03362258 | 0.15490281 | Lars2 | 1 |
| GO:0016832 | MF | GO:0016832 | aldehyde-lyase activity | 1/74 | 13/28171 | 0.03362258 | 0.03362258 | 0.15490281 | Aldoc | 1 |
| GO:0005313 | MF | GO:0005313 | L-glutamate transmembrane transporter activity | 1/74 | 15/28171 | 0.038695328 | 0.038695328 | 0.15490281 | Slc1a2 | 1 |
| GO:0015355 | MF | GO:0015355 | secondary active monocarboxylate transmembrane transporter activity | 1/74 | 16/28171 | 0.041221843 | 0.041221843 | 0.15490281 | Slc32a1 | 1 |
| GO:0016646 | MF | GO:0016646 | oxidoreductase activity, acting on the CH-NH group of donors, NAD or NADP as | 1/74 | 16/28171 | 0.041221843 | 0.041221843 | 0.15490281 | Aldh1l1 | 1 |
| GO:0016722 | MF | GO:0016722 | oxidoreductase activity, acting on metal ions | 1/74 | 16/28171 | 0.041221843 | 0.041221843 | 0.15490281 | Hephl1 | 1 |
| GO:0042813 | MF | GO:0042813 | Wnt receptor activity | 1/74 | 16/28171 | 0.041221843 | 0.041221843 | 0.15490281 | Fzd1 | 1 |
| GO:0045125 | MF | GO:0045125 | bioactive lipid receptor activity | 1/74 | 16/28171 | 0.041221843 | 0.041221843 | 0.15490281 | S1pr1 | 1 |
| GO:0070403 | MF | GO:0070403 | NAD+ binding | 1/74 | 16/28171 | 0.041221843 | 0.041221843 | 0.15490281 | Hadh | 1 |
| GO:1901681 | MF | GO:1901681 | sulfur compound binding | 3/74 | 293/28171 | 0.042054039 | 0.042054039 | 0.15490281 | Apoe/Ltbp2/Ftl1 | 3 |
| GO:0140101 | MF | GO:0140101 | catalytic activity, acting on a tRNA | 2/74 | 126/28171 | 0.043496187 | 0.043496187 | 0.15490281 | Lars2/Trmt61a | 2 |
| GO:0004029 | MF | GO:0004029 | aldehyde dehydrogenase (NAD+) activity | 1/74 | 17/28171 | 0.043741807 | 0.043741807 | 0.15490281 | Aldh1l1 | 1 |
| GO:0099508 | MF | GO:0099508 | voltage-gated monoatomic ion channel activity involved in regulation of presynapti | 1/74 | 17/28171 | 0.043741807 | 0.043741807 | 0.15490281 | Kcnj3 | 1 |
| GO:0008199 | MF | GO:0008199 | ferric iron binding | 1/74 | 18/28171 | 0.046255237 | 0.046255237 | 0.15490281 | Ftl1 | 1 |
| GO:0005324 | MF | GO:0005324 | long-chain fatty acid transporter activity | 1/74 | 19/28171 | 0.048762149 | 0.048762149 | 0.15490281 | Slc2a1 | 1 |
| GO:0140693 | MF | GO:0140693 | molecular condensate scaffold activity | 1/74 | 19/28171 | 0.048762149 | 0.048762149 | 0.15490281 | Ubqln2 | 1 |

Cre+_vs_Cre-_enrichGO

| | ONTOLOGY | ID | Description | GeneRati | BgRatio | pvalue | p.adjust | qvalue | geneID | Count |
|---|---|---|---|---|---|---|---|---|---|---|
| GO:009712 | BP | GO:009712 | receptor localization to synapse | 3/19 | 86/28564 | ################ | ################ | 0.00821527 | Kif5a/Camk2a/Nptx | 3 |
| GO:009963 | BP | GO:009963 | protein localization to postsynaptic specialization membrane | 2/19 | 41/28564 | 0.000338453 | 0.000338453 | 0.026330415 | Camk2a/Nptxr | 2 |
| GO:009964 | BP | GO:009964 | neurotransmitter receptor localization to postsynaptic specialization m | 2/19 | 41/28564 | 0.000338453 | 0.000338453 | 0.026330415 | Camk2a/Nptxr | 2 |
| GO:000761 | BP | GO:000761 | long-term memory | 2/19 | 43/28564 | 0.000372415 | 0.000372415 | 0.026330415 | Apoe/Camk2n1 | 2 |
| GO:003150 | BP | GO:003150 | protein-containing complex localization | 3/19 | 239/2856 | 0.000507651 | 0.000507651 | 0.030764434 | Kif5a/Camk2a/Nptx | 3 |
| GO:005507 | BP | GO:005507 | calcium ion homeostasis | 3/19 | 347/2856 | 0.001490597 | 0.001490597 | 0.056689707 | Apoe/Camk2n1/Calr | 3 |
| GO:000657 | BP | GO:000657 | cellular modified amino acid metabolic process | 2/19 | 128/2856 | 0.003241169 | 0.003241169 | 0.056689707 | Aldh1l1/Ctsb | 2 |
| GO:009907 | BP | GO:009907 | regulation of postsynaptic membrane neurotransmitter receptor levels | 2/19 | 133/2856 | 0.003493449 | 0.003493449 | 0.056689707 | Camk2a/Nptxr | 2 |
| GO:006099 | BP | GO:006099 | dendritic spine development | 2/19 | 134/2856 | 0.003544977 | 0.003544977 | 0.056689707 | Apoe/Camk2a | 2 |
| GO:005192 | BP | GO:005192 | positive regulation of calcium ion transport | 2/19 | 150/2856 | 0.004417596 | 0.004417596 | 0.056689707 | Calm3/Camk2a | 2 |
| GO:003437 | BP | GO:003437 | very-low-density lipoprotein particle remodeling | 1/19 | 10/28564 | 0.006632896 | 0.006632896 | 0.056689707 | Apoe | 1 |
| GO:200064 | BP | GO:200064 | positive regulation of receptor catabolic process | 1/19 | 10/28564 | 0.006632896 | 0.006632896 | 0.056689707 | Apoe | 1 |
| GO:000551 | BP | GO:000551 | detection of calcium ion | 1/19 | 11/28564 | 0.007293888 | 0.007293888 | 0.056689707 | Calm3 | 1 |
| GO:003438 | BP | GO:003438 | high-density lipoprotein particle clearance | 1/19 | 11/28564 | 0.007293888 | 0.007293888 | 0.056689707 | Apoe | 1 |
| GO:004356 | BP | GO:004356 | positive regulation of insulin-like growth factor receptor signaling path | 1/19 | 11/28564 | 0.007293888 | 0.007293888 | 0.056689707 | Myorg | 1 |
| GO:009964 | BP | GO:009964 | axo-dendritic protein transport | 1/19 | 11/28564 | 0.007293888 | 0.007293888 | 0.056689707 | Kif5a | 1 |
| GO:004352 | BP | GO:004352 | negative regulation of neuron apoptotic process | 2/19 | 199/2856 | 0.00763864 | 0.00763864 | 0.056689707 | Apoe/Pcdhgc3 | 2 |
| GO:001703 | BP | GO:001703 | protein import | 1/19 | 12/28564 | 0.007954464 | 0.007954464 | 0.056689707 | Apoe | 1 |
| GO:009020 | BP | GO:009020 | negative regulation of triglyceride metabolic process | 1/19 | 12/28564 | 0.007954464 | 0.007954464 | 0.056689707 | Apoe | 1 |
| GO:009709 | BP | GO:009709 | presynaptic membrane organization | 1/19 | 12/28564 | 0.007954464 | 0.007954464 | 0.056689707 | Apoe | 1 |
| GO:009951 | BP | GO:009951 | dense core granule cytoskeletal transport | 1/19 | 12/28564 | 0.007954464 | 0.007954464 | 0.056689707 | Kif5a | 1 |
| GO:190027 | BP | GO:190027 | negative regulation of long-term synaptic potentiation | 1/19 | 12/28564 | 0.007954464 | 0.007954464 | 0.056689707 | Apoe | 1 |
| GO:190294 | BP | GO:190294 | regulation of tau-protein kinase activity | 1/19 | 12/28564 | 0.007954464 | 0.007954464 | 0.056689707 | Apoe | 1 |
| GO:190590 | BP | GO:190590 | negative regulation of amyloid fibril formation | 1/19 | 12/28564 | 0.007954464 | 0.007954464 | 0.056689707 | Apoe | 1 |
| GO:000646 | BP | GO:000646 | negative regulation of protein kinase activity | 2/19 | 206/2856 | 0.008164285 | 0.008164285 | 0.056689707 | Apoe/Camk2n1 | 2 |
| GO:000751 | BP | GO:000751 | skeletal muscle tissue development | 2/19 | 211/2856 | 0.008549512 | 0.008549512 | 0.056689707 | Meg3/Myorg | 2 |
| GO:009962 | BP | GO:009962 | neurotransmitter receptor diffusion trapping | 1/19 | 13/28564 | 0.008614623 | 0.008614623 | 0.056689707 | Camk2a | 1 |
| GO:000741 | BP | GO:000741 | synapse assembly | 2/19 | 216/2856 | 0.008942818 | 0.008942818 | 0.056689707 | Lingo2/Nptxr | 2 |
| GO:001608 | BP | GO:001608 | synaptic vesicle docking | 1/19 | 14/28564 | 0.009274366 | 0.009274366 | 0.056689707 | Camk2a | 1 |
| GO:003225 | BP | GO:003225 | dense core granule localization | 1/19 | 14/28564 | 0.009274366 | 0.009274366 | 0.056689707 | Kif5a | 1 |
| GO:009711 | BP | GO:009711 | AMPA glutamate receptor clustering | 1/19 | 14/28564 | 0.009274366 | 0.009274366 | 0.056689707 | Apoe | 1 |
| GO:009768 | BP | GO:009768 | glutamate receptor clustering | 1/19 | 14/28564 | 0.009274366 | 0.009274366 | 0.056689707 | Apoe | 1 |
| GO:009893 | BP | GO:009893 | dendritic transport | 1/19 | 14/28564 | 0.009274366 | 0.009274366 | 0.056689707 | Kif5a | 1 |
| GO:009900 | BP | GO:009900 | calmodulin dependent kinase signaling pathway | 1/19 | 14/28564 | 0.009274366 | 0.009274366 | 0.056689707 | Camk2a | 1 |
| GO:190184 | BP | GO:190184 | regulation of high voltage-gated calcium channel activity | 1/19 | 14/28564 | 0.009274366 | 0.009274366 | 0.056689707 | Calm3 | 1 |
| GO:190293 | BP | GO:190293 | negative regulation of alcohol biosynthetic process | 1/19 | 14/28564 | 0.009274366 | 0.009274366 | 0.056689707 | Apoe | 1 |
| GO:200082 | BP | GO:200082 | regulation of behavioral fear response | 1/19 | 14/28564 | 0.009274366 | 0.009274366 | 0.056689707 | Apoe | 1 |
| GO:200114 | BP | GO:200114 | positive regulation of phospholipid transport | 1/19 | 14/28564 | 0.009274366 | 0.009274366 | 0.056689707 | Apoe | 1 |
| GO:003237 | BP | GO:003237 | negative regulation of sterol transport | 1/19 | 15/28564 | 0.009933693 | 0.009933693 | 0.056689707 | Apoe | 1 |
| GO:003237 | BP | GO:003237 | negative regulation of cholesterol transport | 1/19 | 15/28564 | 0.009933693 | 0.009933693 | 0.056689707 | Apoe | 1 |
| GO:190580 | BP | GO:190580 | negative regulation of synapse organization | 1/19 | 15/28564 | 0.009933693 | 0.009933693 | 0.056689707 | Apoe | 1 |
| GO:003367 | BP | GO:003367 | negative regulation of kinase activity | 2/19 | 230/2856 | 0.010086614 | 0.010086614 | 0.056689707 | Apoe/Camk2n1 | 2 |
| GO:001569 | BP | GO:001569 | magnesium ion transport | 1/19 | 16/28564 | 0.010592605 | 0.010592605 | 0.056689707 | Slc41a3 | 1 |
| GO:003248 | BP | GO:003248 | Cdc42 protein signal transduction | 1/19 | 16/28564 | 0.010592605 | 0.010592605 | 0.056689707 | Apoe | 1 |
| GO:003370 | BP | GO:003370 | phospholipid efflux | 1/19 | 16/28564 | 0.010592605 | 0.010592605 | 0.056689707 | Apoe | 1 |
| GO:004215 | BP | GO:004215 | lipoprotein catabolic process | 1/19 | 16/28564 | 0.010592605 | 0.010592605 | 0.056689707 | Apoe | 1 |
| GO:004364 | BP | GO:004364 | dicarboxylic acid catabolic process | 1/19 | 16/28564 | 0.010592605 | 0.010592605 | 0.056689707 | Aldh1l1 | 1 |
| GO:005104 | BP | GO:005104 | positive regulation of membrane protein ectodomain proteolysis | 1/19 | 16/28564 | 0.010592605 | 0.010592605 | 0.056689707 | Apoe | 1 |
| GO:190295 | BP | GO:190295 | regulation of dendritic spine maintenance | 1/19 | 16/28564 | 0.010592605 | 0.010592605 | 0.056689707 | Apoe | 1 |
| GO:005165 | BP | GO:005165 | maintenance of location in cell | 2/19 | 242/2856 | 0.011116241 | 0.011116241 | 0.056689707 | Apoe/Calm3 | 2 |
| GO:003816 | BP | GO:003816 | angiotensin-activated signaling pathway | 1/19 | 17/28564 | 0.011251101 | 0.011251101 | 0.056689707 | Camk2a | 1 |
| GO:005508 | BP | GO:005508 | fatty acid homeostasis | 1/19 | 17/28564 | 0.011251101 | 0.011251101 | 0.056689707 | Apoe | 1 |
| GO:190022 | BP | GO:190022 | regulation of amyloid-beta clearance | 1/19 | 18/28564 | 0.011909181 | 0.011909181 | 0.056689707 | Apoe | 1 |
| GO:009917 | BP | GO:009917 | postsynapse organization | 2/19 | 251/2856 | 0.011917832 | 0.011917832 | 0.056689707 | Apoe/Nptxr | 2 |
| GO:001905 | BP | GO:001905 | viral life cycle | 2/19 | 257/2856 | 0.012466049 | 0.012466049 | 0.056689707 | Apoe/Ctsb | 2 |
| GO:000659 | BP | GO:000659 | thyroid hormone generation | 1/19 | 19/28564 | 0.012566847 | 0.012566847 | 0.056689707 | Ctsb | 1 |
| GO:001054 | BP | GO:001054 | negative regulation of platelet activation | 1/19 | 19/28564 | 0.012566847 | 0.012566847 | 0.056689707 | Apoe | 1 |
| GO:003251 | BP | GO:003251 | positive regulation of phosphoprotein phosphatase activity | 1/19 | 19/28564 | 0.012566847 | 0.012566847 | 0.056689707 | Calm3 | 1 |
| GO:005100 | BP | GO:005100 | positive regulation of nitric-oxide synthase activity | 1/19 | 19/28564 | 0.012566847 | 0.012566847 | 0.056689707 | Apoe | 1 |
| GO:007151 | BP | GO:007151 | genomic imprinting | 1/19 | 19/28564 | 0.012566847 | 0.012566847 | 0.056689707 | Meg3 | 1 |
| GO:190243 | BP | GO:190243 | negative regulation of amyloid-beta formation | 1/19 | 19/28564 | 0.012566847 | 0.012566847 | 0.056689707 | Apoe | 1 |
| GO:003437 | BP | GO:003437 | plasma lipoprotein particle assembly | 1/19 | 20/28564 | 0.013224098 | 0.013224098 | 0.056689707 | Apoe | 1 |
| GO:003564 | BP | GO:003564 | locomotory exploration behavior | 1/19 | 20/28564 | 0.013224098 | 0.013224098 | 0.056689707 | Apoe | 1 |
| GO:004311 | BP | GO:004311 | negative regulation of vascular permeability | 1/19 | 20/28564 | 0.013224098 | 0.013224098 | 0.056689707 | Apoe | 1 |
| GO:004369 | BP | GO:004369 | reverse cholesterol transport | 1/19 | 20/28564 | 0.013224098 | 0.013224098 | 0.056689707 | Apoe | 1 |
| GO:006031 | BP | GO:006031 | regulation of ryanodine-sensitive calcium-release channel activity | 1/19 | 20/28564 | 0.013224098 | 0.013224098 | 0.056689707 | Calm3 | 1 |
| GO:005134 | BP | GO:005134 | negative regulation of transferase activity | 2/19 | 269/2856 | 0.013595265 | 0.013595265 | 0.056689707 | Apoe/Camk2n1 | 2 |
| GO:004479 | BP | GO:004479 | positive regulation by host of viral process | 1/19 | 21/28564 | 0.013880934 | 0.013880934 | 0.056689707 | Apoe | 1 |
| GO:004554 | BP | GO:004554 | regulation of cholesterol biosynthetic process | 1/19 | 21/28564 | 0.013880934 | 0.013880934 | 0.056689707 | Apoe | 1 |
| GO:009008 | BP | GO:009008 | regulation of inclusion body assembly | 1/19 | 21/28564 | 0.013880934 | 0.013880934 | 0.056689707 | Apoe | 1 |
| GO:010611 | BP | GO:010611 | regulation of sterol biosynthetic process | 1/19 | 21/28564 | 0.013880934 | 0.013880934 | 0.056689707 | Apoe | 1 |
| GO:015010 | BP | GO:015010 | transport across blood-brain barrier | 1/19 | 21/28564 | 0.013880934 | 0.013880934 | 0.056689707 | Apoe | 1 |
| GO:190121 | BP | GO:190121 | negative regulation of neuron death | 2/19 | 273/2856 | 0.01398129 | 0.01398129 | 0.056689707 | Apoe/Pcdhgc3 | 2 |
| GO:000674 | BP | GO:000674 | NADPH regeneration | 1/19 | 22/28564 | 0.014537357 | 0.014537357 | 0.056689707 | Aldh1l1 | 1 |
| GO:009020 | BP | GO:009020 | positive regulation of cholesterol metabolic process | 1/19 | 22/28564 | 0.014537357 | 0.014537357 | 0.056689707 | Apoe | 1 |
| GO:009896 | BP | GO:009896 | regulation of postsynaptic neurotransmitter receptor activity | 1/19 | 23/28564 | 0.015193365 | 0.015193365 | 0.056689707 | Nptxr | 1 |
| GO:004669 | BP | GO:004669 | decidualization | 1/19 | 24/28564 | 0.01584896 | 0.01584896 | 0.056689707 | Ctsb | 1 |
| GO:000670 | BP | GO:000670 | steroid catabolic process | 1/19 | 25/28564 | 0.016504141 | 0.016504141 | 0.056689707 | Apoe | 1 |
| GO:001066 | BP | GO:001066 | positive regulation of cardiac muscle cell apoptotic process | 1/19 | 25/28564 | 0.016504141 | 0.016504141 | 0.056689707 | Camk2a | 1 |
| GO:001089 | BP | GO:001089 | negative regulation of steroid biosynthetic process | 1/19 | 25/28564 | 0.016504141 | 0.016504141 | 0.056689707 | Apoe | 1 |
| GO:006500 | BP | GO:006500 | protein-lipid complex assembly | 1/19 | 25/28564 | 0.016504141 | 0.016504141 | 0.056689707 | Apoe | 1 |
| GO:000308 | BP | GO:000308 | positive regulation of systemic arterial blood pressure | 1/19 | 26/28564 | 0.017158909 | 0.017158909 | 0.056689707 | Camk2n1 | 1 |
| GO:000673 | BP | GO:000673 | one-carbon metabolic process | 1/19 | 26/28564 | 0.017158909 | 0.017158909 | 0.056689707 | Aldh1l1 | 1 |
| GO:004593 | BP | GO:004593 | negative regulation of steroid metabolic process | 1/19 | 26/28564 | 0.017158909 | 0.017158909 | 0.056689707 | Apoe | 1 |
| GO:199077 | BP | GO:199077 | response to angiotensin | 1/19 | 26/28564 | 0.017158909 | 0.017158909 | 0.056689707 | Camk2a | 1 |
| GO:003436 | BP | GO:003436 | protein-lipid complex remodeling | 1/19 | 27/28564 | 0.017813264 | 0.017813264 | 0.056721799 | Apoe | 1 |
| GO:005080 | BP | GO:005080 | regulation of synapse structure or activity | 2/19 | 313/2856 | 0.018099624 | 0.018099624 | 0.056721799 | Apoe/Lingo2 | 2 |
| GO:000687 | BP | GO:000687 | intracellular calcium ion homeostasis | 2/19 | 316/2856 | 0.018427022 | 0.018427022 | 0.056721799 | Apoe/Calm3 | 2 |
| GO:000726 | BP | GO:000726 | nitric oxide mediated signal transduction | 1/19 | 28/28564 | 0.018467206 | 0.018467206 | 0.056721799 | Apoe | 1 |
| GO:009706 | BP | GO:009706 | dendritic spine maintenance | 1/19 | 28/28564 | 0.018467206 | 0.018467206 | 0.056721799 | Apoe | 1 |
| GO:007084 | BP | GO:007084 | inclusion body assembly | 1/19 | 29/28564 | 0.019120736 | 0.019120736 | 0.056721799 | Apoe | 1 |
| GO:009890 | BP | GO:009890 | regulation of cardiac muscle cell action potential | 1/19 | 29/28564 | 0.019120736 | 0.019120736 | 0.056721799 | Calm3 | 1 |
| GO:199000 | BP | GO:199000 | amyloid fibril formation | 1/19 | 29/28564 | 0.019120736 | 0.019120736 | 0.056721799 | Apoe | 1 |
| GO:005070 | BP | GO:005070 | regulation of protein secretion | 2/19 | 336/2856 | 0.020673882 | 0.020673882 | 0.058828098 | Apoe/Camk2n1 | 2 |
| GO:005140 | BP | GO:005140 | neuron apoptotic process | 2/19 | 337/2856 | 0.020789124 | 0.020789124 | 0.058828098 | Apoe/Pcdhgc3 | 2 |
| GO:007029 | BP | GO:007029 | sarcoplasmic reticulum calcium ion transport | 1/19 | 32/28564 | 0.021078852 | 0.021078852 | 0.058828098 | Calm3 | 1 |
| GO:001092 | BP | GO:001092 | positive regulation of phosphatase activity | 1/19 | 33/28564 | 0.021730734 | 0.021730734 | 0.059747021 | Calm3 | 1 |
| GO:000834 | BP | GO:000834 | determination of adult lifespan | 1/19 | 34/28564 | 0.022382205 | 0.022382205 | 0.059747021 | Meg3 | 1 |
| GO:005099 | BP | GO:005099 | regulation of nitric-oxide synthase activity | 1/19 | 34/28564 | 0.022382205 | 0.022382205 | 0.059747021 | Apoe | 1 |
| GO:004428 | BP | GO:004428 | small molecule catabolic process | 2/19 | 355/2856 | 0.022909871 | 0.022909871 | 0.059747021 | Aldh1l1/Apoe | 2 |
| GO:005509 | BP | GO:005509 | response to lipoprotein particle | 1/19 | 35/28564 | 0.023033265 | 0.023033265 | 0.059747021 | Apoe | 1 |
| GO:004353 | BP | GO:004353 | negative regulation of blood vessel endothelial cell migration | 1/19 | 36/28564 | 0.023683914 | 0.023683914 | 0.059747021 | Apoe | 1 |
| GO:009724 | BP | GO:009724 | amyloid-beta clearance | 1/19 | 37/28564 | 0.024334153 | 0.024334153 | 0.059747021 | Apoe | 1 |
| GO:005134 | BP | GO:005134 | negative regulation of hydrolase activity | 2/19 | 370/2856 | 0.02474326 | 0.02474326 | 0.059747021 | Camk2a/Ctsb | 2 |
| GO:001633 | BP | GO:001633 | calcium-dependent cell-cell adhesion via plasma membrane cell adh | 1/19 | 38/28564 | 0.024983981 | 0.024983981 | 0.059747021 | Pcdhgc3 | 1 |
| GO:007140 | BP | GO:007140 | cellular response to lipoprotein particle stimulus | 1/19 | 38/28564 | 0.024983981 | 0.024983981 | 0.059747021 | Apoe | 1 |
| GO:005072 | BP | GO:005072 | regulation of inflammatory response | 2/19 | 377/2856 | 0.02561903 | 0.02561903 | 0.059747021 | Apoe/Camk2n1 | 2 |
| GO:000676 | BP | GO:000676 | water-soluble vitamin metabolic process | 1/19 | 39/28564 | 0.025633399 | 0.025633399 | 0.059747021 | Aldh1l1 | 1 |
| GO:001993 | BP | GO:001993 | cGMP-mediated signaling | 1/19 | 39/28564 | 0.025633399 | 0.025633399 | 0.059747021 | Apoe | 1 |
| GO:003243 | BP | GO:003243 | melanosome organization | 1/19 | 39/28564 | 0.025633399 | 0.025633399 | 0.059747021 | Apoe | 1 |
| GO:004616 | BP | GO:004616 | alcohol catabolic process | 1/19 | 39/28564 | 0.025633399 | 0.025633399 | 0.059747021 | Apoe | 1 |
| GO:005509 | BP | GO:005509 | acylglycerol homeostasis | 1/19 | 39/28564 | 0.025633399 | 0.025633399 | 0.059747021 | Apoe | 1 |
| GO:007032 | BP | GO:007032 | triglyceride homeostasis | 1/19 | 39/28564 | 0.025633399 | 0.025633399 | 0.059747021 | Apoe | 1 |
| GO:005196 | BP | GO:005196 | positive regulation of nervous system development | 2/19 | 379/2856 | 0.025871584 | 0.025871584 | 0.059813699 | Apoe/Lingo2 | 2 |
| GO:003057 | BP | GO:003057 | collagen catabolic process | 1/19 | 40/28564 | 0.026282408 | 0.026282408 | 0.059813699 | Ctsb | 1 |
| GO:003564 | BP | GO:003564 | exploration behavior | 1/19 | 40/28564 | 0.026282408 | 0.026282408 | 0.059813699 | Apoe | 1 |
| GO:004508 | BP | GO:004508 | regulation of innate immune response | 2/19 | 385/2856 | 0.026635424 | 0.026635424 | 0.059813699 | Apoe/Tril | 2 |
| GO:000641 | BP | GO:000641 | tRNA aminoacylation for protein translation | 1/19 | 41/28564 | 0.026931007 | 0.026931007 | 0.059813699 | Lars2 | 1 |
| GO:000202 | BP | GO:000202 | response to dietary excess | 1/19 | 43/28564 | 0.028226977 | 0.028226977 | 0.060082283 | Apoe | 1 |
| GO:004303 | BP | GO:004303 | tRNA aminoacylation | 1/19 | 44/28564 | 0.028874348 | 0.028874348 | 0.060082283 | Lars2 | 1 |
| GO:190210 | BP | GO:190210 | regulation of mitochondrial membrane permeability involved in apoptc | 1/19 | 44/28564 | 0.028874348 | 0.028874348 | 0.060082283 | Camk2a | 1 |
| GO:000650 | BP | GO:000650 | membrane protein ectodomain proteolysis | 1/19 | 45/28564 | 0.029521311 | 0.029521311 | 0.060082283 | Apoe | 1 |
| GO:000690 | BP | GO:000690 | vesicle docking involved in exocytosis | 1/19 | 45/28564 | 0.029521311 | 0.029521311 | 0.060082283 | Camk2a | 1 |
| GO:000193 | BP | GO:000193 | negative regulation of endothelial cell proliferation | 1/19 | 46/28564 | 0.030167866 | 0.030167866 | 0.060082283 | Apoe | 1 |
| GO:000293 | BP | GO:000293 | response to ischemia | 1/19 | 46/28564 | 0.030167866 | 0.030167866 | 0.060082283 | Camk2a | 1 |
| GO:004848 | BP | GO:004848 | synaptic vesicle transport | 1/19 | 46/28564 | 0.030167866 | 0.030167866 | 0.060082283 | Kif5a | 1 |
| GO:003420 | BP | GO:003420 | amyloid-beta formation | 1/19 | 48/28564 | 0.031459752 | 0.031459752 | 0.060142414 | Apoe | 1 |
| GO:003577 | BP | GO:003577 | positive regulation of insulin secretion involved in cellular response to | 1/19 | 48/28564 | 0.031459752 | 0.031459752 | 0.060142414 | Camk2n1 | 1 |
| GO:004874 | BP | GO:004874 | skeletal muscle fiber development | 1/19 | 48/28564 | 0.031459752 | 0.031459752 | 0.060142414 | Myorg | 1 |
| GO:199057 | BP | GO:199057 | potassium ion import across plasma membrane | 1/19 | 48/28564 | 0.031459752 | 0.031459752 | 0.060142414 | Kcnj3 | 1 |
| GO:000673 | BP | GO:000673 | NADP metabolic process | 1/19 | 49/28564 | 0.032105083 | 0.032105083 | 0.060142414 | Aldh1l1 | 1 |
| GO:001054 | BP | GO:001054 | regulation of platelet activation | 1/19 | 49/28564 | 0.032105083 | 0.032105083 | 0.060142414 | Apoe | 1 |
| GO:003414 | BP | GO:003414 | toll-like receptor 4 signaling pathway | 1/19 | 49/28564 | 0.032105083 | 0.032105083 | 0.060142414 | Tril | 1 |
| GO:006013 | BP | GO:006013 | maternal process involved in female pregnancy | 1/19 | 50/28564 | 0.032750007 | 0.032750007 | 0.060142414 | Ctsb | 1 |
| GO:190102 | BP | GO:190102 | positive regulation of calcium ion transmembrane transporter activity | 1/19 | 50/28564 | 0.032750007 | 0.032750007 | 0.060142414 | Calm3 | 1 |
| GO:003559 | BP | GO:003559 | establishment of protein localization to extracellular region | 2/19 | 439/2856 | 0.033915228 | 0.033915228 | 0.060781354 | Apoe/Camk2n1 | 2 |
| GO:190004 | BP | GO:190004 | negative regulation of hemostasis | 1/19 | 52/28564 | 0.034038633 | 0.034038633 | 0.060781354 | Apoe | 1 |
| GO:005081 | BP | GO:005081 | negative regulation of coagulation | 1/19 | 53/28564 | 0.034682336 | 0.034682336 | 0.060781354 | Apoe | 1 |
| GO:007169 | BP | GO:007169 | protein localization to extracellular region | 2/19 | 445/2856 | 0.034767737 | 0.034767737 | 0.060781354 | Apoe/Camk2n1 | 2 |
| GO:001490 | BP | GO:001490 | myotube cell development | 1/19 | 54/28564 | 0.035325633 | 0.035325633 | 0.060781354 | Myorg | 1 |
| GO:004477 | BP | GO:004477 | mitotic cell cycle phase transition | 2/19 | 450/2856 | 0.035484624 | 0.035484624 | 0.060781354 | Calm3/Camk2a | 2 |
| GO:003134 | BP | GO:003134 | positive regulation of defense response | 2/19 | 452/2856 | 0.035773013 | 0.035773013 | 0.060781354 | Tril/Camk2n1 | 2 |
| GO:004577 | BP | GO:004577 | positive regulation of axon extension | 1/19 | 55/28564 | 0.035968524 | 0.035968524 | 0.060781354 | Apoe | 1 |
| GO:000166 | BP | GO:000166 | behavioral fear response | 1/19 | 56/28564 | 0.036611009 | 0.036611009 | 0.060781354 | Apoe | 1 |
| GO:007099 | BP | GO:007099 | neuron death | 2/19 | 461/2856 | 0.037082236 | 0.037082236 | 0.060781354 | Apoe/Pcdhgc3 | 2 |
| GO:190265 | BP | GO:190265 | secondary alcohol biosynthetic process | 1/19 | 57/28564 | 0.037253088 | 0.037253088 | 0.060781354 | Apoe | 1 |
| GO:200122 | BP | GO:200122 | regulation of neuron migration | 1/19 | 57/28564 | 0.037253088 | 0.037253088 | 0.060781354 | Camk2a | 1 |
| GO:003305 | BP | GO:003305 | cellular pigmentation | 1/19 | 58/28564 | 0.037894761 | 0.037894761 | 0.061123029 | Apoe | 1 |
| GO:014089 | BP | GO:014089 | cell surface toll-like receptor signaling pathway | 1/19 | 58/28564 | 0.037894761 | 0.037894761 | 0.061123029 | Tril | 1 |
| GO:003545 | BP | GO:003545 | cellular response to interferon-beta | 1/19 | 59/28564 | 0.03853603 | 0.03853603 | 0.061456352 | Camk2a | 1 |
| GO:190595 | BP | GO:190595 | negative regulation of lipid localization | 1/19 | 59/28564 | 0.03853603 | 0.03853603 | 0.061456352 | Apoe | 1 |
| GO:003432 | BP | GO:003432 | cell junction assembly | 2/19 | 477/2856 | 0.039455488 | 0.039455488 | 0.061761747 | Lingo2/Nptxr | 2 |
| GO:009747 | BP | GO:009747 | synaptic vesicle localization | 1/19 | 61/28564 | 0.039817352 | 0.039817352 | 0.062099044 | Kif5a | 1 |
| GO:004259 | BP | GO:004259 | fear response | 1/19 | 62/28564 | 0.040457407 | 0.040457407 | 0.062182818 | Apoe | 1 |
| GO:000283 | BP | GO:000283 | regulation of response to biotic stimulus | 2/19 | 485/2856 | 0.040663721 | 0.040663721 | 0.062263586 | Apoe/Tril | 2 |
| GO:000275 | BP | GO:000275 | cell surface pattern recognition receptor signaling pathway | 1/19 | 63/28564 | 0.041097057 | 0.041097057 | 0.062263586 | Tril | 1 |
| GO:003476 | BP | GO:003476 | regulation of monoatomic ion transmembrane transport | 2/19 | 492/2856 | 0.041732561 | 0.041732561 | 0.062561764 | Kcnj3/Calm3 | 2 |
| GO:004231 | BP | GO:004231 | vasodilation | 1/19 | 64/28564 | 0.041736303 | 0.041736303 | 0.062561764 | Apoe | 1 |
| GO:000740 | BP | GO:000740 | axonogenesis | 2/19 | 498/2856 | 0.042657266 | 0.042657266 | 0.063271543 | Apoe/Kif5a | 2 |
| GO:001065 | BP | GO:001065 | cardiac muscle cell apoptotic process | 1/19 | 66/28564 | 0.043013585 | 0.043013585 | 0.063356998 | Camk2a | 1 |
| GO:001590 | BP | GO:001590 | long-chain fatty acid transport | 1/19 | 67/28564 | 0.04365162 | 0.04365162 | 0.063633941 | Apoe | 1 |
| GO:004828 | BP | GO:004828 | lung alveolus development | 1/19 | 67/28564 | 0.04365162 | 0.04365162 | 0.063633941 | Meg3 | 1 |
| GO:000630 | BP | GO:000630 | DNA alkylation | 1/19 | 69/28564 | 0.044926483 | 0.044926483 | 0.064604363 | Meg3 | 1 |
| GO:000630 | BP | GO:000630 | DNA methylation | 1/19 | 69/28564 | 0.044926483 | 0.044926483 | 0.064604363 | Meg3 | 1 |
| GO:003545 | BP | GO:003545 | response to interferon-beta | 1/19 | 71/28564 | 0.046199735 | 0.046199735 | 0.064681235 | Camk2a | 1 |
| GO:005511 | BP | GO:005511 | regulation of cardiac muscle contraction | 1/19 | 71/28564 | 0.046199735 | 0.046199735 | 0.064681235 | Calm3 | 1 |
| GO:009960 | BP | GO:009960 | regulation of neurotransmitter receptor activity | 1/19 | 72/28564 | 0.046835758 | 0.046835758 | 0.065355993 | Nptxr | 1 |
| GO:004866 | BP | GO:004866 | negative regulation of smooth muscle cell proliferation | 1/19 | 74/28564 | 0.048106599 | 0.048106599 | 0.065830083 | Apoe | 1 |
| GO:006104 | BP | GO:006104 | negative regulation of wound healing | 1/19 | 76/28564 | 0.049375835 | 0.049375835 | 0.066074918 | Apoe | 1 |
| GO:009055 | BP | GO:009055 | regulation of membrane permeability | 1/19 | 76/28564 | 0.049375835 | 0.049375835 | 0.066074918 | Camk2a | 1 |
| GO:003283 | CC | GO:003283 | dendrite cytoplasm | 2/19 | 33/28585 | 0.000218304 | 0.000218304 | 0.008732155 | Kif5a/Camk2a | 2 |
| GO:000807 | CC | GO:000807 | voltage-gated potassium channel complex | 2/19 | 77/28585 | 0.001188866 | 0.001188866 | 0.015851546 | Kcnj3/Calm3 | 2 |
| GO:003470 | CC | GO:003470 | potassium channel complex | 2/19 | 91/28585 | 0.001654649 | 0.001654649 | 0.016546489 | Kcnj3/Calm3 | 2 |
| GO:000587 | CC | GO:000587 | microtubule | 3/19 | 441/2858 | 0.002940651 | 0.002940651 | 0.023525209 | Apoe/Kif5a/Calm3 | 3 |
| GO:004238 | CC | GO:004238 | sarcolemma | 2/19 | 166/2858 | 0.005371992 | 0.005371992 | 0.030242456 | Kcnj3/Ctsb | 2 |
| GO:004262 | CC | GO:004262 | chylomicron | 1/19 | 12/28585 | 0.007948641 | 0.007948641 | 0.030242456 | Apoe | 1 |
| GO:003042 | CC | GO:003042 | growth cone | 2/19 | 213/2858 | 0.008693613 | 0.008693613 | 0.030242456 | Kif5a/Calm3 | 2 |
| GO:003042 | CC | GO:003042 | site of polarized growth | 2/19 | 221/2858 | 0.00933102 | 0.00933102 | 0.030242456 | Kif5a/Calm3 | 2 |
| GO:003525 | CC | GO:003525 | ciliary rootlet | 1/19 | 16/28585 | 0.01058486 | 0.01058486 | 0.030242456 | Kif5a | 1 |
| GO:190409 | CC | GO:190409 | peptidase inhibitor complex | 1/19 | 24/28585 | 0.015837401 | 0.015837401 | 0.036907508 | Ctsb | 1 |
| GO:004308 | CC | GO:004308 | synaptic cleft | 1/19 | 25/28585 | 0.016492108 | 0.016492108 | 0.036907508 | Apoe | 1 |
| GO:009952 | CC | GO:009952 | postsynaptic cytosol | 1/19 | 28/28585 | 0.018453754 | 0.018453754 | 0.036907508 | Camk2a | 1 |
| GO:009744 | CC | GO:009744 | apical dendrite | 1/19 | 30/28585 | 0.019759459 | 0.019759459 | 0.037608684 | Kif5a | 1 |
| GO:009868 | CC | GO:009868 | parallel fiber to Purkinje cell synapse | 1/19 | 37/28585 | 0.024316478 | 0.024316478 | 0.037608684 | Kcnj3 | 1 |
| GO:004247 | CC | GO:004247 | melanosome | 1/19 | 39/28585 | 0.025614792 | 0.025614792 | 0.037608684 | Apoe | 1 |
| GO:004877 | CC | GO:004877 | pigment granule | 1/19 | 39/28585 | 0.025614792 | 0.025614792 | 0.037608684 | Apoe | 1 |
| GO:199077 | CC | GO:199077 | lipoprotein particle | 1/19 | 43/28585 | 0.028206513 | 0.028206513 | 0.037608684 | Apoe | 1 |
| GO:199035 | CC | GO:199035 | transporter complex | 2/19 | 415/2858 | 0.030549936 | 0.030549936 | 0.037743355 | Kcnj3/Calm3 | 2 |
| GO:000587 | CC | GO:000587 | kinesin complex | 1/19 | 49/28585 | 0.032081852 | 0.032081852 | 0.037743355 | Kif5a | 1 |
| GO:000563 | CC | GO:000563 | nuclear envelope | 2/19 | 440/2858 | 0.034010996 | 0.034010996 | 0.038104815 | Apoe/Myorg | 2 |
| GO:001406 | CC | GO:001406 | postsynaptic density | 2/19 | 442/2858 | 0.034294334 | 0.034294334 | 0.038104815 | Camk2n1/Camk2a | 2 |
| GO:003227 | CC | GO:003227 | asymmetric synapse | 2/19 | 462/2858 | 0.037179106 | 0.037179106 | 0.040193628 | Camk2n1/Camk2a | 2 |
| GO:000577 | CC | GO:000577 | multivesicular body | 1/19 | 59/28585 | 0.038508234 | 0.038508234 | 0.040534983 | Apoe | 1 |
| GO:004339 | MF | GO:004339 | proteoglycan binding | 2/18 | 43/28171 | 0.00034283 | 0.00034283 | 0.019848062 | Apoe/Ctsb | 2 |
| GO:003525 | MF | GO:003525 | glutamate receptor binding | 2/18 | 74/28171 | 0.001013498 | 0.001013498 | 0.029338096 | Calm3/Camk2a | 2 |
| GO:001988 | MF | GO:001988 | protein kinase regulator activity | 2/18 | 173/2817 | 0.005377752 | 0.005377752 | 0.033910587 | Camk2n1/Calm3 | 2 |
| GO:001085 | MF | GO:001085 | cyclase activator activity | 1/18 | 10/28171 | 0.006372224 | 0.006372224 | 0.033910587 | Calm3 | 1 |
| GO:001674 | MF | GO:001674 | hydroxymethyl-, formyl- and related transferase activity | 1/18 | 10/28171 | 0.006372224 | 0.006372224 | 0.033910587 | Aldh1l1 | 1 |
| GO:003143 | MF | GO:003143 | titin binding | 1/18 | 12/28171 | 0.007642058 | 0.007642058 | 0.033910587 | Calm3 | 1 |
| GO:000216 | MF | GO:000216 | aminoacyl-tRNA editing activity | 1/18 | 13/28171 | 0.0082764 | 0.0082764 | 0.033910587 | Lars2 | 1 |
| GO:000817 | MF | GO:000817 | adenylate cyclase binding | 1/18 | 16/28171 | 0.01017713 | 0.01017713 | 0.033910587 | Calm3 | 1 |
| GO:001664 | MF | GO:001664 | oxidoreductase activity, acting on the CH-NH group of donors, NAD or | 1/18 | 16/28171 | 0.01017713 | 0.01017713 | 0.033910587 | Aldh1l1 | 1 |
| GO:007254 | MF | GO:007254 | protein phosphatase activator activity | 1/18 | 16/28171 | 0.01017713 | 0.01017713 | 0.033910587 | Calm3 | 1 |
| GO:000402 | MF | GO:000402 | aldehyde dehydrogenase (NAD+) activity | 1/18 | 17/28171 | 0.010809942 | 0.010809942 | 0.033910587 | Aldh1l1 | 1 |
| GO:009950 | MF | GO:009950 | voltage-gated monoatomic ion channel activity involved in regulation o | 1/18 | 17/28171 | 0.010809942 | 0.010809942 | 0.033910587 | Kcnj3 | 1 |
| GO:012002 | MF | GO:012002 | cholesterol transfer activity | 1/18 | 21/28171 | 0.01333737 | 0.01333737 | 0.033910587 | Apoe | 1 |
| GO:003279 | MF | GO:003279 | GTPase activating protein binding | 1/18 | 22/28171 | 0.013968273 | 0.013968273 | 0.033910587 | Camk2a | 1 |
| GO:004815 | MF | GO:004815 | tau protein binding | 1/18 | 22/28171 | 0.013968273 | 0.013968273 | 0.033910587 | Apoe | 1 |
| GO:005099 | MF | GO:005099 | nitric-oxide synthase binding | 1/18 | 22/28171 | 0.013968273 | 0.013968273 | 0.033910587 | Calm3 | 1 |
| GO:000524 | MF | GO:000524 | inward rectifier potassium channel activity | 1/18 | 23/28171 | 0.014598795 | 0.014598795 | 0.033910587 | Kcnj3 | 1 |
| GO:000468 | MF | GO:000468 | calmodulin-dependent protein kinase activity | 1/18 | 24/28171 | 0.015228936 | 0.015228936 | 0.033910587 | Camk2a | 1 |
| GO:000993 | MF | GO:000993 | calcium-dependent protein serine/threonine kinase activity | 1/18 | 24/28171 | 0.015228936 | 0.015228936 | 0.033910587 | Camk2a | 1 |
| GO:000857 | MF | GO:000857 | plus-end-directed microtubule motor activity | 1/18 | 26/28171 | 0.016488078 | 0.016488078 | 0.03409189 | Kif5a | 1 |
| GO:005075 | MF | GO:005075 | low-density lipoprotein particle receptor binding | 1/18 | 29/28171 | 0.018373939 | 0.018373939 | 0.034378257 | Apoe | 1 |
| GO:007181 | MF | GO:007181 | lipoprotein particle binding | 1/18 | 30/28171 | 0.0190018 | 0.0190018 | 0.034378257 | Apoe | 1 |
| GO:007181 | MF | GO:007181 | protein-lipid complex binding | 1/18 | 30/28171 | 0.0190018 | 0.0190018 | 0.034378257 | Apoe | 1 |
| GO:004227 | MF | GO:004227 | peptide binding | 2/18 | 361/2817 | 0.021882237 | 0.021882237 | 0.035629976 | Apoe/Ctsb | 2 |
| GO:014082 | MF | GO:014082 | metal cation:monoatomic cation antiporter activity | 1/18 | 36/28171 | 0.02276101 | 0.02276101 | 0.035629976 | Slc41a3 | 1 |
| GO:000153 | MF | GO:000153 | lipopolysaccharide binding | 1/18 | 37/28171 | 0.023386221 | 0.023386221 | 0.035629976 | Tril | 1 |
| GO:009771 | MF | GO:009771 | disordered domain specific binding | 1/18 | 37/28171 | 0.023386221 | 0.023386221 | 0.035629976 | Calm3 | 1 |
| GO:000481 | MF | GO:000481 | aminoacyl-tRNA ligase activity | 1/18 | 41/28171 | 0.025883287 | 0.025883287 | 0.036537935 | Lars2 | 1 |
| GO:001687 | MF | GO:001687 | ligase activity, forming carbon-oxygen bonds | 1/18 | 41/28171 | 0.025883287 | 0.025883287 | 0.036537935 | Lars2 | 1 |
| GO:004354 | MF | GO:004354 | phosphatidylinositol 3-kinase binding | 1/18 | 42/28171 | 0.026506611 | 0.026506611 | 0.036537935 | Calm3 | 1 |
| GO:000524 | MF | GO:000524 | calcium channel regulator activity | 1/18 | 45/28171 | 0.028374324 | 0.028374324 | 0.038092484 | Hpcal4 | 1 |
| GO:004687 | MF | GO:004687 | metal ion transmembrane transporter activity | 2/18 | 430/2817 | 0.03026621 | 0.03026621 | 0.038092484 | Kcnj3/Slc41a3 | 2 |
| GO:001989 | MF | GO:001989 | kinesin binding | 1/18 | 54/28171 | 0.033957176 | 0.033957176 | 0.04095712 | Kif5a | 1 |
| GO:004353 | MF | GO:004353 | protein serine/threonine kinase activator activity | 1/18 | 63/28171 | 0.039509722 | 0.039509722 | 0.045748099 | Calm3 | 1 |
| GO:000551 | MF | GO:000551 | collagen binding | 1/18 | 71/28171 | 0.044420002 | 0.044420002 | 0.048522345 | Ctsb | 1 |
| GO:000377 | MF | GO:000377 | microtubule motor activity | 1/18 | 74/28171 | 0.046255237 | 0.046255237 | 0.049591384 | Kif5a | 1 |
| GO:004830 | MF | GO:004830 | calcium-dependent protein binding | 1/18 | 76/28171 | 0.047476877 | 0.047476877 | 0.04997566 | Calm3 | 1 |

Cre+_vs_Cre-_enrichGO

| | ONTOLOGY | ID | Description | GeneRati | BgRatio | pvalue | p.adjust | qvalue | geneID | Count |
|---|---|---|---|---|---|---|---|---|---|---|
| GO:007180 | BP | GO:007180 | potassium ion transmembrane transport | 7/61 | 207/2856 | ################ | ################ | 0.000209008 | Slc24a4/Kcnd2/Dpp10/Cacna1d/Hcn1/Rgs7/Kcnq5 | 7 |
| GO:004239 | BP | GO:004239 | regulation of membrane potential | 9/61 | 488/2856 | ################ | ################ | 0.000209008 | Cacnb4/Slc24a4/Kcnd2/Gabrb2/Ryr2/Cacna1d/Gabrg3/Hcn1/R | 9 |
| GO:000681 | BP | GO:000681 | potassium ion transport | 7/61 | 245/2856 | ################ | ################ | 0.000209008 | Slc24a4/Kcnd2/Dpp10/Cacna1d/Hcn1/Rgs7/Kcnq5 | 7 |
| GO:003476 | BP | GO:003476 | regulation of monoatomic ion transmembrane transport | 9/61 | 492/2856 | ################ | ################ | 0.000209008 | Npsr1/Cacnb4/Kcnd2/Dpp10/Ryr2/Cacna1d/Hcn1/Rgs7/Kcnq5 | 9 |
| GO:001972 | BP | GO:001972 | calcium-mediated signaling | 6/61 | 179/2856 | ################ | ################ | 0.000249417 | Slc24a4/Ryr2/Cacna1d/Grm1/Nrg1/Rit2 | 6 |
| GO:008601 | BP | GO:008601 | SA node cell action potential | 3/61 | 13/28564 | ################ | ################ | 0.000249417 | Ryr2/Cacna1d/Hcn1 | 3 |
| GO:008601 | BP | GO:008601 | SA node cell to atrial cardiac muscle cell signaling | 3/61 | 13/28564 | ################ | ################ | 0.000249417 | Ryr2/Cacna1d/Hcn1 | 3 |
| GO:008607 | BP | GO:008607 | SA node cell to atrial cardiac muscle cell communication | 3/61 | 13/28564 | ################ | ################ | 0.000249417 | Ryr2/Cacna1d/Hcn1 | 3 |
| GO:004327 | BP | GO:004327 | positive regulation of monoatomic ion transport | 6/61 | 274/2856 | ################ | ################ | 0.001508751 | Npsr1/Cacnb4/Ryr2/Cacna1d/Hcn1/Rgs7 | 6 |
| GO:000721 | BP | GO:000721 | gamma-aminobutyric acid signaling pathway | 3/61 | 30/28564 | ################ | ################ | 0.001725115 | Cacnb4/Gabrb2/Gabrg3 | 3 |
| GO:000687 | BP | GO:000687 | intracellular calcium ion homeostasis | 6/61 | 316/2856 | ################ | ################ | 0.002634455 | Npsr1/Cacnb4/Slc24a4/Inpp4b/Ryr2/Tmem178 | 6 |
| GO:005507 | BP | GO:005507 | calcium ion homeostasis | 6/61 | 347/2856 | ################ | ################ | 0.003987898 | Npsr1/Cacnb4/Slc24a4/Inpp4b/Ryr2/Tmem178 | 6 |
| GO:001097 | BP | GO:001097 | positive regulation of neuron projection development | 5/61 | 237/2856 | 0.000153599 | 0.000153599 | 0.00572799 | Pacsin1/Serpini1/Tiam1/Nrg1/Rit2 | 5 |
| GO:003524 | BP | GO:003524 | synaptic transmission, glutamatergic | 4/61 | 127/2856 | 0.000159851 | 0.000159851 | 0.00572799 | Cacnb4/Syt1/Grm1/Hcn1 | 4 |
| GO:190247 | BP | GO:190247 | chloride transmembrane transport | 3/61 | 56/28564 | 0.000236973 | 0.000236973 | 0.008060515 | Gabrb2/Ano3/Gabrg3 | 3 |
| GO:000551 | BP | GO:000551 | detection of calcium ion | 2/61 | 11/28564 | 0.00024369 | 0.00024369 | 0.008060515 | Syt1/Ryr2 | 2 |
| GO:000762 | BP | GO:000762 | locomotory behavior | 5/61 | 270/2856 | 0.000280806 | 0.000280806 | 0.008622819 | Cacnb4/Kcnd2/Grm1/Nrg1/Chl1 | 5 |
| GO:000627 | BP | GO:000627 | RNA-templated DNA biosynthetic process | 3/61 | 60/28564 | 0.000290769 | 0.000290769 | 0.008622819 | Rad50/Pkib/Ptges3 | 3 |
| GO:000700 | BP | GO:000700 | telomere maintenance via telomerase | 3/61 | 60/28564 | 0.000290769 | 0.000290769 | 0.008622819 | Rad50/Pkib/Ptges3 | 3 |
| GO:003134 | BP | GO:003134 | positive regulation of cell projection organization | 6/61 | 459/2856 | 0.000438374 | 0.000438374 | 0.011702902 | Pacsin1/Serpini1/Tiam1/Nrg1/Rit2/Gpm6a | 6 |
| GO:190399 | BP | GO:190399 | regulation of eating behavior | 2/61 | 15/28564 | 0.000462673 | 0.000462673 | 0.011702902 | Npsr1/Slc24a4 | 2 |
| GO:009755 | BP | GO:009755 | calcium ion transmembrane import into cytosol | 4/61 | 193/2856 | 0.000780437 | 0.000780437 | 0.017192823 | Npsr1/Slc24a4/Ryr2/Cacna1d | 4 |
| GO:007058 | BP | GO:007058 | calcium ion transmembrane transport | 5/61 | 339/2856 | 0.000789187 | 0.000789187 | 0.017192823 | Npsr1/Cacnb4/Slc24a4/Ryr2/Cacna1d | 5 |
| GO:005196 | BP | GO:005196 | regulation of synaptic transmission, glutamatergic | 3/61 | 88/28564 | 0.000893679 | 0.000893679 | 0.018049337 | Syt1/Grm1/Hcn1 | 3 |
| GO:001922 | BP | GO:001922 | transmission of nerve impulse | 3/61 | 101/2856 | 0.001330736 | 0.001330736 | 0.02487897 | Cacnb4/Kcnd2/Hcn1 | 3 |
| GO:200005 | BP | GO:200005 | regulation of non-canonical Wnt signaling pathway | 2/61 | 26/28564 | 0.001410594 | 0.001410594 | 0.025273148 | Tiam1/Rspo3 | 2 |
| GO:004692 | BP | GO:004692 | regulation of neurotransmitter secretion | 3/61 | 104/2856 | 0.001447555 | 0.001447555 | 0.025406065 | Cacnb4/Mctp1/Syt1 | 3 |
| GO:003195 | BP | GO:003195 | positive regulation of protein autophosphorylation | 2/61 | 27/28564 | 0.001521351 | 0.001521351 | 0.026167242 | Rad50/Nrg1 | 2 |
| GO:000682 | BP | GO:000682 | chloride transport | 3/61 | 108/2856 | 0.00161306 | 0.00161306 | 0.027059079 | Gabrb2/Ano3/Gabrg3 | 3 |
| GO:009914 | BP | GO:009914 | regulation of postsynaptic neurotransmitter receptor internaliz | 2/61 | 28/28564 | 0.00163613 | 0.00163613 | 0.027059079 | Pacsin1/Nrg1 | 2 |
| GO:006004 | BP | GO:006004 | heart contraction | 4/61 | 242/2856 | 0.001796493 | 0.001796493 | 0.028610816 | Ryr2/Cacna1d/Sgcz/Hcn1 | 4 |
| GO:001407 | BP | GO:001407 | response to purine-containing compound | 3/61 | 114/2856 | 0.001882779 | 0.001882779 | 0.0289141 | Pde3a/Ryr2/Hcn1 | 3 |
| GO:003010 | BP | GO:003010 | regulation of endocytosis | 4/61 | 248/2856 | 0.001963864 | 0.001963864 | 0.02933705 | Mctp1/Pacsin1/Nrg1/Rit2 | 4 |
| GO:004326 | BP | GO:004326 | regulation of potassium ion transport | 3/61 | 116/2856 | 0.001978545 | 0.001978545 | 0.02933705 | Dpp10/Cacna1d/Rgs7 | 3 |
| GO:000165 | BP | GO:000165 | eye development | 5/61 | 426/2856 | 0.002160549 | 0.002160549 | 0.030893679 | Gpd2/Olfm3/Rhoj/Hcn1/Gpm6a | 5 |
| GO:005197 | BP | GO:005197 | positive regulation of telomerase activity | 2/61 | 33/28564 | 0.00226976 | 0.00226976 | 0.030984023 | Pkib/Ptges3 | 2 |
| GO:005158 | BP | GO:005158 | regulation of neurotransmitter transport | 3/61 | 124/2856 | 0.002391792 | 0.002391792 | 0.031645251 | Cacnb4/Mctp1/Syt1 | 3 |
| GO:009900 | BP | GO:009900 | vesicle-mediated transport in synapse | 4/61 | 271/2856 | 0.002705763 | 0.002705763 | 0.035256911 | Cacnb4/Syt1/Pacsin1/Nrg1 | 4 |
| GO:000681 | BP | GO:000681 | calcium ion transport | 5/61 | 452/2856 | 0.002789365 | 0.002789365 | 0.035581942 | Npsr1/Cacnb4/Slc24a4/Ryr2/Cacna1d | 5 |
| GO:000958 | BP | GO:000958 | detection of abiotic stimulus | 3/61 | 140/2856 | 0.00336955 | 0.00336955 | 0.038019894 | Cacnb4/Slc24a4/Ano3 | 3 |
| GO:003241 | BP | GO:003241 | positive regulation of transporter activity | 3/61 | 140/2856 | 0.00336955 | 0.00336955 | 0.038019894 | Cacnb4/Ryr2/Hcn1 | 3 |
| GO:000958 | BP | GO:000958 | detection of external stimulus | 3/61 | 141/2856 | 0.003437613 | 0.003437613 | 0.038019894 | Cacnb4/Slc24a4/Ano3 | 3 |
| GO:004567 | BP | GO:004567 | negative regulation of osteoclast differentiation | 2/61 | 43/28564 | 0.003828954 | 0.003828954 | 0.039201198 | Inpp4b/Tmem178 | 2 |
| GO:008600 | BP | GO:008600 | cardiac muscle cell action potential involved in contraction | 2/61 | 43/28564 | 0.003828954 | 0.003828954 | 0.039201198 | Ryr2/Cacna1d | 2 |
| GO:003220 | BP | GO:003220 | telomere organization | 3/61 | 150/2856 | 0.004088291 | 0.004088291 | 0.040092452 | Rad50/Pkib/Ptges3 | 3 |
| GO:008600 | BP | GO:008600 | membrane repolarization | 2/61 | 45/28564 | 0.004186373 | 0.004186373 | 0.040092452 | Slc24a4/Cacna1d | 2 |
| GO:000320 | BP | GO:000320 | cardiac chamber morphogenesis | 3/61 | 152/2856 | 0.004242341 | 0.004242341 | 0.040092452 | Dnah11/Ryr2/Nrg1 | 3 |
| GO:001923 | BP | GO:001923 | sensory perception of pain | 3/61 | 152/2856 | 0.004242341 | 0.004242341 | 0.040092452 | Kcnd2/Ptges3/Grm1 | 3 |
| GO:004246 | BP | GO:004246 | eye photoreceptor cell development | 2/61 | 46/28564 | 0.004370674 | 0.004370674 | 0.040416987 | Olfm3/Hcn1 | 2 |
| GO:003240 | BP | GO:003240 | regulation of transporter activity | 4/61 | 317/2856 | 0.004726322 | 0.004726322 | 0.042785654 | Cacnb4/Ryr2/Cacna1d/Hcn1 | 4 |
| GO:000325 | BP | GO:000325 | regulation of membrane depolarization | 2/61 | 51/28564 | 0.005347443 | 0.005347443 | 0.04598801 | Cacna1d/Hcn1 | 2 |
| GO:009964 | BP | GO:009964 | signal release from synapse | 3/61 | 169/2856 | 0.005694948 | 0.005694948 | 0.048016231 | Cacnb4/Mctp1/Syt1 | 3 |
| GO:004275 | BP | GO:004275 | eating behavior | 2/61 | 53/28564 | 0.005763642 | 0.005763642 | 0.048123612 | Npsr1/Slc24a4 | 2 |
| GO:000958 | BP | GO:000958 | detection of light stimulus | 2/61 | 54/28564 | 0.00597714 | 0.00597714 | 0.04942635 | Cacnb4/Slc24a4 | 2 |
| GO:005501 | BP | GO:005501 | ventricular cardiac muscle tissue morphogenesis | 2/61 | 56/28564 | 0.006414855 | 0.006414855 | 0.052540717 | Ryr2/Nrg1 | 2 |
| GO:005123 | BP | GO:005123 | maintenance of location | 4/61 | 364/2856 | 0.007649177 | 0.007649177 | 0.059263893 | Npsr1/Ryr2/Nrg1/Rit2 | 4 |
| GO:000322 | BP | GO:000322 | ventricular cardiac muscle tissue development | 2/61 | 66/28564 | 0.008813744 | 0.008813744 | 0.065343272 | Ryr2/Nrg1 | 2 |
| GO:004828 | BP | GO:004828 | lung alveolus development | 2/61 | 67/28564 | 0.009072543 | 0.009072543 | 0.06556628 | Meg3/Ptges3 | 2 |
| GO:000176 | BP | GO:000176 | neuron migration | 3/61 | 204/2856 | 0.009532734 | 0.009532734 | 0.067753319 | Nrg1/Chl1/Gpm6a | 3 |
| GO:009873 | BP | GO:009873 | import across plasma membrane | 3/61 | 208/2856 | 0.01004697 | 0.01004697 | 0.069218836 | Slc24a4/Cacna1d/Grm1 | 3 |
| GO:000741 | BP | GO:000741 | synapse assembly | 3/61 | 216/2856 | 0.011123243 | 0.011123243 | 0.074734289 | Gabrb2/Nrg1/Gpm6a | 3 |
| GO:000683 | BP | GO:000683 | neurotransmitter transport | 3/61 | 225/2856 | 0.012410956 | 0.012410956 | 0.081030758 | Cacnb4/Mctp1/Syt1 | 3 |
| GO:004649 | BP | GO:004649 | nicotinamide nucleotide metabolic process | 2/61 | 79/28564 | 0.012437279 | 0.012437279 | 0.081030758 | Aldh1l1/Gpd2 | 2 |
| GO:000209 | BP | GO:000209 | regulation of receptor internalization | 2/61 | 83/28564 | 0.013662484 | 0.013662484 | 0.084530473 | Pacsin1/Nrg1 | 2 |
| GO:000150 | BP | GO:000150 | regulation of neurotransmitter levels | 3/61 | 242/2856 | 0.015068334 | 0.015068334 | 0.089991441 | Cacnb4/Mctp1/Syt1 | 3 |
| GO:006117 | BP | GO:006117 | regulation of insulin secretion involved in cellular response to | 2/61 | 90/28564 | 0.015926973 | 0.015926973 | 0.093816418 | Cacna1d/Tiam1 | 2 |
| GO:002167 | BP | GO:002167 | nerve development | 2/61 | 93/28564 | 0.016943374 | 0.016943374 | 0.096881821 | Slc24a4/Gabrb2 | 2 |
| GO:000741 | BP | GO:000741 | axon guidance | 3/61 | 253/2856 | 0.016946351 | 0.016946351 | 0.096881821 | Ablim1/Epha6/Chl1 | 3 |
| GO:005125 | BP | GO:005125 | protein complex oligomerization | 3/61 | 254/2856 | 0.017123299 | 0.017123299 | 0.096881821 | Kcnd2/Syt1/Hcn1 | 3 |
| GO:004668 | BP | GO:004668 | response to organophosphorus | 2/61 | 95/28564 | 0.017635994 | 0.017635994 | 0.097851322 | Pde3a/Hcn1 | 2 |
| GO:004354 | BP | GO:004354 | positive regulation of GTPase activity | 3/61 | 259/2856 | 0.01802363 | 0.01802363 | 0.098103303 | Rasgef1b/Tiam1/Rgs7 | 3 |
| GO:009865 | BP | GO:009865 | import into cell | 3/61 | 264/2856 | 0.018949994 | 0.018949994 | 0.10116564 | Slc24a4/Cacna1d/Grm1 | 3 |
| GO:003432 | BP | GO:003432 | cell junction assembly | 4/61 | 477/2856 | 0.018967769 | 0.018967769 | 0.10116564 | Gabrb2/Nrg1/Patj/Gpm6a | 4 |
| GO:007025 | BP | GO:007025 | actin-mediated cell contraction | 2/61 | 99/28564 | 0.019056783 | 0.019056783 | 0.10116564 | Ryr2/Cacna1d | 2 |
| GO:002182 | BP | GO:002182 | substrate-independent telencephalic tangential migration | 1/61 | 10/28564 | 0.021154791 | 0.021154791 | 0.102942062 | Nrg1 | 1 |
| GO:002184 | BP | GO:002184 | substrate-independent telencephalic tangential interneuron mi | 1/61 | 10/28564 | 0.021154791 | 0.021154791 | 0.102942062 | Nrg1 | 1 |
| GO:003360 | BP | GO:003360 | positive regulation of dopamine secretion | 1/61 | 10/28564 | 0.021154791 | 0.021154791 | 0.102942062 | Syt1 | 1 |
| GO:005177 | BP | GO:005177 | response to redox state | 1/61 | 10/28564 | 0.021154791 | 0.021154791 | 0.102942062 | Ryr2 | 1 |
| GO:007125 | BP | GO:007125 | cellular response to electrical stimulus | 1/61 | 10/28564 | 0.021154791 | 0.021154791 | 0.102942062 | Grm1 | 1 |
| GO:007187 | BP | GO:007187 | response to epinephrine | 1/61 | 10/28564 | 0.021154791 | 0.021154791 | 0.102942062 | Ryr2 | 1 |
| GO:007187 | BP | GO:007187 | cellular response to epinephrine stimulus | 1/61 | 10/28564 | 0.021154791 | 0.021154791 | 0.102942062 | Ryr2 | 1 |
| GO:190527 | BP | GO:190527 | regulation of modification of postsynaptic actin cytoskeleton | 1/61 | 10/28564 | 0.021154791 | 0.021154791 | 0.102942062 | Tiam1 | 1 |
| GO:200072 | BP | GO:200072 | positive regulation of cardiac muscle cell differentiation | 1/61 | 10/28564 | 0.021154791 | 0.021154791 | 0.102942062 | Nrg1 | 1 |
| GO:002260 | BP | GO:002260 | regulation of cell morphogenesis | 3/61 | 278/2856 | 0.021682682 | 0.021682682 | 0.102942062 | Syt1/Rhoj/Tiam1 | 3 |
| GO:200125 | BP | GO:200125 | positive regulation of chromosome organization | 2/61 | 109/2856 | 0.022811014 | 0.022811014 | 0.102942062 | Rad50/Pkib | 2 |
| GO:003813 | BP | GO:003813 | ERBB4 signaling pathway | 1/61 | 11/28564 | 0.023245901 | 0.023245901 | 0.102942062 | Nrg1 | 1 |
| GO:006028 | BP | GO:006028 | epithelial cilium movement involved in determination of left/right | 1/61 | 11/28564 | 0.023245901 | 0.023245901 | 0.102942062 | Dnah11 | 1 |
| GO:007141 | BP | GO:007141 | cellular response to purine-containing compound | 1/61 | 11/28564 | 0.023245901 | 0.023245901 | 0.102942062 | Ryr2 | 1 |
| GO:007142 | BP | GO:007142 | cellular response to histamine | 1/61 | 11/28564 | 0.023245901 | 0.023245901 | 0.102942062 | Gabrb2 | 1 |
| GO:190014 | BP | GO:190014 | regulation of Schwann cell migration | 1/61 | 11/28564 | 0.023245901 | 0.023245901 | 0.102942062 | Tiam1 | 1 |
| GO:200122 | BP | GO:200122 | negative regulation of neuron migration | 1/61 | 11/28564 | 0.023245901 | 0.023245901 | 0.102942062 | Nrg1 | 1 |
| GO:000330 | BP | GO:000330 | cardiac muscle hypertrophy | 2/61 | 111/2856 | 0.023595669 | 0.023595669 | 0.102942062 | Ryr2/Tiam1 | 2 |
| GO:000834 | BP | GO:000834 | adult locomotory behavior | 2/61 | 112/2856 | 0.023992136 | 0.023992136 | 0.102942062 | Cacnb4/Chl1 | 2 |
| GO:000607 | BP | GO:000607 | glycerol-3-phosphate metabolic process | 1/61 | 12/28564 | 0.025332617 | 0.025332617 | 0.102942062 | Gpd2 | 1 |
| GO:000682 | BP | GO:000682 | manganese ion transport | 1/61 | 12/28564 | 0.025332617 | 0.025332617 | 0.102942062 | Ryr2 | 1 |
| GO:000964 | BP | GO:000964 | response to light intensity | 1/61 | 12/28564 | 0.025332617 | 0.025332617 | 0.102942062 | Slc24a4 | 1 |
| GO:001062 | BP | GO:001062 | regulation of Schwann cell proliferation | 1/61 | 12/28564 | 0.025332617 | 0.025332617 | 0.102942062 | Nrg1 | 1 |
| GO:006001 | BP | GO:006001 | righting reflex | 1/61 | 12/28564 | 0.025332617 | 0.025332617 | 0.102942062 | Npsr1 | 1 |
| GO:006037 | BP | GO:006037 | cardiac muscle cell myoblast differentiation | 1/61 | 12/28564 | 0.025332617 | 0.025332617 | 0.102942062 | Nrg1 | 1 |
| GO:009891 | BP | GO:009891 | regulation of ventricular cardiac muscle cell action potential | 1/61 | 12/28564 | 0.025332617 | 0.025332617 | 0.102942062 | Ryr2 | 1 |
| GO:001470 | BP | GO:001470 | striated muscle tissue development | 3/61 | 296/2856 | 0.025497476 | 0.025497476 | 0.102942062 | Ryr2/Sgcz/Nrg1 | 3 |
| GO:004853 | BP | GO:004853 | mucosa-associated lymphoid tissue development | 1/61 | 13/28564 | 0.027414948 | 0.027414948 | 0.102942062 | Cacnb4 | 1 |
| GO:004854 | BP | GO:004854 | Peyer's patch development | 1/61 | 13/28564 | 0.027414948 | 0.027414948 | 0.102942062 | Cacnb4 | 1 |
| GO:190163 | BP | GO:190163 | positive regulation of synaptic vesicle membrane organization | 1/61 | 13/28564 | 0.027414948 | 0.027414948 | 0.102942062 | Syt1 | 1 |
| GO:190433 | BP | GO:190433 | regulation of dopaminergic neuron differentiation | 1/61 | 13/28564 | 0.027414948 | 0.027414948 | 0.102942062 | Tiam1 | 1 |
| GO:200084 | BP | GO:200084 | regulation of glucocorticoid secretion | 1/61 | 13/28564 | 0.027414948 | 0.027414948 | 0.102942062 | Nrg1 | 1 |
| GO:005080 | BP | GO:005080 | regulation of synapse structure or activity | 3/61 | 313/2856 | 0.029411446 | 0.029411446 | 0.102942062 | Tiam1/Nrg1/Gpm6a | 3 |
| GO:001088 | BP | GO:001088 | regulation of cardiac muscle contraction by calcium ion signali | 1/61 | 14/28564 | 0.029492903 | 0.029492903 | 0.102942062 | Ryr2 | 1 |
| GO:004298 | BP | GO:004298 | amyloid precursor protein biosynthetic process | 1/61 | 14/28564 | 0.029492903 | 0.029492903 | 0.102942062 | Necab1 | 1 |
| GO:004298 | BP | GO:004298 | regulation of amyloid precursor protein biosynthetic process | 1/61 | 14/28564 | 0.029492903 | 0.029492903 | 0.102942062 | Necab1 | 1 |
| GO:004517 | BP | GO:004517 | apical protein localization | 1/61 | 14/28564 | 0.029492903 | 0.029492903 | 0.102942062 | Hcn1 | 1 |
| GO:006067 | BP | GO:006067 | branching involved in labyrinthine layer morphogenesis | 1/61 | 14/28564 | 0.029492903 | 0.029492903 | 0.102942062 | Rspo3 | 1 |
| GO:200048 | BP | GO:200048 | regulation of hepatic stellate cell activation | 1/61 | 14/28564 | 0.029492903 | 0.029492903 | 0.102942062 | Rian | 1 |
| GO:009865 | BP | GO:009865 | inorganic cation import across plasma membrane | 2/61 | 126/2856 | 0.029824706 | 0.029824706 | 0.102942062 | Slc24a4/Cacna1d | 2 |
| GO:009958 | BP | GO:009958 | inorganic ion import across plasma membrane | 2/61 | 126/2856 | 0.029824706 | 0.029824706 | 0.102942062 | Slc24a4/Cacna1d | 2 |
| GO:000204 | BP | GO:000204 | sprouting angiogenesis | 2/61 | 128/2856 | 0.030699743 | 0.030699743 | 0.102942062 | Rhoj/Rspo3 | 2 |
| GO:003593 | BP | GO:003593 | glucocorticoid secretion | 1/61 | 15/28564 | 0.031566491 | 0.031566491 | 0.102942062 | Nrg1 | 1 |
| GO:004605 | BP | GO:004605 | cAMP metabolic process | 1/61 | 15/28564 | 0.031566491 | 0.031566491 | 0.102942062 | Cacnb4 | 1 |
| GO:009907 | BP | GO:009907 | regulation of postsynaptic membrane neurotransmitter recepto | 2/61 | 133/2856 | 0.032931524 | 0.032931524 | 0.102942062 | Pacsin1/Nrg1 | 2 |
| GO:007037 | BP | GO:007037 | regulation of ERK1 and ERK2 cascade | 3/61 | 329/2856 | 0.033370046 | 0.033370046 | 0.102942062 | Npsr1/Tiam1/Nrg1 | 3 |
| GO:003031 | BP | GO:003031 | osteoclast differentiation | 2/61 | 134/2856 | 0.033385356 | 0.033385356 | 0.102942062 | Inpp4b/Tmem178 | 2 |
| GO:001569 | BP | GO:001569 | magnesium ion transport | 1/61 | 16/28564 | 0.03363572 | 0.03363572 | 0.102942062 | Slc41a3 | 1 |
| GO:003573 | BP | GO:003573 | hepatic stellate cell activation | 1/61 | 16/28564 | 0.03363572 | 0.03363572 | 0.102942062 | Rian | 1 |
| GO:004292 | BP | GO:004292 | glucocorticoid receptor signaling pathway | 1/61 | 16/28564 | 0.03363572 | 0.03363572 | 0.102942062 | Ptges3 | 1 |
| GO:004364 | BP | GO:004364 | dicarboxylic acid catabolic process | 1/61 | 16/28564 | 0.03363572 | 0.03363572 | 0.102942062 | Aldh1l1 | 1 |
| GO:006070 | BP | GO:006070 | regulation of ribonuclease activity | 1/61 | 16/28564 | 0.03363572 | 0.03363572 | 0.102942062 | Pde3a | 1 |
| GO:009705 | BP | GO:009705 | type B pancreatic cell apoptotic process | 1/61 | 16/28564 | 0.03363572 | 0.03363572 | 0.102942062 | Ryr2 | 1 |
| GO:009952 | BP | GO:009952 | postsynapse to nucleus signaling pathway | 1/61 | 16/28564 | 0.03363572 | 0.03363572 | 0.102942062 | Nrg1 | 1 |
| GO:190257 | BP | GO:190257 | protein localization to nucleolus | 1/61 | 16/28564 | 0.03363572 | 0.03363572 | 0.102942062 | Cacnb4 | 1 |
| GO:000827 | BP | GO:000827 | regulation of G protein-coupled receptor signaling pathway | 2/61 | 137/2856 | 0.034761584 | 0.034761584 | 0.104786748 | Slc24a4/Rgs7 | 2 |
| GO:004688 | BP | GO:004688 | regulation of hormone secretion | 3/61 | 337/2856 | 0.035448683 | 0.035448683 | 0.104786748 | Cacna1d/Tiam1/Nrg1 | 3 |
| GO:000335 | BP | GO:000335 | regulation of cilium beat frequency | 1/61 | 17/28564 | 0.035700601 | 0.035700601 | 0.104786748 | Dnah11 | 1 |
| GO:000673 | BP | GO:000673 | NADH metabolic process | 1/61 | 17/28564 | 0.035700601 | 0.035700601 | 0.104786748 | Gpd2 | 1 |
| GO:008601 | BP | GO:008601 | membrane depolarization during cardiac muscle cell action pot | 1/61 | 17/28564 | 0.035700601 | 0.035700601 | 0.104786748 | Cacna1d | 1 |
| GO:009871 | BP | GO:009871 | L-glutamate import across plasma membrane | 1/61 | 17/28564 | 0.035700601 | 0.035700601 | 0.104786748 | Grm1 | 1 |
| GO:190207 | BP | GO:190207 | response to salt | 3/61 | 340/2856 | 0.036245156 | 0.036245156 | 0.106023245 | Syt1/Ryr2/Hcn1 | 3 |
| GO:002178 | BP | GO:002178 | glial cell fate commitment | 1/61 | 18/28564 | 0.037761142 | 0.037761142 | 0.106824284 | Nrg1 | 1 |
| GO:003279 | BP | GO:003279 | positive regulation of CREB transcription factor activity | 1/61 | 18/28564 | 0.037761142 | 0.037761142 | 0.106824284 | Cacna1d | 1 |
| GO:005113 | BP | GO:005113 | chaperone-mediated protein complex assembly | 1/61 | 18/28564 | 0.037761142 | 0.037761142 | 0.106824284 | Ptges3 | 1 |
| GO:005501 | BP | GO:005501 | ventricular cardiac muscle cell differentiation | 1/61 | 18/28564 | 0.037761142 | 0.037761142 | 0.106824284 | Nrg1 | 1 |
| GO:004308 | BP | GO:004308 | regulation of GTPase activity | 3/61 | 353/2856 | 0.039802988 | 0.039802988 | 0.10836368 | Rasgef1b/Tiam1/Rgs7 | 3 |
| GO:001405 | BP | GO:001405 | gamma-aminobutyric acid secretion | 1/61 | 19/28564 | 0.039817352 | 0.039817352 | 0.10836368 | Cacnb4 | 1 |
| GO:004595 | BP | GO:004595 | positive regulation of calcium ion-dependent exocytosis | 1/61 | 19/28564 | 0.039817352 | 0.039817352 | 0.10836368 | Syt1 | 1 |
| GO:007151 | BP | GO:007151 | genomic imprinting | 1/61 | 19/28564 | 0.039817352 | 0.039817352 | 0.10836368 | Meg3 | 1 |
| GO:003162 | BP | GO:003162 | receptor internalization | 2/61 | 151/2856 | 0.041467554 | 0.041467554 | 0.109360441 | Pacsin1/Nrg1 | 2 |
| GO:004001 | BP | GO:004001 | negative regulation of locomotion | 3/61 | 360/2856 | 0.04178996 | 0.04178996 | 0.109360441 | Mctp1/Nrg1/Mcc | 3 |
| GO:000072 | BP | GO:000072 | DNA double-strand break processing | 1/61 | 20/28564 | 0.04186924 | 0.04186924 | 0.109360441 | Rad50 | 1 |
| GO:004311 | BP | GO:004311 | negative regulation of vascular permeability | 1/61 | 20/28564 | 0.04186924 | 0.04186924 | 0.109360441 | Pde3a | 1 |
| GO:007088 | BP | GO:007088 | positive regulation of calcineurin-NFAT signaling cascade | 1/61 | 20/28564 | 0.04186924 | 0.04186924 | 0.109360441 | Nrg1 | 1 |
| GO:009732 | BP | GO:009732 | plasma membrane tubulation | 1/61 | 20/28564 | 0.04186924 | 0.04186924 | 0.109360441 | Pacsin1 | 1 |
| GO:009888 | BP | GO:009888 | modification of postsynaptic actin cytoskeleton | 1/61 | 20/28564 | 0.04186924 | 0.04186924 | 0.109360441 | Tiam1 | 1 |
| GO:010605 | BP | GO:010605 | positive regulation of calcineurin-mediated signaling | 1/61 | 20/28564 | 0.04186924 | 0.04186924 | 0.109360441 | Nrg1 | 1 |
| GO:005089 | BP | GO:005089 | cognition | 3/61 | 364/2856 | 0.042947601 | 0.042947601 | 0.111222319 | Dnah11/Nrg1/Chl1 | 3 |
| GO:003599 | BP | GO:003599 | response to muscle stretch | 1/61 | 21/28564 | 0.043916815 | 0.043916815 | 0.111222319 | Ryr2 | 1 |
| GO:009005 | BP | GO:009005 | positive regulation of cell migration involved in sprouting angio | 1/61 | 21/28564 | 0.043916815 | 0.043916815 | 0.111222319 | Rhoj | 1 |
| GO:007133 | BP | GO:007133 | cellular response to glucose stimulus | 2/61 | 156/2856 | 0.043971615 | 0.043971615 | 0.111222319 | Cacna1d/Tiam1 | 2 |
| GO:000183 | BP | GO:000183 | epithelial to mesenchymal transition | 2/61 | 158/2856 | 0.044988744 | 0.044988744 | 0.113129591 | Rian/Tiam1 | 2 |
| GO:007133 | BP | GO:007133 | cellular response to hexose stimulus | 2/61 | 158/2856 | 0.044988744 | 0.044988744 | 0.113129591 | Cacna1d/Tiam1 | 2 |
| GO:000674 | BP | GO:000674 | NADPH regeneration | 1/61 | 22/28564 | 0.045960086 | 0.045960086 | 0.11325408 | Aldh1l1 | 1 |
| GO:001712 | BP | GO:001712 | plasma membrane phospholipid scrambling | 1/61 | 22/28564 | 0.045960086 | 0.045960086 | 0.11325408 | Ano3 | 1 |
| GO:004547 | BP | GO:004547 | locomotor rhythm | 1/61 | 22/28564 | 0.045960086 | 0.045960086 | 0.11325408 | Kcnd2 | 1 |
| GO:003572 | BP | GO:003572 | sodium ion transmembrane transport | 2/61 | 166/2856 | 0.049143602 | 0.049143602 | 0.11788774 | Slc24a4/Hcn1 | 2 |
| GO:000686 | BP | GO:000686 | amino acid transport | 2/61 | 167/2856 | 0.049672487 | 0.049672487 | 0.11788774 | Cacnb4/Grm1 | 2 |
| GO:003470 | CC | GO:003470 | ion channel complex | 10/61 | 301/2858 | ################ | ################ | ################ | Cacnb4/Kcnd2/Dpp10/Olfm3/Gabrb2/Ryr2/Cacna1d/Gabrg3/Hc | 10 |
| GO:199035 | CC | GO:199035 | transporter complex | 10/61 | 415/2858 | ################ | ################ | ################ | Cacnb4/Kcnd2/Dpp10/Olfm3/Gabrb2/Ryr2/Cacna1d/Gabrg3/Hc | 10 |
| GO:004273 | CC | GO:004273 | presynaptic membrane | 8/61 | 238/2858 | ################ | ################ | ################ | Syt1/Gabrb2/Cacna1d/Grm1/Nrg1/Hcn1/Gpm6a/Kcnq5 | 8 |
| GO:004521 | CC | GO:004521 | postsynaptic membrane | 9/61 | 348/2858 | ################ | ################ | ################ | Kcnd2/Gabrb2/Pacsin1/Grm1/Gabrg3/Tiam1/Nrg1/Hcn1/Rgs7 | 9 |
| GO:004878 | CC | GO:004878 | presynaptic active zone | 5/61 | 122/2858 | ################ | ################ | ################ | Gabrb2/Cacna1d/Nrg1/Hcn1/Gpm6a | 5 |
| GO:000807 | CC | GO:000807 | voltage-gated potassium channel complex | 4/61 | 77/28585 | ################ | ################ | 0.000219128 | Kcnd2/Dpp10/Hcn1/Kcnq5 | 4 |
| GO:003125 | CC | GO:003125 | leading edge membrane | 5/61 | 159/2858 | ################ | ################ | 0.000219128 | Pacsin1/Cacna1d/Gabrg3/Tiam1/Hcn1 | 5 |
| GO:009957 | CC | GO:009957 | postsynaptic specialization | 7/61 | 486/2858 | ################ | ################ | 0.000622575 | Kcnd2/Gabrb2/Pacsin1/Ablim1/Grm1/Tiam1/Nrg1 | 7 |
| GO:003259 | CC | GO:003259 | dendrite membrane | 3/61 | 50/28585 | 0.000168727 | 0.000168727 | 0.001172207 | Cacna1d/Gabrg3/Hcn1 | 3 |
| GO:009898 | CC | GO:009898 | GABA-ergic synapse | 4/61 | 134/2858 | 0.000195869 | 0.000195869 | 0.001275728 | Kcnd2/Gabrb2/Gabrg3/Nrg1 | 4 |
| GO:003125 | CC | GO:003125 | cell leading edge | 6/61 | 412/2858 | 0.000245832 | 0.000245832 | 0.001506956 | Pacsin1/Ablim1/Cacna1d/Gabrg3/Tiam1/Hcn1 | 6 |
| GO:004238 | CC | GO:004238 | sarcolemma | 4/61 | 166/2858 | 0.000442309 | 0.000442309 | 0.00224277 | Kcnd2/Ryr2/Cacna1d/Sgcz | 4 |
| GO:003227 | CC | GO:003227 | asymmetric synapse | 6/61 | 462/2858 | 0.000451952 | 0.000451952 | 0.00224277 | Kcnd2/Pacsin1/Ablim1/Grm1/Tiam1/Nrg1 | 6 |
| GO:190271 | CC | GO:190271 | GABA-A receptor complex | 2/61 | 19/28585 | 0.000748275 | 0.000748275 | 0.003249088 | Gabrb2/Gabrg3 | 2 |
| GO:004430 | CC | GO:004430 | neuron projection terminus | 4/61 | 224/2858 | 0.001350488 | 0.001350488 | 0.005412885 | Syt1/Pacsin1/Hcn1/Kcnq5 | 4 |
| GO:009747 | CC | GO:009747 | ribbon synapse | 2/61 | 27/28585 | 0.001519155 | 0.001519155 | 0.005863406 | Pacsin1/Cacna1d | 2 |
| GO:015003 | CC | GO:015003 | distal axon | 5/61 | 397/2858 | 0.001583961 | 0.001583961 | 0.005895194 | Pacsin1/Tiam1/Hcn1/Gpm6a/Kcnq5 | 5 |
| GO:004367 | CC | GO:004367 | axon terminus | 3/61 | 202/2858 | 0.009263142 | 0.009263142 | 0.030166154 | Pacsin1/Hcn1/Kcnq5 | 3 |
| GO:004319 | CC | GO:004319 | dendritic spine | 3/61 | 206/2858 | 0.0097685 | 0.0097685 | 0.030206716 | Kcnd2/Grm1/Tiam1 | 3 |
| GO:004430 | CC | GO:004430 | neuron spine | 3/61 | 212/2858 | 0.010556246 | 0.010556246 | 0.031430627 | Kcnd2/Grm1/Tiam1 | 3 |
| GO:003123 | CC | GO:003123 | extrinsic component of cytoplasmic side of plasma membrane | 2/61 | 95/28585 | 0.017611727 | 0.017611727 | 0.048298088 | Ryr2/Tiam1 | 2 |
| GO:006007 | CC | GO:006007 | excitatory synapse | 2/61 | 99/28585 | 0.019030636 | 0.019030636 | 0.050851093 | Syt1/Gpm6a | 2 |
| GO:012019 | CC | GO:012019 | cone photoreceptor outer segment | 1/61 | 10/28585 | 0.021139396 | 0.021139396 | 0.053730429 | Slc24a4 | 1 |
| GO:006009 | CC | GO:006009 | kinocilium | 1/61 | 11/28585 | 0.023229002 | 0.023229002 | 0.056295501 | Tiam1 | 1 |
| GO:009773 | CC | GO:009773 | 9+2 non-motile cilium | 1/61 | 11/28585 | 0.023229002 | 0.023229002 | 0.056295501 | Tiam1 | 1 |
| GO:003477 | CC | GO:003477 | secretory granule lumen | 1/61 | 14/28585 | 0.02947153 | 0.02947153 | 0.06824986 | Serpini1 | 1 |
| GO:006020 | CC | GO:006020 | cytoplasmic vesicle lumen | 1/61 | 14/28585 | 0.02947153 | 0.02947153 | 0.06824986 | Serpini1 | 1 |
| GO:003001 | CC | GO:003001 | Z disc | 2/61 | 131/2858 | 0.031988351 | 0.031988351 | 0.072467889 | Ryr2/Cacna1d | 2 |
| GO:003067 | CC | GO:003067 | synaptic vesicle membrane | 2/61 | 143/2858 | 0.037529292 | 0.037529292 | 0.075620557 | Mctp1/Syt1 | 2 |
| GO:009950 | CC | GO:009950 | exocytic vesicle membrane | 2/61 | 143/2858 | 0.037529292 | 0.037529292 | 0.075620557 | Mctp1/Syt1 | 2 |
| GO:001601 | CC | GO:001601 | dystrophin-associated glycoprotein complex | 1/61 | 20/28585 | 0.041839091 | 0.041839091 | 0.079274068 | Sgcz | 1 |
| GO:004429 | CC | GO:004429 | dendrite terminus | 1/61 | 20/28585 | 0.041839091 | 0.041839091 | 0.079274068 | Rgs7 | 1 |
| GO:000172 | CC | GO:000172 | ruffle | 2/61 | 154/2858 | 0.042906652 | 0.042906652 | 0.079845085 | Pacsin1/Tiam1 | 2 |
| GO:000569 | CC | GO:000569 | telomerase holoenzyme complex | 1/61 | 21/28585 | 0.043885225 | 0.043885225 | 0.080233376 | Ptges3 | 1 |
| GO:000024 | CC | GO:000024 | pericentriolar material | 1/61 | 24/28585 | 0.049997875 | 0.049997875 | 0.083186026 | Tiam1 | 1 |
| GO:003301 | CC | GO:003301 | sarcoplasmic reticulum membrane | 1/61 | 24/28585 | 0.049997875 | 0.049997875 | 0.083186026 | Ryr2 | 1 |
| GO:004512 | CC | GO:004512 | pronucleus | 1/61 | 24/28585 | 0.049997875 | 0.049997875 | 0.083186026 | Rad50 | 1 |
| GO:009703 | CC | GO:009703 | perinuclear endoplasmic reticulum | 1/61 | 24/28585 | 0.049997875 | 0.049997875 | 0.083186026 | Kcnd2 | 1 |
| GO:000521 | MF | GO:000521 | monoatomic ion channel activity | 11/59 | 431/2817 | ################ | ################ | ################ | Cacnb4/Slc24a4/Kcnd2/Gabrb2/Ano3/Ryr2/Cacna1d/Gabrg3/H | 11 |
| GO:001526 | MF | GO:001526 | channel activity | 11/59 | 483/2817 | ################ | ################ | ################ | Cacnb4/Slc24a4/Kcnd2/Gabrb2/Ano3/Ryr2/Cacna1d/Gabrg3/H | 11 |
| GO:002280 | MF | GO:002280 | passive transmembrane transporter activity | 11/59 | 483/2817 | ################ | ################ | ################ | Cacnb4/Slc24a4/Kcnd2/Gabrb2/Ano3/Ryr2/Cacna1d/Gabrg3/H | 11 |
| GO:004687 | MF | GO:004687 | metal ion transmembrane transporter activity | 9/59 | 430/2817 | ################ | ################ | ################ | Cacnb4/Slc24a4/Kcnd2/Ryr2/Cacna1d/Slc41a3/Hcn1/Gpm6a/K | 9 |
| GO:002285 | MF | GO:002285 | GABA-gated chloride ion channel activity | 2/59 | 13/28171 | 0.000331391 | 0.000331391 | 0.003292112 | Gabrb2/Gabrg3 | 2 |
| GO:009896 | MF | GO:009896 | postsynaptic neurotransmitter receptor activity | 3/59 | 68/28171 | 0.000396959 | 0.000396959 | 0.003505309 | Gabrb2/Grm1/Gabrg3 | 3 |
| GO:000551 | MF | GO:000551 | calmodulin binding | 4/59 | 192/2817 | 0.000710184 | 0.000710184 | 0.005454639 | Slc24a4/Syt1/Ryr2/Rit2 | 4 |
| GO:000489 | MF | GO:000489 | GABA-A receptor activity | 2/59 | 19/28171 | 0.000720663 | 0.000720663 | 0.005454639 | Gabrb2/Gabrg3 | 2 |
| GO:014029 | MF | GO:014029 | small molecule sensor activity | 2/59 | 33/28171 | 0.002183684 | 0.002183684 | 0.011196477 | Syt1/Rad50 | 2 |
| GO:014082 | MF | GO:014082 | metal cation:monoatomic cation antiporter activity | 2/59 | 36/28171 | 0.00259505 | 0.00259505 | 0.012889885 | Slc24a4/Slc41a3 | 2 |
| GO:004830 | MF | GO:004830 | calcium-dependent protein binding | 2/59 | 76/28171 | 0.011127061 | 0.011127061 | 0.04210993 | Slc24a4/Syt1 | 2 |
| GO:003097 | MF | GO:003097 | receptor tyrosine kinase binding | 2/59 | 79/28171 | 0.01198085 | 0.01198085 | 0.044286617 | Tiam1/Nrg1 | 2 |
| GO:000554 | MF | GO:000554 | phosphatidylinositol-4,5-bisphosphate binding | 2/59 | 94/28171 | 0.016660575 | 0.016660575 | 0.060185332 | Syt1/Hcn1 | 2 |
| GO:001529 | MF | GO:001529 | antiporter activity | 2/59 | 97/28171 | 0.017676129 | 0.017676129 | 0.061324015 | Slc24a4/Slc41a3 | 2 |
| GO:000554 | MF | GO:000554 | phospholipid binding | 4/59 | 480/2817 | 0.018133227 | 0.018133227 | 0.061324015 | Syt1/Pacsin1/Tiam1/Hcn1 | 4 |
| GO:001536 | MF | GO:001536 | calcium:monoatomic cation antiporter activity | 1/59 | 10/28171 | 0.020750521 | 0.020750521 | 0.065897103 | Slc24a4 | 1 |
| GO:001674 | MF | GO:001674 | hydroxymethyl-, formyl- and related transferase activity | 1/59 | 10/28171 | 0.020750521 | 0.020750521 | 0.065897103 | Aldh1l1 | 1 |
| GO:000500 | MF | GO:000500 | transmembrane-ephrin receptor activity | 1/59 | 11/28171 | 0.022802143 | 0.022802143 | 0.065897103 | Epha6 | 1 |
| GO:000823 | MF | GO:000823 | dipeptidyl-peptidase activity | 1/59 | 11/28171 | 0.022802143 | 0.022802143 | 0.065897103 | Dpp10 | 1 |
| GO:009952 | MF | GO:009952 | G protein-coupled neurotransmitter receptor activity | 1/59 | 11/28171 | 0.022802143 | 0.022802143 | 0.065897103 | Grm1 | 1 |
| GO:190168 | MF | GO:190168 | sulfur compound binding | 3/59 | 293/2817 | 0.023599861 | 0.023599861 | 0.066984568 | Ltbp2/Ryr2/Rspo3 | 3 |
| GO:000001 | MF | GO:000001 | single-stranded DNA endodeoxyribonuclease activity | 1/59 | 12/28171 | 0.024849539 | 0.024849539 | 0.069294191 | Rad50 | 1 |
| GO:000486 | MF | GO:000486 | serine-type endopeptidase inhibitor activity | 2/59 | 119/2817 | 0.025886527 | 0.025886527 | 0.069738904 | Itih5/Serpini1 | 2 |
| GO:000216 | MF | GO:000216 | aminoacyl-tRNA editing activity | 1/59 | 13/28171 | 0.026892718 | 0.026892718 | 0.07007421 | Lars2 | 1 |
| GO:004304 | MF | GO:004304 | single-stranded telomeric DNA binding | 1/59 | 13/28171 | 0.026892718 | 0.026892718 | 0.07007421 | Rad50 | 1 |
| GO:000486 | MF | GO:000486 | cAMP-dependent protein kinase inhibitor activity | 1/59 | 14/28171 | 0.028931689 | 0.028931689 | 0.070875572 | Pkib | 1 |
| GO:000411 | MF | GO:000411 | 3',5'-cyclic-AMP phosphodiesterase activity | 1/59 | 15/28171 | 0.030966459 | 0.030966459 | 0.070875572 | Pde3a | 1 |
| GO:000860 | MF | GO:000860 | cAMP-dependent protein kinase regulator activity | 1/59 | 15/28171 | 0.030966459 | 0.030966459 | 0.070875572 | Pkib | 1 |
| GO:003423 | MF | GO:003423 | protein kinase A catalytic subunit binding | 1/59 | 15/28171 | 0.030966459 | 0.030966459 | 0.070875572 | Ryr2 | 1 |
| GO:001664 | MF | GO:001664 | oxidoreductase activity, acting on the CH-NH group of donors | 1/59 | 16/28171 | 0.032997038 | 0.032997038 | 0.070875572 | Aldh1l1 | 1 |
| GO:007074 | MF | GO:007074 | C2H2 zinc finger domain binding | 1/59 | 16/28171 | 0.032997038 | 0.032997038 | 0.070875572 | Ebf1 | 1 |
| GO:014066 | MF | GO:014066 | ATP-dependent DNA damage sensor activity | 1/59 | 16/28171 | 0.032997038 | 0.032997038 | 0.070875572 | Rad50 | 1 |
| GO:000402 | MF | GO:000402 | aldehyde dehydrogenase (NAD+) activity | 1/59 | 17/28171 | 0.035023434 | 0.035023434 | 0.073248456 | Aldh1l1 | 1 |
| GO:000465 | MF | GO:000465 | polypeptide N-acetylgalactosaminyltransferase activity | 1/59 | 18/28171 | 0.037045655 | 0.037045655 | 0.074535562 | Galnt13 | 1 |
| GO:001712 | MF | GO:001712 | phospholipid scramblase activity | 1/59 | 18/28171 | 0.037045655 | 0.037045655 | 0.074535562 | Ano3 | 1 |
| GO:002282 | MF | GO:002282 | solute:potassium antiporter activity | 1/59 | 18/28171 | 0.037045655 | 0.037045655 | 0.074535562 | Slc24a4 | 1 |
| GO:014061 | MF | GO:014061 | DNA damage sensor activity | 1/59 | 19/28171 | 0.03906371 | 0.03906371 | 0.076655234 | Rad50 | 1 |
| GO:000856 | MF | GO:000856 | minus-end-directed microtubule motor activity | 1/59 | 20/28171 | 0.041077608 | 0.041077608 | 0.079624118 | Dnah11 | 1 |
| GO:003021 | MF | GO:003021 | semaphorin receptor binding | 1/59 | 22/28171 | 0.045092963 | 0.045092963 | 0.083341952 | Rit2 | 1 |
| GO:003055 | MF | GO:003055 | cAMP binding | 1/59 | 22/28171 | 0.045092963 | 0.045092963 | 0.083341952 | Hcn1 | 1 |
| GO:003459 | MF | GO:003459 | phosphatidylinositol bisphosphate phosphatase activity | 1/59 | 22/28171 | 0.045092963 | 0.045092963 | 0.083341952 | Inpp4b | 1 |
| GO:007018 | MF | GO:007018 | DNA polymerase binding | 1/59 | 22/28171 | 0.045092963 | 0.045092963 | 0.083341952 | Ptges3 | 1 |
| GO:003050 | MF | GO:003050 | ankyrin binding | 1/59 | 23/28171 | 0.047094438 | 0.047094438 | 0.08455908 | Cacna1d | 1 |
| GO:000485 | MF | GO:000485 | enzyme inhibitor activity | 3/59 | 392/2817 | 0.049074817 | 0.049074817 | 0.08455908 | Itih5/Pkib/Serpini1 | 3 |
